# A catalytic mechanism for *Renilla*-type bioluminescence

**DOI:** 10.1101/2022.02.09.479090

**Authors:** Andrea Schenkmayerova, Martin Toul, Daniel Pluskal, Racha Baatallah, Glwadys Gagnot, Gaspar P. Pinto, Vinicius T. Santana, Marketa Stuchla, Petr Neugebauer, Pimchai Chaiyen, Jiri Damborsky, David Bednar, Yves L. Janin, Zbynek Prokop, Martin Marek

## Abstract

The widely used coelenterazine-powered *Renilla* luciferase was discovered over 40 years ago but the oxidative mechanism by which it generates blue photons remains unclear. Here we decipher *Renilla*-type bioluminescence through crystallographic, spectroscopic, and computational experiments. Structures of ancestral and extant luciferases complexed with the substrate-like analogue azacoelenterazine or a reaction product were obtained, providing unprecedented snapshots of coelenterazine-to-coelenteramide oxidation. Bound coelenterazine adopts a Y-shaped conformation, enabling the deprotonated imidazopyrazinone component to attack O_2_ via a radical charge-transfer mechanism. A high emission intensity is secured by an aspartate from a conserved proton-relay system, which protonates the excited coelenteramide product. Another aspartate on the rim of the catalytic pocket fine-tunes the electronic state of coelenteramide and promotes the formation of the blue light-emitting phenolate anion. The results obtained also reveal structural features distinguishing flash-type from glow-type bioluminescence, providing insights that will guide the engineering of next-generation luciferase–luciferin pairs for ultrasensitive optical bioassays.

## Introduction

Bioluminescence is a fascinating phenomenon involving the emission of light by a living creature. There is enormous interest in harnessing bioluminescent systems to design ultrasensitive optical bioassays and enable a circular bio-economy.^1–4^ Bioluminescent organisms generate light via the oxidation of a substrate (a luciferin), which is catalyzed by a class of enzymes called luciferases.^5^

One of the most popular bioluminescent reporters is a luciferase isolated from the sea pansy *Renilla reniformis,* a soft coral that displays bioluminescence upon mechanical stimulus.^6^ *Renilla* luciferase, henceforth referred to as RLuc, is a 36 kDa protein that is active as a monomer.^7,8^ RLuc displays remarkable sequence and structural similarity to a family of haloalkane dehalogenases (HLDs), indicating a common evolutionary history.^9–11^ Unlike HLDs, which belong to the α/β-hydrolase family (EC 3.8.1.5)^12^, the RLuc luciferase is an ATP-independent monooxygenase (EC 1.13.12.5)^7,8^ that catalyzes the conversion of coelenterazine (CTZ) to coelenteramide (CEI). The blue light emission of *Renilla* luciferase has fascinated scientists for decades. However, despite intensive efforts^9,13–15^, a detailed understanding of its catalytic mechanism at the molecular level remains elusive.

CTZ, or 2-(*p*-hydroxybenzyl)-6-(*p*-hydroxyphenyl)-8-benzylimidazo [1,2-a]pyrazine-3-(7H)-one, is the most common marine luciferin. It features an imidazo[1,2-a]pyrazine ring system that emits a photon after undergoing an O_2_-mediated oxidation whose other products are CEI and CO_2_.^16–19^ Importantly, CTZ can spontaneously emit a photon as a result of autooxidation; this process is known as chemiluminescence and is favored in aprotic solvents such as dimethyl sulfoxide (DMSO).^20,21^ As shown in **Fig. 1a**, mechanisms for CTZ oxidation have been suggested by McCarpa^16^ and Goto^17^, and more recently updated by other authors.^22,23^ It was originally assumed that the reaction starts via deprotonation of the N7-nitrogen to form a CTZ anion that reacts directly with O_2_ at the C2 carbon to form a CTZ peroxide ion. However, no known photoproteins or luciferases have an amino acid in the vicinity of the N7-nitrogen of the bound CTZ that could potentially mediate this initial deprotonation. Griffiths and coworkers therefore proposed that CTZ may bind to a protein as its O10H tautomer, or as an already deprotonated species (**Fig. 1a**).^23^ Following the oxygen addition, the resulting nucleophilic peroxide moiety performs an intramolecular attack on the C3 carbonyl to form a cyclic dioxetanone intermediate. Subsequent ring decomposition via decarboxylation leads to loss of CO_2_ and the formation of an aminopyrazine product (the CEI ion) in a singlet excited state. While this is the light-emitting species in chemiluminescence, bioluminescent systems probably protonate the N1^-^ amide ion to form a light-emitting neutral CEI species. Moreover, some photoproteins and also possibly some luciferases remove another proton from the 6-(*p*-hydroxyphenyl) substituent to form a light-emitting 6-*p*-hydroxyphenolate ion. Finally, it has been demonstrated that some bioluminescent systems do not produce CEI as the major product of CTZ oxidation but instead convert CTZ into coelenteramine (CNM) and (*p*-hydroxyphenyl)acetic acid.^22^ It is not currently clear whether CTZ oxidation proceeds via a single electron transfer process involving radical intermediates, as seen in the oxidation of firefly luciferin by the luciferase of *Photinus pyralis*^24^.

**Fig. 1.**
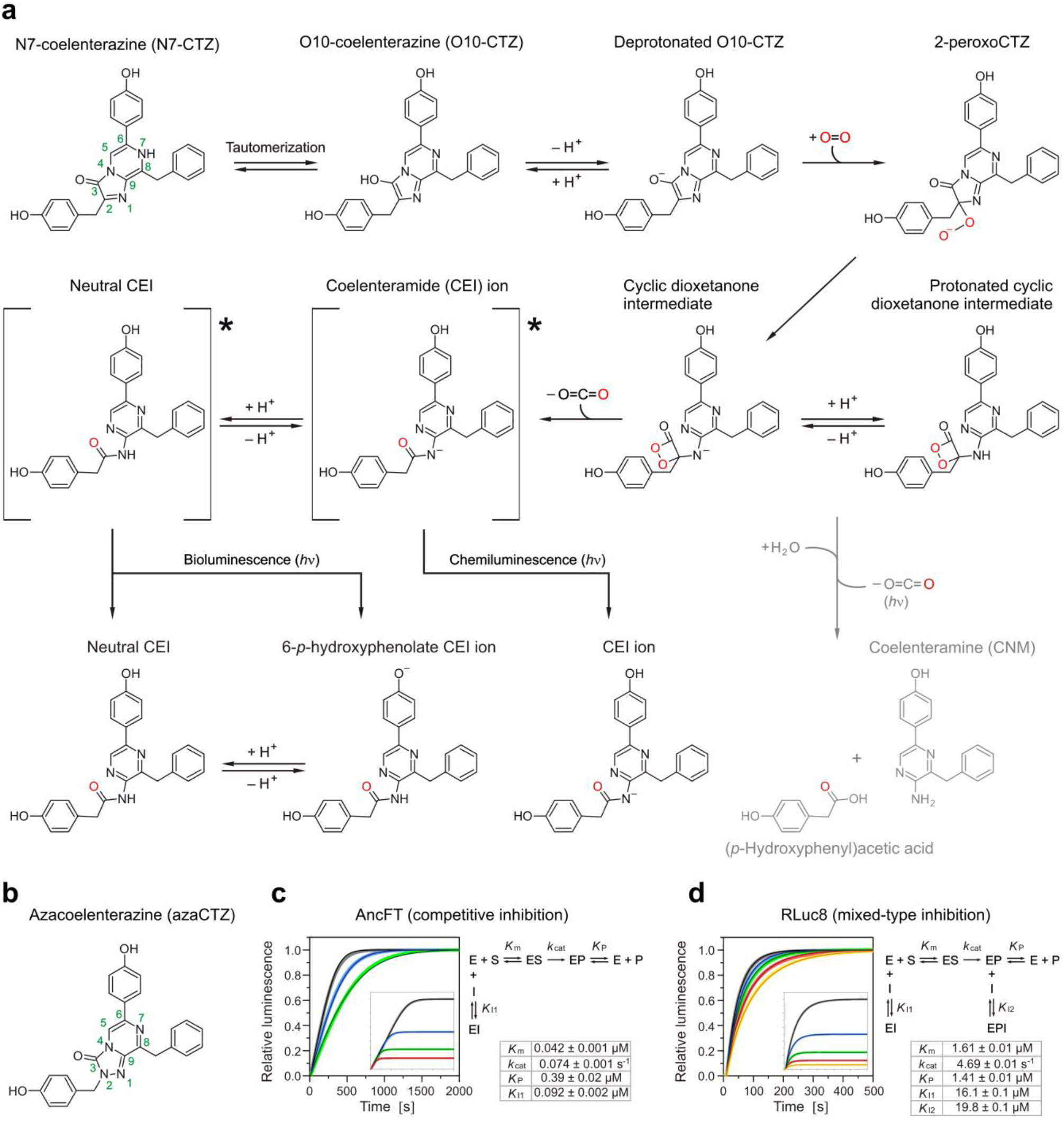
Inhibitory effects of azacoelenterazine (azaCTZ) on AncFT and RLuc8 bioluminescence. (**a**) General bioluminescent and chemiluminescent reaction mechanisms of coelenterazine (CTZ) conversion. CTZ exists in two tautomeric forms, N7-CTZ and O10-CTZ. The reaction is initiated by the dissociation of a proton from the O10-oxygen to produce anionic O10-CTZ, whose C2 carbon then attacks O_2_ to form the 2-peroxoCTZ ion. The distal oxygen of 2-peroxoCTZ performs an intramolecular nucleophilic attack on the C3-carbon to form a cyclic dioxetanone intermediate whose subsequent decarboxylative decomposition releases CO_2_ and forms an anionic excited-state aminopyrazine product: the coelenteramide (CEI) ion. This unstable anionic species then decomposes with the emission of light, resulting in chemiluminescence. Bioluminescent systems protonate the CEI anion at the amide position to form the corresponding neutral species. Light can be emitted from this neutral species, but more often another proton is removed to form the 6-*p*-hydroxyphenolate ion, which emits blue light with an emission maximum at ~480 nm. Some bioluminescent systems do not produce CEI as the major product of CTZ oxidation but instead convert CTZ into coelenteramine (CNM) and (*p*-hydroxyphenyl)acetic acid, which also results in light emission. (**b**) Structure of azaCTZ. (**c**) Reaction progress curves obtained upon mixing 0.013 μM AncFT with 0.275 μM CTZ in the presence of 0 μM (black), 0.3 μM (blue) and 0.5 μM (green) azaCTZ, showing that azaCTZ is a competitive inhibitor of AncFT bioluminescence. Inset: kinetic data recorded upon mixing 0.02 μM AncFT with 0.138 μM (black), 0.275 μM (blue), 0.55 μM (green) and 1.1 μM (red) CTZ. (**d**) Reaction progress curves recorded upon mixing 0.02 μM Rluc8 with 2 μM coelenterazine in the presence of 0 μM (red), 4.5 μM (green), 9 μM (blue) 18 μM (yellow) and 30 μM (cyan) azaCTZ, showing that azaCTZ is a mixed-type inhibitor of RLuc8 bioluminescence. Inset: kinetic data recorded upon mixing 0.02 μM Rluc8 with 0.25 μM (cyan), 0.5 μM (yellow), 1 μM (blue), 2 μM (green) and 4 μM CTZ (red). Each trace represents an average of three replicates. Solid lines represent the best global fit of the kinetic data. Relative luminescence values are plotted. Steady-state kinetic parameters were obtained by global fitting of kinetic data using numerical integration of rate equations and are reported as values ± standard errors (SE) based on the global fit.

A key barrier to a deeper understanding of *Renilla*-type catalysis is the limited availability of structural data on substrate- and product-bound enzyme complexes. Crystallization of native RLuc is challenging, so mutagenesis experiments were performed to obtain mutants more amenable to crystallization.^25,26^ As a result, a stabilized eight-point mutant designated RLuc8^26^ was crystallized by Loening and coworkers.^14^ A partial density for CEI bound outside of the active site was observed in one of their structures, although this structure is unlikely to be biologically relevant. In addition, we recently co-crystallized multiple RLuc8 mutants in the presence of excess CTZ. These efforts yielded a CEI-bound RLuc8 structure, providing the first biologically-relevant insights into a post-catalytic complex.^13^ Unfortunately, the lack of a stable non-oxidizable CTZ analogue and the intrinsic flexibility of extant RLuc mutants seriously complicate the acquisition of high-resolution structures of catalytically-favored Michaelis enzyme-substrate complexes.

High protein stability is known to facilitate crystallization^27–29^, and it has been demonstrated that the structures of ancestral enzymes can be used as molecular scaffolds to unravel the structures and catalytic mechanisms of extant enzymes that are difficult to crystallize.^30–35^ We previously used ancestral sequence reconstruction to obtain a stable ancestral enzyme Anc^HLD-RLuc^ that was reconstructed from the catalytically distinct but structurally related HLDs and *Renilla* luciferase.^9^ This ancestor enzyme turned out to have dual functions, with dehalogenase activity comparable to that of contemporary HLDs as well as promiscuous luciferase activity significantly lower than that of the stabilized RLuc8. We recently subjected Anc^HLD-RLuc^ to insertion-deletion backbone mutagenesis and thereby discovered key structural elements responsible for acquisition of luciferase activity. In particular, transplanting a loop-helix fragment from the extant RLuc luciferase into the Anc^HLD-RLuc^ ancestor yielded a fragment-transplanted AncFT enzyme with 7,000-fold higher catalytic efficiency and 100-fold longer lasting glow-type bioluminescence.^13^ AncFT displays higher stability, enhanced substrate affinity, and lower product inhibition than the “parent” ancestral enzyme^13^, and is thus a perfect surrogate system for dissecting the bioluminescence of α/β-hydrolase fold luciferases.

Here we present an atomistic description of the catalytic mechanism of *Renilla*-type luciferases that was inferred from co-crystal structures of AncFT and RLuc8 luciferases complexed with either a newly synthesized non-oxidizable CTZ analogue (azacoelenterazine) or a catalytic product. These complexes reveal key structural features of the catalytic machinery responsible for CTZ conversion. Moreover, we delineate the biophysical factors responsible for flash-type and glow-type bioluminescence, and show how the electronic state of the CEI ion is tuned to favor blue light emission. The proposed mechanism is supported by data from spectroscopic, mutagenesis, and computational experiments. Collectively, our findings provide mechanistic understanding of *Renilla* bioluminescence and will facilitate the engineering of next-generation luciferase-luciferin bioluminescent pairs.

## Results

### Azacoelenterazine acts as a non-oxidizable substrate analogue

To probe the mechanism of CTZ monooxygenation by *Renilla*-type luciferases, we designed and synthesized the non-oxidizable CTZ analogue 8-benzyl-2-(4-hydroxybenzyl)-6-(4-hydroxyphenyl)- [1,2,4]triazolo[4,3-a]pyrazin-3(2H)-one, which is henceforth referred to as azacoelenterazine (azaCTZ). The C2 core carbon atom of CTZ, which interacts with O_2_ during the monooxygenation reaction, is replaced with a nitrogen atom in azaCTZ, to prevent dioxygen attack (**Fig. 1b**). The synthesis of azaCTZ was achieved in 19% overall yield in four steps from a chloropyrazine intermediate (**Supplementary Fig. 1**).^36^

Having synthesized azaCTZ, we studied its *in vitro* effects on the bioluminescent reactions of AncFT and RLuc8 by performing steady-state kinetics experiments. These experiments confirmed that azaCTZ is a non-oxidizable CTZ analogue that mimics the binding of CTZ. A detailed description of the kinetic analysis is provided in **Supplementary Notes 1** and **2**. Numerical integration of the luminescence progress curves obtained for the two enzymes in the presence of mixtures of CTZ and azaCTZ provided precise estimates of their steady-state kinetic parameters that agreed with previously reported values (**Fig. 1c,d**).^13^ Global fittings based on kinetic data recorded in the absence and presence of different concentrations of azaCTZ were systematically analyzed to clarify the mechanisms by which the inhibitor interacted with RLuc8 and AncFT. In the case of AncFT, a model in which the inhibitor competes with the substrate for a single binding site gave the best agreement with the kinetic data (**Fig. 1c** and **Supplementary Figs. 4-6**). Additionally, the dissociation constant for the enzyme-inhibitor complex (*K*_I_ = 0.092 ± 0.002 μM) calculated using the competitive model is similar to the Michaelis constant of AncFT, further supporting the assumption that azaCTZ binds to the active site of this enzyme in a very similar manner to the native substrate. AzaCTZ is thus a pure competitive inhibitor of bioluminescent CTZ degradation catalyzed by AncFT (**Supplementary Fig. 2**).

A more complex mixed-type inhibition model gave the best fit to the kinetic data acquired for the reaction of RLuc8 (**Fig. 1d** and **Supplementary Figs. 6-8**). The full time-course kinetics of the substrate-to-product conversion provided additional estimates of the equilibrium dissociation constants for the enzyme-product complex, making it possible to elucidate the more complex inhibitory mechanism in this case. The proposed model includes competitive binding of azaCTZ to the active site of RLuc8 (in keeping with the model selected for AncFT) but also suggests that the inhibitor can bind to an enzyme-product complex. The fact that RLuc8 can simultaneously bind to the substrate and the inhibitor but does so with affinities that are an order of magnitude lower than those of AncFT (*K*_m_ = 1.66 μM and *K*_I1_ = 16.1 μM for RLuc8, as compared to *K*_m_ = 0.042 μM and *K*_I1_ = 0.092 μM for AncFT) can be explained by the greater conformational flexibility of RLuc8. These kinetic data agree with our anaerobic equilibrium binding experiments, which also showed that the affinity of AncFT for CTZ and CEI greatly exceeded that of RLuc8 (**Supplementary Fig. 9**). Collectively, these results suggest that azaCTZ could be a valuable tool for probing pre-catalytic enzyme-substrate complexes through X-ray crystallography.

### Azacoelenterazine- and coelenteramide-bound AncFT structures

To obtain structural insights into the RLuc catalytic mechanism, we attempted co-crystallization of AncFT with excesses of azaCTZ or its native ligand CTZ. It was expected that the latter ligand would be converted by the enzyme into the catalytic product CEI. After screening co-crystallization conditions and optimizing hits, well-diffracting crystals were obtained. Their structures were then solved by molecular replacement and the initial models obtained were further refined through several cycles of manual building and automatic refinement, yielding structural models with low deviations from ideal geometry (**Supplementary Table 3**). These AncFT structures show a canonical αβα-sandwich architecture and closely resemble the previously reported structure of *apo* AncFT^13^, with root-mean-square deviation (RMSD) values on Cα atoms ranging from 0.1 to 0.3 Å (**Supplementary Fig. 10**).

Inspections of the electron density maps unambiguously revealed azaCTZ or CEI molecules bound in the active site pocket of the corresponding co-crystal structures (**Fig. 2** and **Supplementary Fig. 11**). Both azaCTZ and CEI adopt a Y-shaped conformation when bound to the enzyme pocket. The R^1^ 2-(*para*-hydroxybenzyl) substituent is deeply buried in the active site cavity, where it is anchored in the slot p2 tunnel via multiple polar and nonpolar interactions. In addition, its terminal hydroxyl moiety forms a hydrogen bond with the backbone carbonyl of S143 (~2.8 Å). The triazolopyrazine core of azaCTZ is positioned in close proximity to the conserved catalytic machinery, which consists of a catalytic pentad that was first characterized in the structurally related HLDs^12,37^. The triazolopyrazine core of azaCTZ forms direct contacts with the carboxyl side chains of D118 (~3.3 Å) and W119 (~3.3 Å). The former residue is a nucleophilic aspartate, while the latter tryptophan functions as a halide-stabilizing residue in the HLD reaction^12,37^. The remaining two substituents, namely the R^2^ *6-(para-* hydroxyphenyl) and R^3^ 8-benzyl, occupy the main p1 tunnel and form multiple interactions, most of which are hydrophobic or aromatic in nature. The R^3^ 8-benzyl group forms π-π stacking interactions with F260 (~3.7 Å) and F259 (~4.9 Å), while the hydroxyl group of the R^2^ 6-(*para*-hydroxyphenyl) substituent forms a hydrogen bond with the carboxyl side chain of D160, an aspartate residue located at the rim of the catalytic cavity (helix α4).

**Fig. 2.**
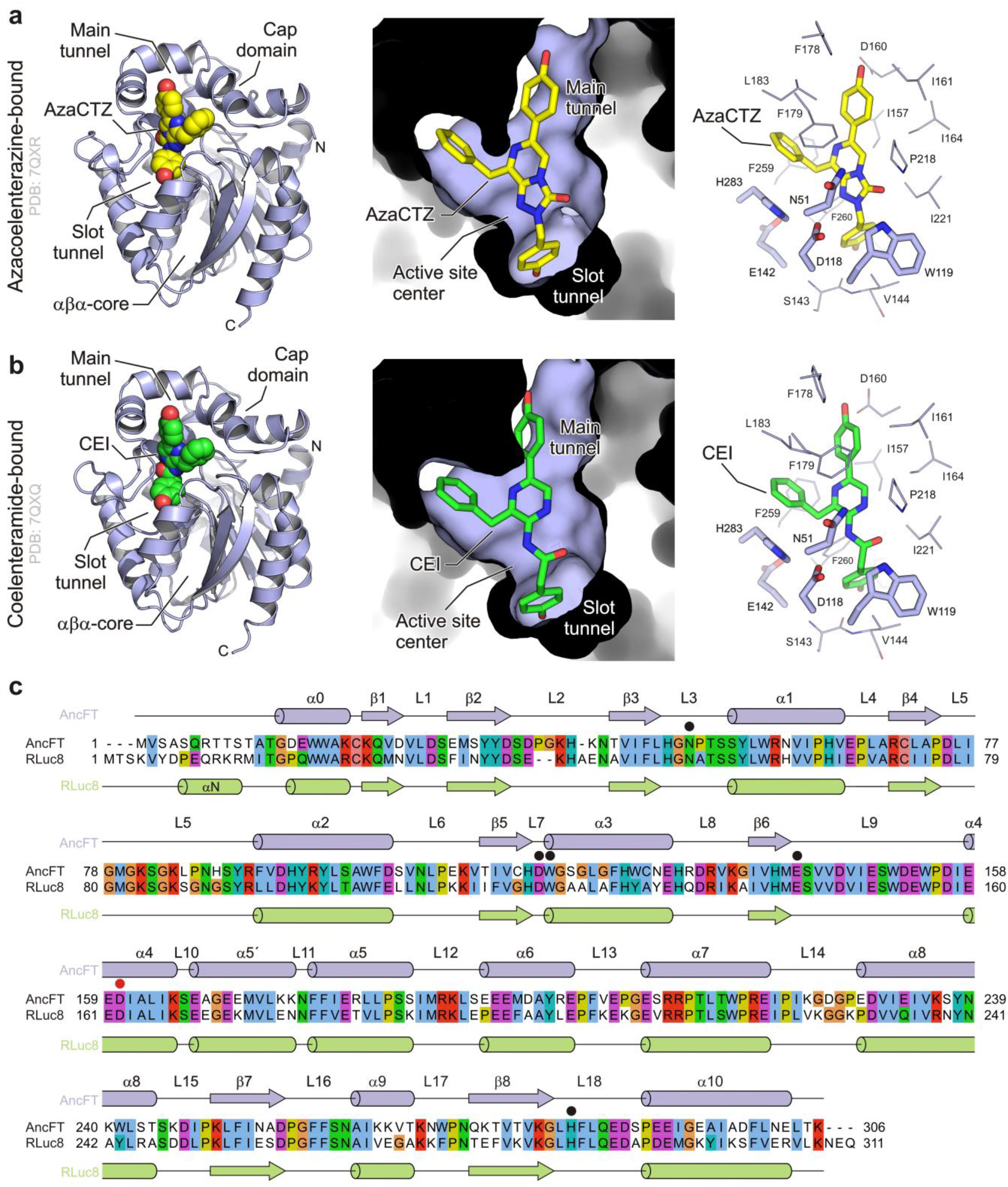
Structures of azacoelenterazine- and coelenteramide-bound AncFT luciferase. (**a**) Binding mode of azacoelenterazine (azaCTZ) to AncFT. (**b**) Binding mode of coelenteramide (CEI) to AncFT. Left panels: cartoon representations of the overall structures of the azaCTZ (yellow) and CEI (green) complexes, shown as space-filling calotte models. Middle panels: cutaway surface representations of the active site cavity with bound azaCTZ and CEI shown as sticks. Right panels: close-up views of the binding modes of azaCTZ and CEI in the AncFT active-site pocket. Residues of the conserved catalytic pentad are shown as light blue sticks, the remaining protein residues are shown as light blue lines, azaCTZ is shown as yellow sticks and the CEI is shown as green sticks. (**c**) Structure-based sequence alignment of AncFT and RLuc8. Secondary structure elements found in AncFT and RLuc8 are shown above and below the alignment, respectively. Catalytic pentad residues are labeled with black dots and the aspartate at the rim of the catalytic pocket is labeled with a red dot.

A similar binding mode is observed in the CEI-bound AncFT complex structure (**Fig. 2b**). There are no major structural differences between the protein backbones of the azaCTZ- and CEI-bound AncFT structures (RMSD ~0.2 Å). In both cases, active-site pocket residues wrap tightly around the bound ligand, preventing both its random motion and free access of solvent molecules. Numerous protein residues including F178, L183, S187, F284, H283, E142, V144, W154, I221 and P218 are involved in this first shell surrounding the bound ligand. Together, these co-crystal structures provide unprecedented snapshots and molecular details of enzyme-substrate and enzyme-product complexes in catalytically-favored states.

### The malleability of the RLuc8 active-site pocket

Structural characterization of RLuc luciferase and its mutants (e.g. RLuc8) has proven to be challenging and previous studies provided little understanding of its catalytic mechanism.^13,14^ We therefore performed a new round of co-crystallizations with RLuc8 mutants in the presence of molar excesses of either azaCTZ or CTZ in order to obtain molecular insights into its catalysis. The co-crystallization experiments using the RLuc8-D162A and RLuc8-D120A mutants yielded well-diffracting crystals, and the corresponding structures could be solved by molecular replacement. Crystallographic and refinement statistics for the resulting complexes are presented in **Supplementary Table 4**. As for the AncFT proteins, interpretation of the electron density maps of the RLuc8-D162A complexes unambiguously identified azaCTZ or CEI. Unexpectedly, we also observed that coelenteramine (CNM), another product of CTZ oxidation^22^, was bound to RLuc8-D120A (**Fig. 3c** and **Supplementary Fig. 12**). In every instance, these ligands were located inside the catalytic pocket.

**Fig. 3.**
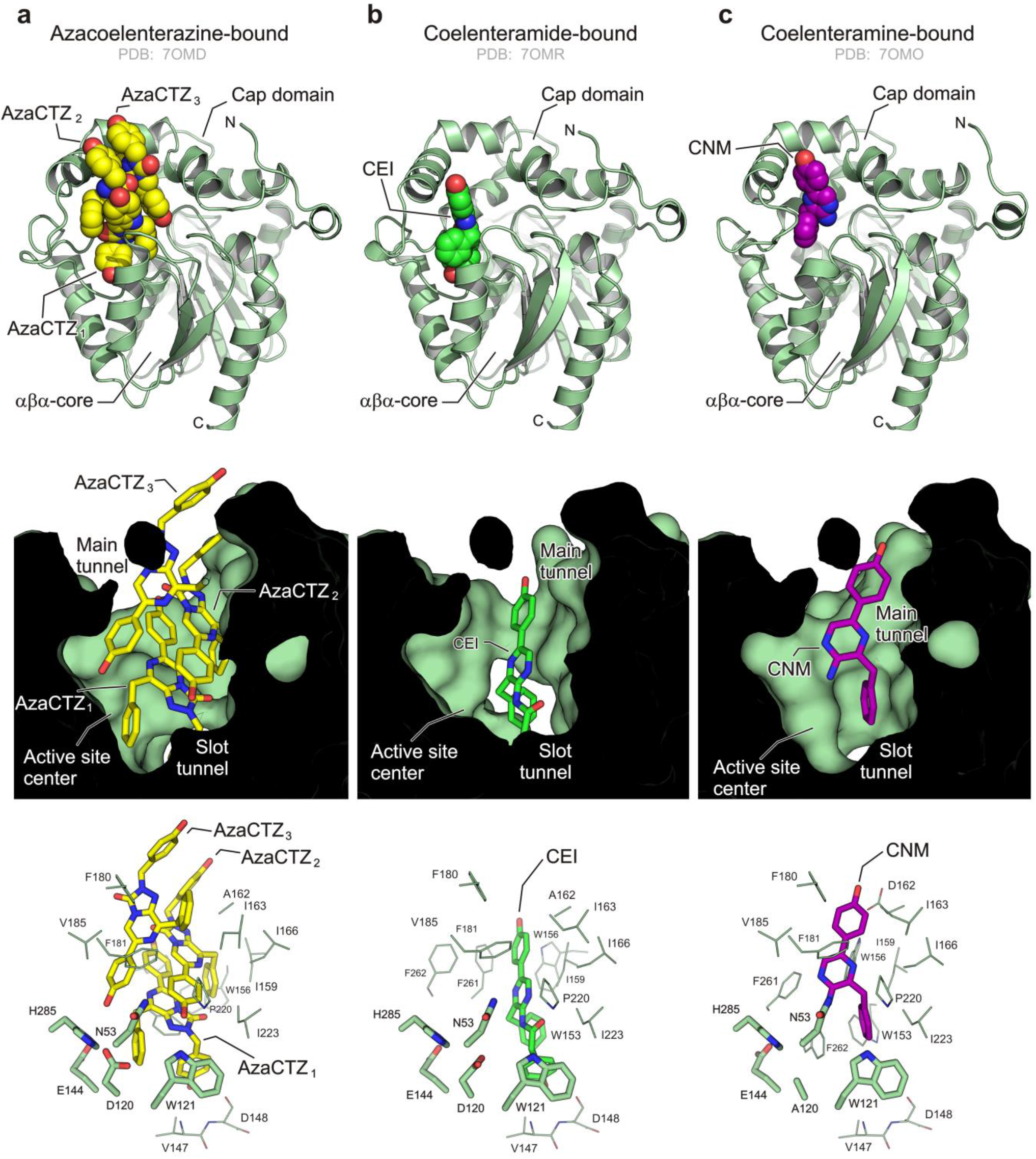
Structures of azacoelenterazine-, coelenteramide-, and coelenteramine-bound RLuc8 variants. Snapshots of RLuc8-D162A/azaCTZ (**a**), RLuc8-D162A/CEI (**b**) and RLuc8-D120A/CNM (**c**) complexes. Top panels: cartoon representations of the overall structures of azaCTZ (yellow), CEI (green), and CNM (magenta) complexes, shown as space-filling calotte models. Middle panels: cutaway surface representations of the active site cavity with bound ligands shown as sticks. Bottom panels: close-up views of the binding modes of azaCTZ, CEI and CNM in the RLuc8 active-site pocket. Residues of the conserved catalytic pentad are shown as light green sticks, the remaining protein residues are shown as pale green lines, azaCTZ is shown as yellow sticks, CEI is shown as green sticks, and CNM is shown as magenta sticks.

A common feature of all the newly determined RLuc8 complex structures is a voluminous active-site cavity (**Fig. 3**). This is most pronounced in the RLuc8-D162A/azaCTZ complex, where we found as many as three azaCTZ molecules bound in the same active-site pocket. One azaCTZ molecule is deeply buried in the cavity with an orientation resembling that seen in the AncFT/azaCTZ complex (**Fig. 2a**), while the remaining two azaCTZ molecules are rotated through ~180° and occupying the remaining volume of the main p1 tunnel (**Fig. 3a**). A similarly open catalytic pocket is seen in the RLuc8-D162A/CEI and RLuc8-D162A/CNM complexes in which only a single ligand (CEI or CNM, respectively) is bound (**Fig. 3b,c**). These new ligand-bound structures show that RLuc8 explores a very large conformational space and provide a structural basis for the accommodation of two or more ligands in its catalytic pocket, in keeping with the results of our kinetics experiments (**Fig. 1d**).

To demonstrate this extreme malleability, we generated a gallery of RLuc8 crystallographic snapshots illustrating active site cavity opening states ranging from minimally to maximally open (**Fig. 4a**). The main structural elements responsible for this malleability are the α4 helix, the L9 loop, and the L16 loop. Conformational sampling of these structural elements dramatically changes the catalytic pocket’s volume and shape. Bulky aromatic residues in the L9 loop (W153 and W156) and the L16 loop (F261 and F262) appear to play particularly important roles in determining the openness of the active site. Although this conformational flexibility may be important in some steps of the catalytic cycle, it greatly complicated our attempts to obtain a crystallographic structure of a biologically relevant Michaelis complex.

**Fig. 4.**
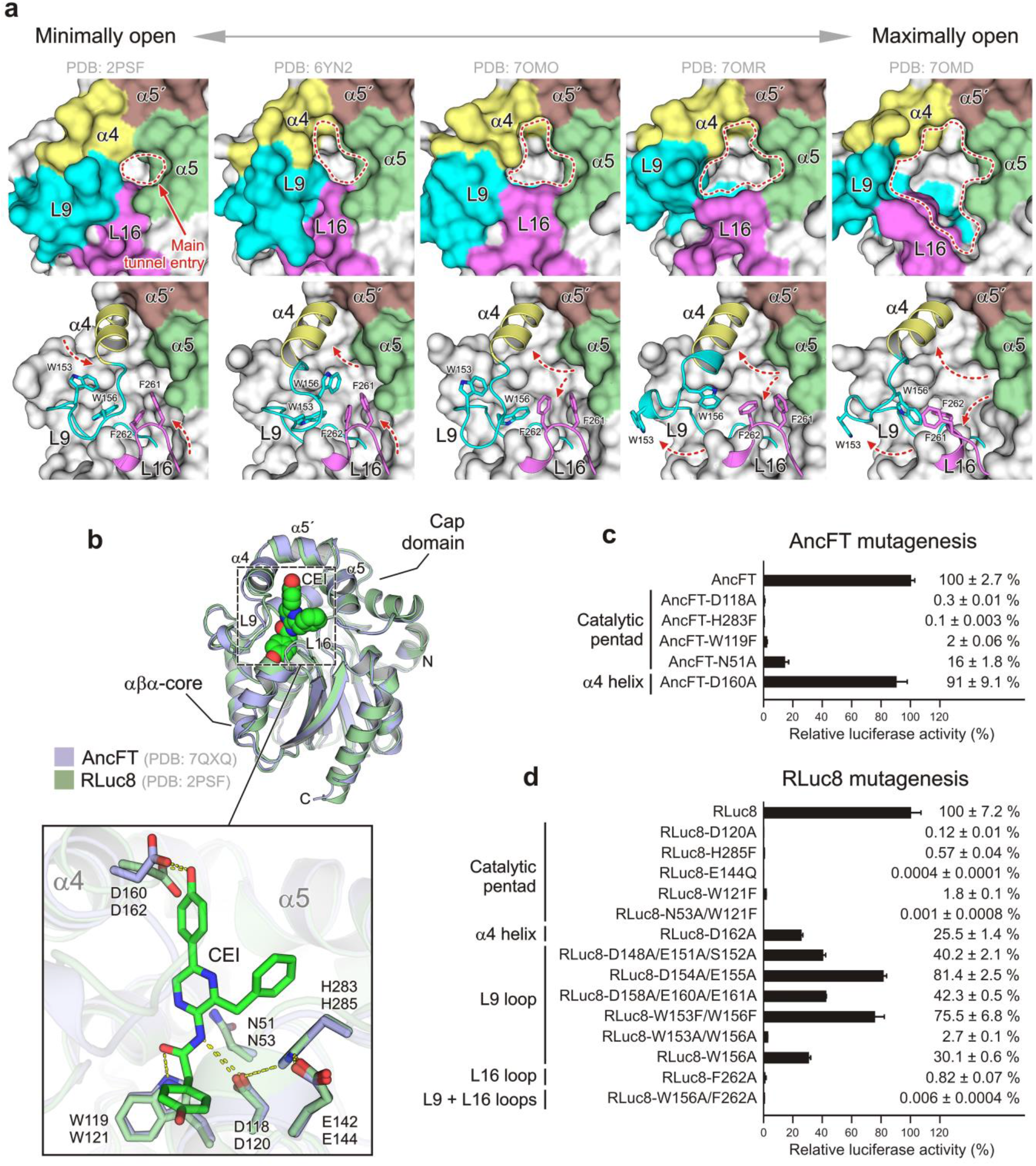
Structural and functional determinants of *Renilla*-type bioluminescence. (**a**) A gallery of RLuc8 structures revealing the malleability of the RLuc8 active site cavity. Top panels: surface representations of the main tunnel entry to the catalytic pocket. The pocket walls are formed by residues from the α4 (yellow), α5’ (brown) and α5 (green) helices and the L9 (cyan) and L16 (violet) loops. Bottom panels: a combination of surface and cartoon representations highlighting the protein backbone motions that regulate the volume of the active-site cavity. Residues W153 and W156 from the L9 loop and F261 and F262 from the L16 loop are shown as sticks. Note that the reorientation of the α4 helix, coupled with conformational changes of loops L9 and L16, regulates the opening and closing of the main access tunnel. (**b**) Structural comparison of AncFT and RLuc8. Top panel: Superposition of the CEI-bound AncFT complex (light blue; PDB: 6TOX) and a CEI-RLuc8 complex showing minimal opening of the active site pocket (pale green; PDB: 2PSF). In both cases, the CEI molecule is shown as a green space-filling calotte model. Bottom inset: Close-up view of CEI in the active site cavities of the superposed structures. (**c,d**) Mutagenesis of AncFT (**c**) and RLuc8 (**d**). Data indicate the average relative luciferase activities of each mutant. Assays were done in triplicate; error bars represent standard deviations.

### A basis for distinguishing flash-type and glow-type bioluminescence

We previously showed that although RLuc8 displays high catalytic turnover (*k*_cat_ = ~4.7 s^-1^), it has a relatively low affinity for its substrate (*K*_m_ = ~1.5 μM) and exhibits significant product inhibition (*K*_p_ = ~1.2 μM). This may explain why the bioluminescent signal decays rapidly after a strong initial flash.^13^ On the other hand, compared to RLuc8, AncFT displayed >20-fold higher affinity toward the substrate (*K*_m_ = ~0.064 μM), exhibited distinctly weaker product inhibition (*K*_p_ = ~0.5 μM), and generated a markedly stable glow-type bioluminescent signal.^13^ The co-crystal structures of the two enzymes reported here are consistent with these findings. The RLuc8 catalytic pocket is voluminous and malleable (**Figs. 3** and **4a**), which allows simultaneous accommodation of two or more ligand molecules and explains its tendency toward product inhibition and flash-type bioluminescence. Conversely, the AncFT catalytic pocket is less malleable and can only accommodate a single ligand molecule (**Fig. 2**), which explains its higher affinity for CTZ, lack of product inhibition, and potentially highly stable glow-type bioluminescence. The structural and kinetic data for RLuc8 highlight the importance of our engineered AncFT protein, which is intrinsically less flexible than RLuc8 and is thus a valuable surrogate for studying key chemical steps of the luciferase reaction.

### Catalytic analogy between AncFT and RLuc8

Next, we aimed to demonstrate that AncFT and RLuc8 are functionally analogous in terms of their catalytic chemistry. We therefore performed pairwise protein backbone comparisons, revealing that the greatest similarity existed between AncFT and the *apo*-form of RLuc8 with a minimally open active site cavity. Superposition of these two structures revealed that their backbones have very similar geometries (**Fig. 4b**). Importantly, the positioning of the side chains of the putative catalytic pentad residues is identical, allowing analogous chemistry. Moreover, the D160 aspartate residue at the rim of the cap domain in AncFT, which forms a hydrogen bond with the R^2^ 6-(*p*-hydroxyphenyl) substituent of the substrate, overlaps well with the D162 residue in RLuc8. Structural comparisons thus showed that AncFT and RLuc8 have very similar active site chemistries.

To experimentally validate the functional roles of the residues highlighted in our co-crystal structures, we performed structure-based mutagenesis of AncFT and RLuc8. All of the generated mutants were recombinantly prepared as soluble proteins. As shown in **Fig. 4c**, mutating the conserved catalytic pentad residues in AncFT severely affected its luciferase activity. The most compromising effects were observed for the AncFT-D118A, AncFT-W119F and AncFT-H283F mutants, which retained only ~0.3%, ~2% and ~0.1% of the activity of AncFT (100%). A less severe inactivating effect was observed for AncFT-N51A (~16%). Interestingly, an aspartate-to-alanine mutation in the cap α4 helix (AncFT-D160A) had only a minimal effect on AncFT catalysis (~91%).

Extensive mutagenesis experiments with RLuc8 have been reported previously^14,15,26,38,39^. We repeated some of these experiments in this work and also performed some new ones. Crucially, we found that the catalytic pentad residues are also critical for RLuc8 activity (**Fig. 4d**). Mutations in the active-site loops L9 and L16 showed that residues with bulky aromatic side chains (W153, W156 and F262) are catalytically important; this is consistent with the structural data showing that these residues make aromatic π-π contacts with CEI. Accordingly, the double-point mutation RLuc8-W156A/F262A caused a > 16,600-fold reduction in luciferase activity (**Fig. 4d**), demonstrating the importance of these bulky aromatic residues. In contrast, mutagenesis of polar and charged residues in the L9 loop showed that they are not critical for the luciferase reaction (**Fig. 4d**). Collectively, our mutagenesis experiments showed that the residues of the conserved catalytic pentad in AncFT and RLuc8 are functionally essential, confirming the catalytic analogy between these two enzymes. This conclusion is also justified by molecular dynamics simulations of key protein-ligand complexes of both AncFT and RLuc8, as discussed below.

### Computational modeling of enzyme-ligand complexes

We performed triplicate 10 ns long MD simulations of the putative mechanistic ground states to determine whether the reaction mechanism inferred for AncFT can be extrapolated to RLuc8. To this end, we selected the most energetically favorable conformations for each putative chemical step and compared their binding geometries in the two enzymes. Because we had previously established that N53 and W121 (N51 and W119 in AncFT) stabilize O_2_ and the leaving CO_2_, we focused on the positions of the other residues involved in the catalytic mechanism (**Fig. 4b**).

The residues D120 (AncFT D118), E144 (AncFT E142), and H285 (AncFT H283) facilitate and stabilize proton transfer from the enzyme to the luciferin. In the E.S. complexes, the histidine residues of these triads form stabilizing hydrogen bonds with the aspartate and glutamate residues: the H285 to D120 and H285 to E144 distances in RLuc8 are 2.4 Å and 2.7 Å, respectively, while the corresponding distances in AncFT (H283 to D118 and H283 to E144) are 2.5 Å and 2.2 Å, respectively. At this point the distance from N1 of the substrate to the aspartate is below 3.5 Å. In the E.2-peroxy-CTZ complexes, dioxygen is already bound to the substrate, the H-bond distances are <3 Å, and the distance from the substrate N1 nitrogen to the catalytic aspartate is 3.3 Å in RLuc and 3.1 Å in AncFT. These distances are optimal for proton transfer from the enzyme’s histidine to the substrate, via the aspartate. In the E.dioxetanone complexes, the proton has been transferred to the substrate and the distance from the substrate to the aspartate is slightly increased. However, the stabilizing hydrogen bond between the histidine and the glutamate remains; these two residues show little fluctuation during the putative chemical steps. Longer distances between the product, the aspartate, and the histidine are seen in the E.CEI complexes. As shown in **Supplementary Fig. 13**, the distances between these residues and from the catalytic residues to the substrate are higher on average in the RLuc8 complexes. However, the orientation of the ligands and catalytic residues in both enzymes are very similar and are stable in all steps, suggesting that these enzymes employ analogous catalytic mechanisms. Furthermore, the simulations confirmed that the conformational dynamics of the L9 and L16 loops are unusually pronounced, in accordance with the crystallographic structures of the protein-ligand complexes.

### Coelenterazine enters the enzymatic pocket with a deprotonated core

We also determined the absorbance peak maxima of CTZ, CTZ analogues, and CEI at various pH values (**Supplementary Fig. 14**), allowing us to determine the p*K*_a_ values and monitor the protonation of individual ionizable groups in both CTZ and the CEI (**Supplementary Fig. 15**; **Supplementary Table 5**). As shown in **Fig. 5a**, comparison of the protonation states of CTZ and CEI with the pH profile of the rate of chemiluminescent CTZ autooxidation revealed that no CTZ conversion occurred when the substrate was fully protonated. At pH values above the p*K*_a_ of the CEI amide group, the conversion proceeded but luminescence was significantly attenuated (**Supplementary Fig. 16**). We thus showed that the O10-deprotonated form of CTZ (or its N7-deprotonated tautomer) and the amide-protonated CEI are critical for efficient oxidation and bright luminescence, which is consistent with the crystallographic results.

**Fig. 5.**
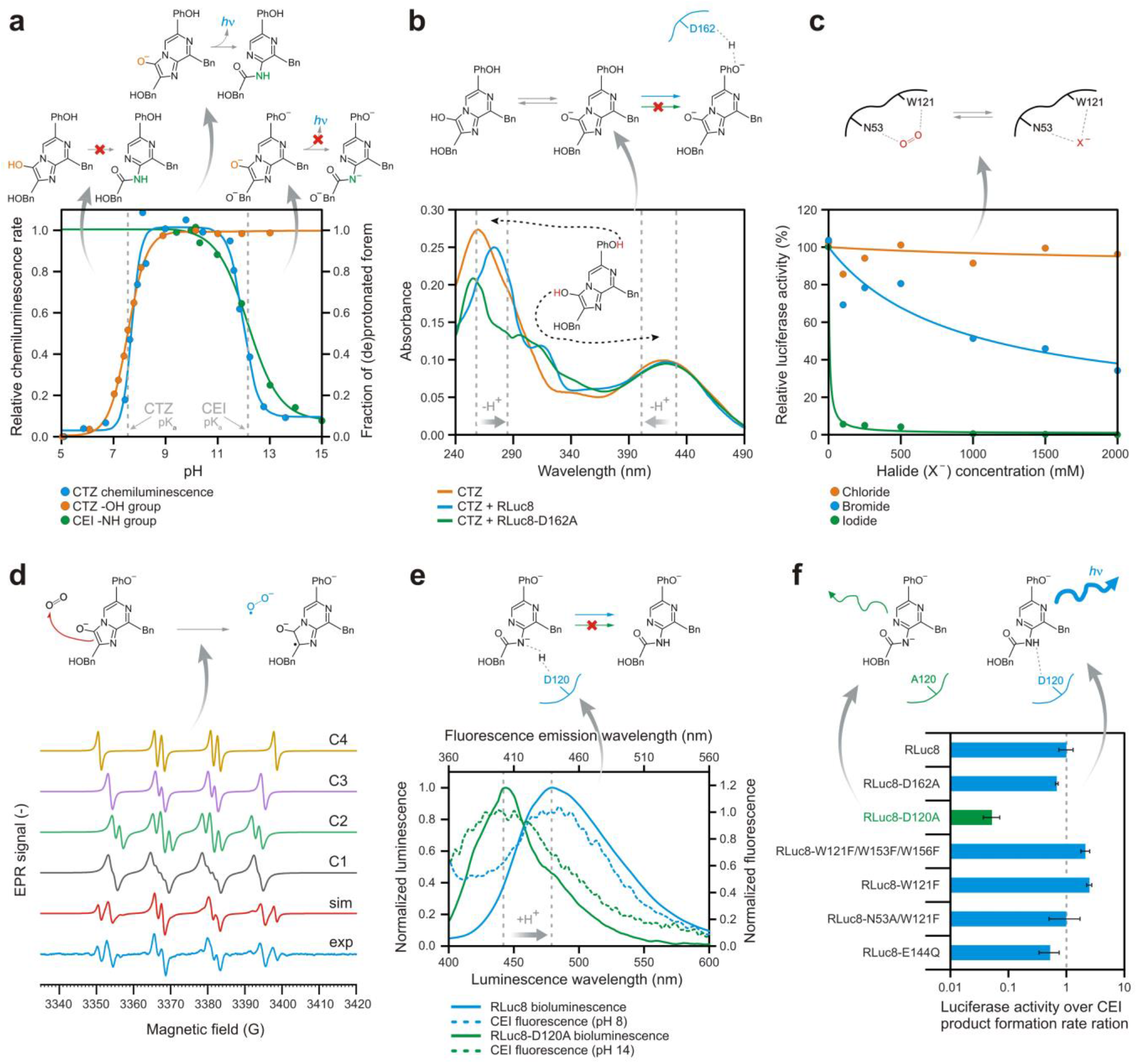
Spectroscopic dissection of the mechanism of *Renilla*-type bioluminescence. (**a**) Comparison of pH-titrated protonation states of coelenterazine (CTZ) and coelenteramide (CEI), and a pH profile of the chemiluminescent autooxidation rate of CTZ. Autooxidation is not observed at pH values below the p*K*_a_ of CTZ. When the pH is above the p*K*_a_ of CEI, autooxidation proceeds (see Figure S15) but luminescence is attenuated, highlighting the importance of deprotonated CTZ and protonated CEI for efficient luminescence. (**b**) Changes in the protonation state of CTZ upon anoxic binding to RLuc8 and RLuc8-D162A. The O10-hydroxy group is half-deprotonated in solution at physiological pH and no further enzymatic deprotonation is observed while the 6-PhOH group is fully protonated in solution and is actively deprotonated by the aspartate D162 upon binding. (**c**) Inhibition of the luciferase activity in the presence of halide (X^-^) anions. The activity is efficiently inhibited by bromide and iodide ions, indicating competition with oxygen for the halide stabilizing residues (N53 and W121). (**d**) A spin-trapping experiment using DMPO to study CTZ autooxidation. The experimental EPR spectrum (black line) was simulated (red line) with 4 different components (C1 to C4) whose hyperfine coupling constants are shown in **Supplementary Table 6**. The EPR signal reveals the presence of both superoxide and carbon-based radicals, suggesting that CTZ oxidation proceeds via a charge transfer radical mechanism. (**e**) Comparison of the bioluminescence spectra of RLuc8 and RLuc8-D120A with the fluorescence spectra of CEI at different pH values. The shift of the RLuc8-D120A luminescence maximum corresponds to the shift of the deprotonated CEI fluorescence maximum, demonstrating the role of aspartate D120 in reprotonating the CEI product. (**f**) Luminescence to product formation rate ratios for selected RLuc8 mutants. RLuc8-D120A generates significantly less luminescence per formed CEI molecule than the parent RLuc8, indicating that emission in this mutant occurs mainly from the less-luminescent deprotonated CEI. This result confirms the role of D120 in tuning the emission wavelength by ensuring the reprotonation of CEI.

Our co-crystal structures revealed that no protein residue could potentially sequester a hydrogen atom from the N7-CTZ form. The deprotonation of the O10-CTZ form could theoretically be mediated by the halide-stabilizing tryptophan W121 (AncFT W119), but this scenario does not seem to be favorable. Indeed, our experiments showed that there was no detectable increase in the deprotonation of this group upon mixing RLuc8 with CTZ under anaerobic conditions (**Fig. 5b**). This makes sense because the p*K*_a_ of this group is 7.55, which is close the physiological pH. Consequently, around half of the CTZ molecules in the bulk solvent will exist in the deprotonated form, so enzymatic deprotonation is not required. We conclude that CTZ enters the luciferase active site with its imidazopyrazinone core deprotonated. In contrast, the capping 6-(*p*-HOPh) group of CTZ was deprotonated by RLuc8 after binding (**Fig. 5b**), indicating that the α4 helix aspartate D162 (AncFT D160) could mediate this deprotonation. This assumption was validated by the finding that the RLuc8-D162A mutant cannot deprotonate the 6-(*p*-HOPh) group (**Fig. 5b**). Our data thus confirmed that the nucleophilic D162 residue (AncFT D160) plays a key role in fine-tuning the emission wavelength by generating a negatively charged emitter, i.e., the 6-*p*-hydroxyphenolate CEI ion.

### Halide ions compete with O_2_ for the enzyme binding site

Molecular oxygen (O_2_), which is a co-substrate of the luciferase reaction, was previously shown to be bound between two halide-stabilizing residues, N51 and W119 in the reconstructed ancestral enzyme Anc^HLD-RLuc^.^9^ This halide binding site is typically occupied by a halide ion during the dehalogenation reaction in haloalkane dehalogenases.^12,37^ To validate this finding, we measured the luciferase activity of RLuc8 in the presence of various halide anions (**Fig. 5c**). The tested halides had substantial inhibitory effects, suggesting that they compete with O_2_ for the halide-stabilizing residues. Bromide and iodide ions exhibited stronger inhibitory effects than chloride ions (**Fig. 5c**).

### Coelenterazine oxidation proceeds via a superoxide radical

Bui and Steiner have suggested that proper mutual positioning of the substrate and molecular oxygen in the cofactor-independent monooxygenase active site allows O_2_ to attack the deprotonated CTZ via a radical mechanism^40^, although no experimental evidence supporting this hypothesis has been presented. We therefore used EPR spectroscopy combined with spin trapping to capture a putative superoxide radical intermediate. The EPR-silent compound 5,5-dimethyl-1-pyrroline-N-oxide (DMPO) was used as a spin trap agent to form the EPR-active adducts DMPO-O^2-^ and/or DMPO-OOH by trapping the superoxide species generated in the CTZ luminescence reaction. The EPR spectrum obtained after subtracting the background from a control experiment (**Supplementary Fig. 17**) is shown as a black line in **Fig. 5d** together with a simulated spectrum (red line) obtained by combining four components (C1-4) representing different radical species in the sample. The spectra are shown in normalized form in **Fig. 5d**, but they did not contribute equally to the final spectrum; their individual contributions are shown in **Supplementary Table 6** together with the hyperfine coupling constants and the g-factor for each component. Component C1 has the typical parameters of a superoxide radical^41–45^, confirming the generation of superoxide in the reaction. Component C2 can also be attributed to a superoxide radical, despite a disagreement of *a_H_^γ^* for this species.^46^ Component C3 can be attributed to a hydroxyl radical, which is expected to be present due to the well-known dismutation of DMPO-OOH into DMPO-OH.^45,47^ Component C4 has typical features of a carbon-based radical but could not be attributed to any particular chemical species. To our knowledge, this is the first experimental evidence of superoxide radical generation during DMSO-activated CTZ chemiluminescence (**Fig. 5d**). These findings support the proposal that CTZ conversion proceeds via a charge-transfer radical mechanism.

### An aspartate of the conserved proton-relay system protonates the CEI ion

As reported previously^48^ and verified by our data (**Fig. 5a**), the excited CEI product must be protonated at the amide nitrogen to avoid pronounced attenuation of its light emission. This CEI reprotonation must occur inside the enzymatic pocket and not in bulk solvent because efficient light emission requires a hydrophobic environment and restricted flexibility^49,50^. This was confirmed by our observations showing that the bioluminescence spectrum corresponds to the fluorescence emission spectrum of the enzyme-bound CEI molecule rather than free CEI (**Supplementary Fig. 18**). Our co-crystal structures suggest that the residue responsible for this reprotonation is probably aspartate 120 (AncFT D118), which is part of a previously discussed conserved catalytic glutamate-histidine-aspartate triad. The side chain carboxylate of this aspartate is located in close proximity (3.3 to 3.6 Å) to the amide nitrogen of the bound CEI. The proton relay system conserved in the HLD fold thus seems to be critical in this final reprotonation step during the luciferase reaction. Interestingly, the bioluminescence spectrum of CEI in the RLuc8-D120A mutant was blue-shifted by ~50 nm, corresponding to the difference in the emission wavelengths of protonated and deprotonated CEI (**Fig. 5e**). This observation agrees with the results of Shimomura and Teranishi^20^, who found that the luminescence of the amide-deprotonated emitter has an emission maximum of 435–458 nm. Surprisingly, RLuc8-D120A yielded a significantly lower luminescence yield per molecule of generated CEI (**Fig. 5f**). The attenuated light output (**Fig. 5f** and **Supplementary Fig. 16**) and shifted emission spectrum (**Figs. 5e** and **6a**,**b**) are additional indicators of emission from the deprotonated CEI form, suggesting that the RLuc8-D120A mutant cannot effectively perform the reprotonation step and thus confirming the proton-transfer role of residue D120 (AncFT D118). Finally, the ratio of CNM to CEI obtained by the action of CTZ RLuc8-D120A on CTZ was 17 times higher than that achieved with RLuc8 (**Supplementary Fig. 19**). This is consistent with our co-crystal structure of RLuc-D120A, in which the CNM molecule is unambiguously present in the active site (**Fig. 3c**).

**Fig. 6.**
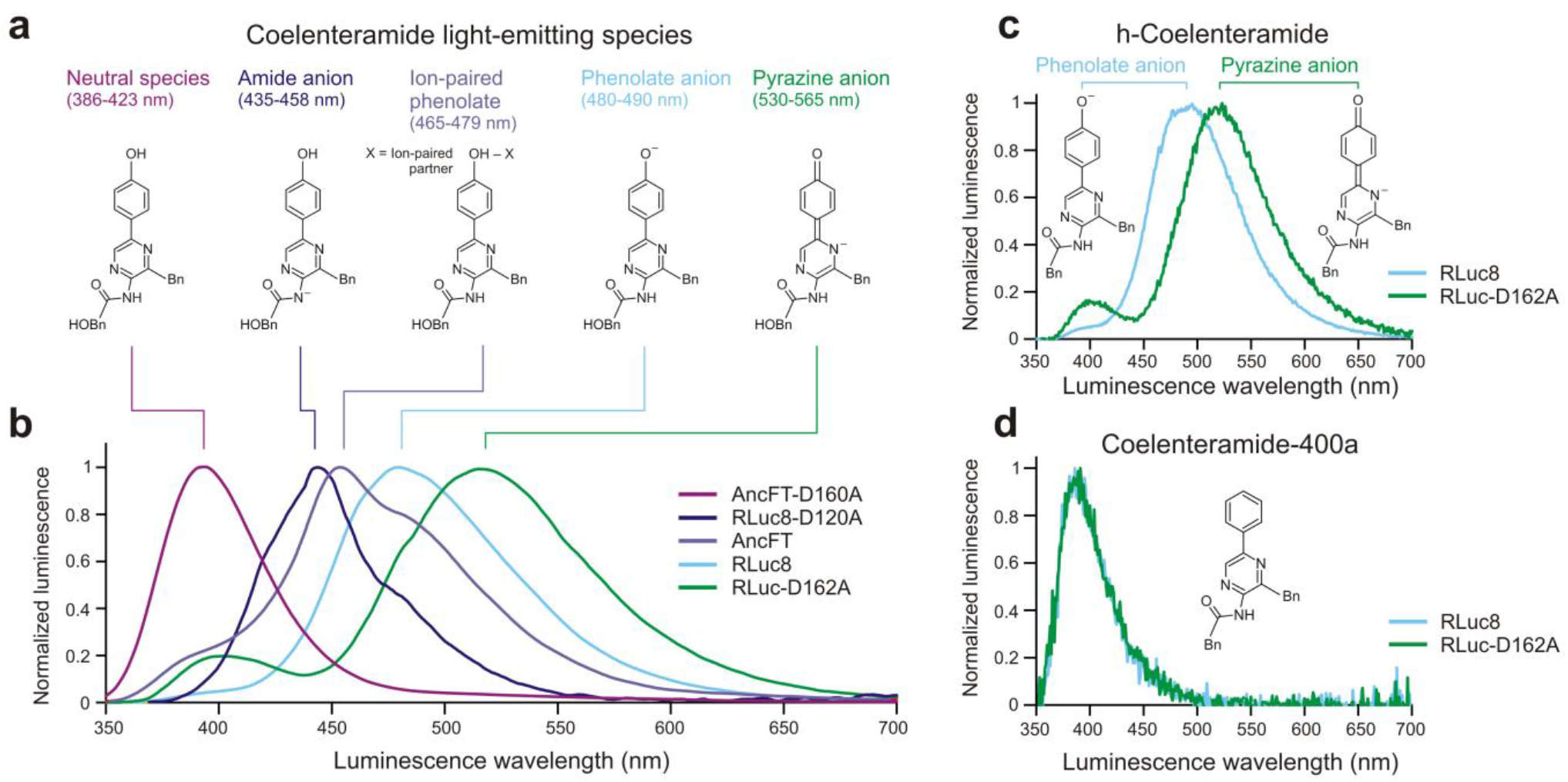
Emission spectra of RLuc8 and AncFT luciferases with native CTZ and its derivatives. (**a**) Light-emitting CEI species. (**b**) Emission spectra of AncFT, AncFT-D160A, RLuc8, RLuc8-D120A and RLuc-D162A luciferase variants with native CTZ. (**c**) Emission spectra of RLuc8 and RLuc8-D162A with h-CTZ. (**d**) Emission spectra of RLuc8 and RLuc8-D162A with CTZ-400a.

### Tuning the electronic state of CEI to favor blue light emission

The *Renilla* luciferase is known for emitting blue light; its emission maximum is ~480 nm.^7,8,15^ Interestingly, the emission of AncFT is slightly shifted toward higher energy wavelengths, with an emission peak at ~450 nm (**Fig. 6a**). It has been suggested that the CEI formed by the action of these enzymes on CTZ features an electron in an excited state, which loses energy and returns to the ground electronic state via the release of a visible photon. The wavelength of the emitted photon depends directly on the energy difference between the excited and ground states, which in turn depends directly on the environment around the CEI.^15^ Although early studies suggested that the CEI exists as an amide anion when it is in the active site pocket^7,20,51^, more recent works have postulated that the phenolate anion is the blue emitter in bioluminescence (**Fig. 6a**).^15,52–54^

Although previous structural studies did not identify a residue that could tune the electronic state of the CEI product^9,13–15^, our structures suggest that this role may be played by an aspartate localized at the rim of the enzymatic pocket. In the CEI-bound AncFT complex, a side chain carboxylate of D160 (RLuc8 D162) is in close proximity (~3 Å) to a hydroxyl group of the R^2^ 6-(*p*-hydroxyphenyl) substituent, suggesting potential hydrogen bonding between the two (**Fig. 2** and **Supplementary Fig. 11**). Similar hydrogen bonding interactions are possible in RLuc8 (**Fig. 3**). Supporting this hypothesis, mutation of this aspartate to alanine caused significant shifts in the emission maxima of both AncFT and RLuc8 (**Fig. 6b**). Specifically, AncFT-D160A emits light at the ultraviolet edge, with a single-peaked maximum at ~390 nm. Conversely, RLuc8-D162A has a two-peak emission spectrum, with a smaller peak at ~400 nm and a major green-shifted peak at ~520 nm. This two-peak spectrum may be associated with a closed-to-open transition of its malleable enzymatic cavity. Furthermore, this shift in the bioluminescence of the RLuc8-D162A mutant was only observed during the conversion of substrates whose R^2^ substituent bore a hydroxyl group amenable to deprotonation (**Fig. 6b,c**); it was not seen with CTZ-400a, which lacks this group (**Fig. 6d**). These observations strongly support the postulated role of D162 in deprotonating this specific substituent. Collectively, our results clearly identify the aspartate residue responsible for fine-tuning the electronic state of the CEI product.

### A blueprint for the reaction mechanism of *Renilla*-type bioluminescence

The results presented above allowed us to delineate a catalytic mechanism for the monooxygenation of coelenterazine by *Renilla*-type luciferases. Co-crystal structures of azaCTZ- and CEI-bound AncFT luciferase in which both ligands were captured in catalytically favored conformations played a vital role in this process. By considering these two structures, we were able to model the binding modes of CTZ and the more short-lived intermediates 2-peroxy-CTZ and CTZ dioxetanone (**Fig. 7a**). The proposed catalytic reaction mechanism is depicted schematically in **Fig. 7b**. The mechanism begins with the entry of the deprotonated form of CTZ into the enzyme active site through the main p1 tunnel. The imidazopyrazinone core of CTZ is readily deprotonated in solution because its p*K*_a_ of 7.55 is close to the physiological pH (**Fig. 5a,b**). Upon binding, the −OH group of the R^2^ 6-(*para*-hydroxyphenyl) substituent is deprotonated by aspartate 160 to give the dianionic O10-CTZ, which affects the emission maximum of the emitted light. We have previously shown^9^ that a dioxygen binding site in RLuc8 overlaps with the halide-binding site of related HLDs and that the binding of molecular oxygen at this site is compatible with the binding mode of CTZ in the enzymatic pocket. In the ternary Michaelis complex obtained after binding of both CTZ and molecular oxygen, the side chains of N51 (RLuc8 N53) and W119 (RLuc8 W121)^9^ position the co-substrate (O_2_) such that it can be directly attacked by the C2 carbon atom of the activated dianion. The initial interaction occurs via a charge-transfer mechanism that generates radical intermediates. The next steps are radical pairing and termination to form a 2-peroxy-CTZ anion that is then cyclized via a nucleophilic addition-elimination mechanism to form a highly unstable energetic dioxetanone structure with a deprotonated amide group. At this stage, the amide group must be protonated by D118 (RLuc8 D120) to ensure that luminescence occurs from the protonated form of CEI rather than the significantly less luminescent deprotonated CEI product. The presence of D118 also prevents unwanted hydrolysis of the amide bond by a water molecule, which yields the CNM side-product. Following this reprotonation step, the unstable dioxetanone ring decomposes, and the released energy excites the newly formed CEI product. As it returns to the ground state, the excited molecule releases a photon, resulting in the bioluminescence signal. The residue D118 is then reprotonated by H283 (RLuc8 H285) (**Fig. 7b**).

**Fig. 7.**
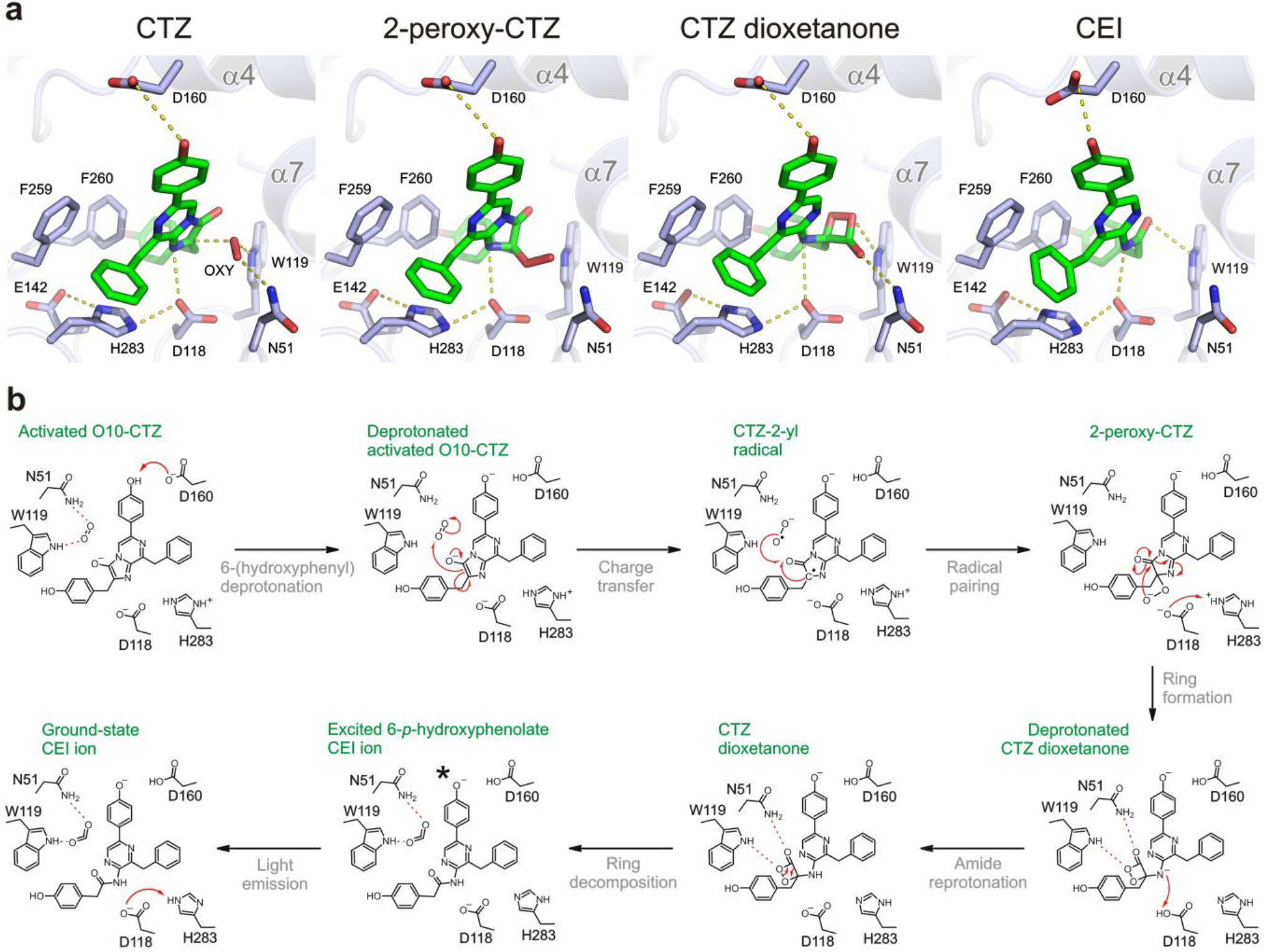
A proposed catalytic mechanism of coelenterazine-powered *Renilla*-type bioluminescence. (**a**) Close-up views of the AncFT active site center with modelled coelenterazine (CTZ), 2-peroxy-coelenterazine (2-peroxy-CTZ), CTZ dioxetanone, and coelenteramide (CEI) ligands. Molecular oxygen (OXY) and selected protein residues are shown as sticks; hydrogen bonds are shown as dashed lines. (**b**) CTZ enters the enzyme active site with a deprotonated imidazopyrazinone core because the p*K*_a_ of the core (7.55) is very close to the physiological pH, as demonstrated experimentally in this work. Upon the binding, the −OH group of the C6-(*p*-hydroxyphenyl) substituent is deprotonated by D160 to form the activated dianion O10-CTZ, which affects the emission maximum of emitted light. In the ternary Michaelis complex, the side chains of N51 and W119 position a co-substrate molecule (dioxygen) such that it can be directly attacked by the C2 carbon of O10-CTZ. Their initial interaction occurs via a charge-transfer radical mechanism. The next step involves radical pairing and termination to form the 2-peroxy-CTZ anion, which then cyclizes via a nucleophilic addition-elimination mechanism. This step yields a highly unstable and energetic dioxetanone structure with a deprotonated amide group. At this point the amide group must be protonated by D118 to avoid significant attenuation of the luminescence signal due to the formation of the deprotonated CEI product. D118 also prevents hydrolysis of the amide bond by a water molecule, which would yield the side-product coelenteramine (CNM). After reprotonation, the unstable dioxetanone ring decomposes and the released energy excites the newly formed CEI product. As it returns to the ground state, the excited molecule releases a photon, representing the bioluminescence signal, together with the final products - ground-state CEI and carbon dioxide. The residue D118 is reprotonated via an interaction with H283.

## Discussion

The CTZ-utilizing *Renilla* luciferase is one of the most popular reporter systems used in biological research and bioimaging technologies. However, despite intensive effort^7–9,13–15,26,26,38,39^, the molecular details of its reaction mechanism remained unknown, largely due to a lack of structural data on its E.S and E.P complexes. In the first structure reported by Loening and co-workers^14^, CEI is bound outside of the RLuc8 active-site pocket, and is involved in crystallographic contacts, which makes it difficult to assess the biological relevance of this binding mode. More recently, we captured CEI bound in the RLuc8 enzymatic pocket but the wide opening of the active site cavity in this structure again made it impossible to infer the reaction mechanism at the molecular level.^13^ In this work, we circumvented these problems by using (i) the stabilized AncFT^13^ surrogate enzyme and (ii) a new non-oxidizable coelenterazine analogue azacoelenterazine (azaCTZ), to decipher the mechanism of bioluminescence catalyzed by α/β-hydrolase fold luciferases.

Our kinetic experiments confirmed that azaCTZ acts as a non-oxidizable substrate analogue in AncFT and RLuc8, and is thus a valuable tool for studying their pre-catalytic Michaelis complexes by X-ray crystallography. A similar strategy using a different CTZ derivative was recently used to probe the substrate-binding site and mechanism of the Ca^2+^-regulated photoprotein aequorin.^55^ We have also proposed a detailed reaction mechanism for *Renilla*-type bioluminescence that was inferred by considering multiple co-crystal structures of AncFT and RLuc8 luciferases and is supported by the results of mutagenesis, spectroscopic, and computational experiments. We show that CTZ adopts a Y-shaped conformation in the active sites of these enzymes, which is required for proper positioning of its imidazopyrazinone core in the enzymatic pocket. Notably, the substrate-binding mode seen in our structures differs markedly from that predicted by Loening and coworkers^15^. We were unable to identify any residue that could potentially initiate the oxidation reaction by deprotonating the bound CTZ substrate at the N7 nitrogen or the O10 oxygen. However, by monitoring the dependence of CTZ autooxidation and chemiluminescence on the pH, we showed that only the O10-deprotonated form of CTZ (or its N7-deprotonated tautomer) can undergo efficient oxidation. Since the first p*K*_a_ of CTZ is 7.55, which is close to the physiological pH, the O10-deprotonated form will be abundant in the bulk solvent and no enzymatic deprotonation is required, in accordance with the calculations of Griffiths and coworkers.^23^ We thus conclude that CTZ enters the enzymatic pocket with its imidazopyrazinone core deprotonated at O10 and binds in an orientation that positions its C2 carbon perfectly for an attack on the bound co-substrate dioxygen via a charge-transfer process that generates superoxide and CTZ radicals. We also present EPR spectroscopic evidence for the participation of the superoxide anion radical in CTZ oxidation, which appears to be a common feature of cofactor-independent bioluminescence.^24^

After radical pairing and cyclization, an unstable dioxetanone intermediate with a deprotonated amide group is formed. Our co-crystal structures reveal that an aspartate of the conserved catalytic triad (Asp-His-Glu) protonates this intermediate, which is required to avoid significant attenuation of the luminescence signal. Unexpectedly, an aspartate-to-alanine mutation resulted in the formation of CNM rather than CEI. Our results thus demonstrate how a tiny change in enzymatic pocket can alter the CTZ oxidation pathway to favor CNM, which was recently identified as a major product of CTZ oxidation in the photoprotein pholasin.^22^ Moreover, we showed that the evolutionarily conserved and functionally important catalytic pentad of HLDs^12,37^ is also important in key chemical steps of the CTZ bioluminescence catalyzed by α/β-hydrolase fold luciferases. Finally, we identified an additional aspartate at the rim of the catalytic pocket that is not critical for the catalysis but is responsible for fine-tuning the electronic state of the CEI product. The electronic state of the CEI phenolate ion is responsible for the generation of blue light with an emission peak at ~480 nm^20^, which is required for energy transfer to green fluorescent protein (GFP) via bioluminescence resonance energy transfer (BRET), which is employed by *R. reniformis.* Together with our previous results^9,13^, the findings presented here provide clear evidence that the evolutionary emergence of a functional luciferase resulted from the optimization of a pre-existing HLD-fold protein. Recent discoveries by other groups support this hypothesis and suggest that this may have happened multiple times via divergent evolution.^10,11,56,57^

The *Renilla* luciferase is a widely-used reporter, although its relatively low stability, product inhibition, and flash-type bioluminescence can limit its applications. We previously showed that the engineered protein AncFT binds the CTZ substrate tightly and exhibits markedly stable glow-type bioluminescence.^13^ Here we report structures that provide a rationale for these distinct behaviors. While the AncFT enzymatic pocket is conformationally constrained and shaped to tightly accommodate a single substrate, the RLuc8 pocket is conformationally malleable and can accommodate up to three ligands. We speculate that this rather unusual behavior of extant RLuc luciferases may be linked to the absence of its native interaction partners, CTZ-binding protein (CBP)^58,59^ and GFP^59,60^, which can modify its dynamics via protein-protein interactions.

In conclusion, the mechanistic insights into visible light production in the active sites of α/β-hydrolase fold luciferases presented herein will make it possible to extend the usefulness of these enzymes for science and society. In the near future, rational protein engineering and focused directed evolution will be used to generate customized luciferases for diverse bioluminescent technologies.

## Conflict of interest

The authors declare no competing financial interest.

## Authors’ contributions

R.B., G.G. and Y.L.J. designed, synthesized and characterized azacoelenterazine; A.S., M.S. and M.M. constructed and cloned DNA mutants and produced all recombinant proteins; D.P., A.S. and Z.P. performed luciferase and steady-state kinetics assays; A.S. and M.M. performed co-crystallization experiments, collected X-ray data and solved protein-ligand structures; M.T. and Z.P. carried out HPLC experiments; V.T.S., A.S., M.T., M.M. and P.N. conducted EPR experiments; M.T., Z.P., and P.C. performed oxygen-free assays and spectral experiments; G.P.P., J.D. and D.B. performed and analyzed MD simulations; M.M., Z.P. and J.D. designed the study and contributed to data interpretation. All authors contributed to the writing of the manuscript and preparation of tables and figures.

## Acknowledgements

The authors would like to express their thanks to the Czech Science Foundation (22-09853S) and the Czech Ministry of Education (INBIO CZ.02.1.01/0.0/0.0/16_026/0008451, RECETOX RI LM2018121, e-INFRA LM2018140). This project has received funding from the European Union’s Horizon 2020 research and innovation program (TEAMING 857560 and Sinfonia 814418) and the Marie Sklodowska-Curie Action (No. 792772). M. M. acknowledges financial support from GAMU of the Masaryk University (MUNI/H/1561/2018), and M. T. is a Brno Ph.D. Talent Scholarship holder funded by the Brno City Municipality. CIISB research infrastructure project (LM2018127) is acknowledged for financial support of the measurements at Biomolecular Interactions and Crystallization Core Facility. P. C. acknowledges research support from VISTEC, TSRI, and the NSRF via the Program Management Unit for Human Resources & Institutional Development, Research and Innovation (no. B05F640089) and the Global Partnership program. P. N. acknowledges ERC grant (No. 714850) under the European Union’s Horizon 2020 research and innovation program. G.G. acknowledges a PhD fellowship from the Université Paris Descartes, Sorbonne Paris Cité. The design and synthesis of azaCTZ also benefited from the Valoexpress funding calls of the Institut Pasteur. The crystallographic experiments were performed using the PXIII beamline at the Swiss Light Source (SLS) in Villigen (Switzerland). We are grateful to the members of the SLS synchrotron for the use of their beamline and help during data collection.

## ONLINE METHODS

### Synthesis and characterization of azacoelenterazine

^1^H NMR and ^13^C NMR spectra were recorded on a Bruker Avance 400 spectrometer at 400 MHz and 100 MHz, respectively. Shifts (δ) are given in ppm with respect to the TMS signal and cross-coupling constants (J) are given in Hertz. Column chromatography were performed either on Merck silica gel 60 (0.035 - 0.070 mm) or neutral alumina containing 1.5% of added water using a solvent pump and an automated collecting system driven by a UV detector set to 254 nm unless required otherwise. Sample deposition was carried out by absorption of the mixture to be purified on a small amount of the solid phase followed by its deposition of the top of the column. The low-resolution mass spectra were obtained on an Agilent 1200 series LC/MSD system using an Agilent Jet-Stream atmospheric electrospray ionization system and the high-resolution mass spectra (HRMS) were obtained using a Waters Micromass Q-Tof with an electrospray ion source. When specified, the anhydrous solvents used were purchased. Experiments under inert atmosphere were carried out by purging the glassware with a stream of dry argon. Then, an argon balloon, fitted with a needle, was used to insure a positive pressure of inert gas during the reaction. Unless stated otherwise, a purity of at least 95% was obtained for all the compounds by means of chromatography or recrystallization and this level of purity was established by TLC, LC/MS and NMR spectroscopy.

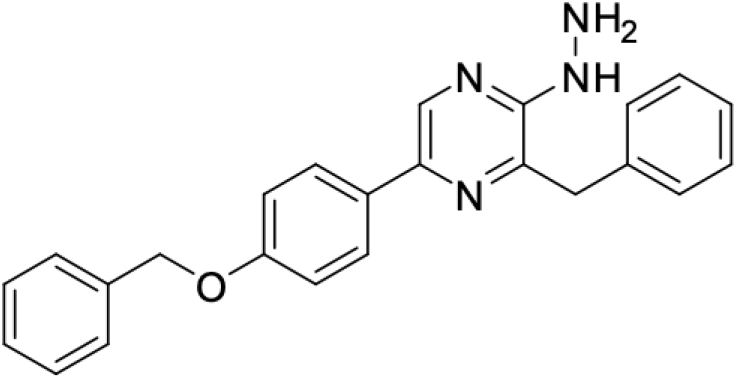

3-Benzyl-5-(4-(benzyloxy)phenyl)-2-hydrazinylpyrazine (**2**): In a 20 mL sealable Biotage vial, 3-benzyl-5-(4-(benzyloxy)phenyl)-2-chloropyrazine (**1**) (2.5 g, 6.46 mmol) and hydrazine hydrate (1.25 mL, 25.84 mmol) were dispersed in *n*-butanol (12 mL). The vial was sealed and heated in a microwave oven at 170 °C for 8 h. The resulting mixture was dispersed in distilled water (200 mL) during 15 minutes at room temperature. The precipitated was then filtered, washed with distilled water, cyclohexane and dried under vacuum at 55°C to give compound **2** as a yellow solid (2.25 g, 91%). ^1^H (DMSO-*d_6_*) *δ* 8.48 (s, 1H), 7.95 (s, 1H), 7.87 (d, *J* = 8.8 *Hz*, 2H), 7.48-7.06 (m, 12 H), 5.15 (s, 2H), 4.26 (s (br), 2H), 4.10 (s, 2H).^13^C (DMSO-*d_6_*) *δ* 158.5, 153.3, 141.2, 139.0, 138.3, 137.6, 135.6, 130.4, 129.5, 128.9, 128.6, 128.3, 128.2, 128.1, 126.6, 115.5, 69.7, 38.5. HRMS (m/z): [M+H]^+^ calcd for C_24_H_23_N_4_O: 383.1872, found: 383.1870.

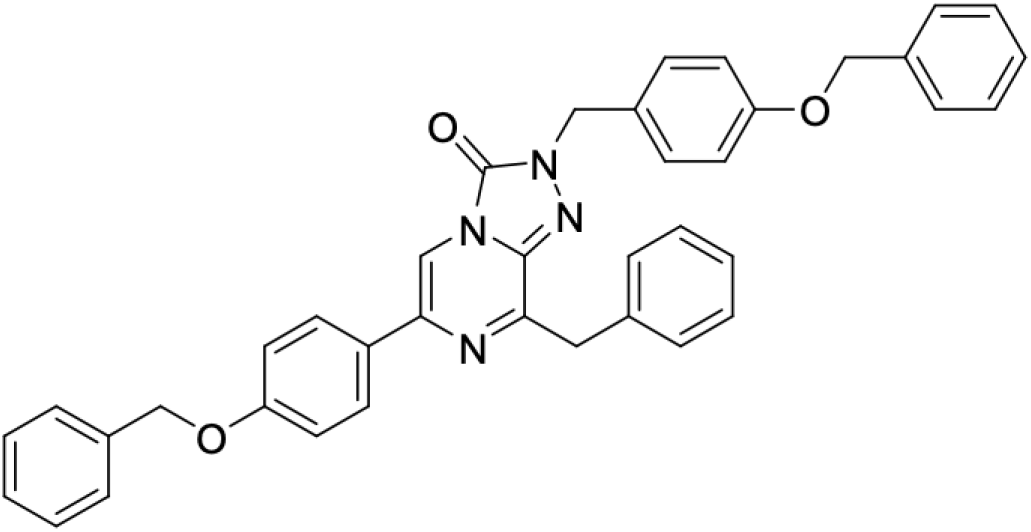

8-Benzyl-2-(4-(benzyloxy)benzyl)-6-(4-(benzyloxy)phenyl)-[1,2,4]triazolo[4,3-*a*]pyrazin-3(2*H*)-one (**5**). In a 250 mL round bottomed flask, compound **2** (1.66 g, 4.34 mmol) and 4-benzyloxybenzaldehyde (**3**) (0.99 g, 4.34 mmol) were dispersed in acetic acid (12 mL). The mixture was stirred at room temperature during 2 minutes. The resulting solid was re-dissolved in dichloromethane (40 mL) and cyanoborohydride (0.55 g, 8.7 mmol) was added. This was stirred at room temperature for 1 h. The solution was then dispersed in water and ethyl acetate, neutralized with 1 N NaOH (1 equivalent in regard with the acetic acid added). This was extracted with ethyl acetate thrice, the organic layer was washed with a saturated solution of sodium hydrogenocarbonate, distilled water, brine and dried over MgSO4. The solvent was removed under vacuum, to give the crude hydrazine **4** (2.43 g) which was considered pure. Under an inert atmosphere, this was then dissolved in dry tetrahydrofuran (70 mL, dried over 4 Å molecular sieves) dry triethylamine (1.14 mL, 8.7 mmol) was added and then solid triphosgene (0.40 g, 1.44 mmol) before stirring at room temperature for 40 minutes. The resulting mixture was diluted with water, extracted twice with ethyl acetate and the organic layer washed with water, brine, dried over magnesium sulfate and concentrated under vacuum to give a solid residue. A chromatography over silica gel (cyclohexane - ethyl acetate 4/1) gave a fraction containing pure compound **5** as a white solid (0.73 g, 27%) and an additional pure fraction (0.42 g, 16%) was obtained by washing the column with ethyl acetate. ^1^H (CDCl_3_) *δ* 7.83-7.80 (m, 3H), 7.51-7.23 (m, 17H), 7.08-7.07 (m, 2H), 6.99-6.95 (m, 2H), 5.17 (s, 2H), 5.14 (s, 2H), 5.08 (s, 2H), 4.34 (s, 2H). ^13^C (CDCl_3_) *δ* 159.4, 158.8, 153.8, 148.5, 136.9, 136.8, 136.3, 136.1, 135.4, 130.0, 129.7, 128.7, 128.6, 128.5, 128.4, 128.1, 128.0, 127.5, 127.4, 127.1, 126.9, 115.3, 115.1, 108.2, 70.2, 70.1, 49.7, 39.6. HRMS (m/z): [M+H]^+^ calcd for C_39_H_33_N_4_O_3_: 605.2552, found: 605.2548.

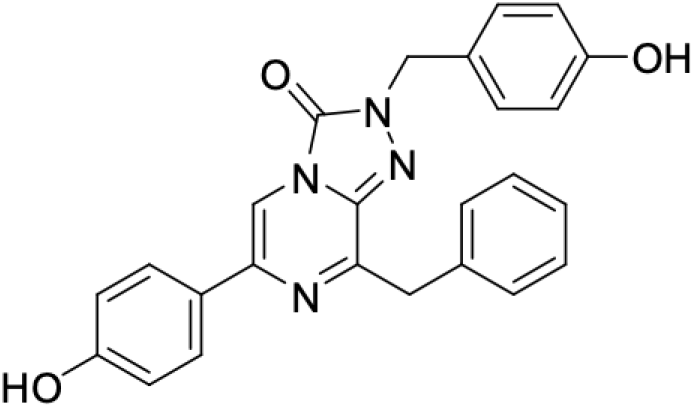

8-Benzyl-2-(4-hydroxybenzyl)-6-(4-hydroxyphenyl)-[1,2,4]triazolo[4,3-a]pyrazin-3(2H)-one (**azaCTZ**): In a 100 mL round bottomed flask, under inert atmosphere (argon) compound **5** (0.34 g, 0.56 mmol) was dissolved in anhydrous dichloromethane (5 mL). This solution was cooled to 0°C and a 1 N solution of boron trichloride in dichloromethane (1.2 mL, 1.18 mmol) was injected. This was allowed to warm back to room temperature and stirred for one hour. Water and ethyl acetate were then added and this was further stirred for 20 min before an extraction using ethyl acetate. The organic layer was washed with water, brine, dried over magnesium sulfate, concentrated under vacuum. A chromatography over silica gel (dichloromethane - ethanol 97/3) gave the azacoelenterazine (azaCTZ) a pale yellow powder (0.12 g, 50%) ^1^H (DMSO-*d_6_*) *δ* 9.63 (s, 1H), 9.42 (s, 1H), 8.05 (s, 1H), 7.81-7.78 (m, 2H), 7.39 (m, 2H), 7.31-7.14 (m, 5H), 6.82 (m, 2H), 6.74 (m, 2H), 5.07 (s, 2H), 4.26 (s, 2H). ^13^C (DMSO-*d_6_*) *δ* 158.4, 157.5, 153.5, 148.5, 136.8, 135.8, 135.5, 129.8, 129.7, 128.8, 127.4, 127.1, 126.9, 126.6, 116.0, 115.7, 108.6, 49.3, 39.1. HRMS (m/z): [M+H]^+^ calcd for C_25_H_21_N_4_O_3_: 425.1614, found: 425.1611.

### Molecular cloning and site-directed mutagenesis

All genes were amplified by a standard PCR and cloned into pET21b expression vector between NdeI and BamHI sites. Mutagenesis was carried out in two step PCR using Phusion polymerase (NEB, UK) according to the manufacturer’s protocol. The list of primers used is available in **Table S7**. After the mutagenesis reactions, the original template was removed by *DpnI* treatment (2 h at 37°C), followed by final inactivation of DpnI (NEB, USA) enzyme (20 min at 80°C). The resulting plasmids were transformed into chemocompetent *E. coli* Dh5α cells, plated on LB-agar (tryptone 20 g/l, yeast extract 10 g/l, NaCl 10 g/l, agar 15g/l) containing ampicillin (100 μg/ml), and incubated at 37°C overnight. Plasmids were isolated from three randomly selected colonies and sent for DNA sequencing (Eurofins Genomics, Germany).

### Overproduction and purification of recombinant enzymes

Overexpression of proteins from pET (amp^R^) was carried out in *E. coli* BL21 cells (NEB, USA) cultivated in LB medium supplemented with ampicillin (100 μg/ml) at 37 °C. Protein production was induced at 20 °C once the OD_600_ reached ~0.6 by adding IPTG to a final concentration of 0.5 mM. Prior to purification, cells were disrupted by sonication using Sonic Dismembrator Model 705 (Fisher Scientific, USA). Lysates were clarified by centrifugation (14000 rpm/1 h/4°C) using a Sigma 6-16K centrifuge (SciQuip, UK) equipped with 12166 rotor. Supernatants containing the recombinant His-tagged proteins at their C-terminal ends were metal-affinity purified using Ni-NTA Superflow Cartridge 5mL (Qiagen, Germany) installed on FPLC system (Bio-Rad Laboratories, USA) and equilibrated in a purification buffer A (500 mM NaCl, 10 mM imidazole 20 mM potassium phosphate buffer pH 7.5). Proteins were eluted with imidazole gradient and monomers were separated using gel permeation chromatography on Äkta FPLC (GE Healthcare, Sweden) equipped with HiLoad^™^ 16/600 Superdex^™^ 200 pg column (GE Healthcare, Sweden) and equilibrated with a GF buffer (50 mM NaCl, 10 mM Tris-HCl pH 7.5). Purified proteins were concentrated to final concentrations using Centrifugal Filter Units Amicon^R^ Ultra-15 Ultracel^R^-10K (Merck Millipore Ltd., Ireland). Purity of proteins was verified on SDS-PAGE. Concentration of protein samples was measured using DeNovix^R^ DS-11 Spectrophotometer (DeNovix Inc., USA).

### Conventional steady-state analysis

For reference enzyme kinetics measurement without the presence of an inhibitor, a 12-point concentration spectrum of CTZ solutions (0.011 μM, 0.022 μM, 0.11 μM, 0.22 μM, 0.44 μM, 0.88 μM, 1.1 μM, 2.2 μm, 4.4 μM, 5.3 μM, 7.04 μM and 8.8 μM) was prepared by either an injection of appropriate amount of CTZ stock solution into 10 ml of precooled 100 mM phosphate buffer (pH 7.5) or by dilution of a CTZ solution prepared in the described way with 100 mM phosphate buffer (pH 7.5). In order to measure the inhibition kinetics, three similar spectra of CTZ solutions were prepared with the difference of replacing the 100 mM phosphate buffer with 0.5 μM, 1 μM or 2 μM azaCTZ solution prepared by injection of appropriate amount of 1 mM azaCTZ ethanol solution into 200 ml of 100 mM phosphate buffer (pH = 7.5). The reactions were carried out in standard translucent 96-well microtitration plates at 37 °C and monitored by luminometer FLUOstar Omega (BMG LABTECH, Germany). The reaction mixture was composed of 90 % of CTZ/azaCTZ solution and 10 % of enzyme solution (c = 8.6 μg.ml^-1^). The reaction was started by automatic injection of 225 μl of CTZ/azaCTZ solution into 25 μl of enzyme solution already present in the well. After injection, intensity of luminescence of all wavelengths was measured for 8 s every 0.08 s. In order to acquire a luminescence baseline, before the substrate injection, the well was monitored for luminescence for 10 s with measurements every 0.5 s. The described measurement was carried out for each concentration of CTZ without the presence of azaCTZ and then for each concentration of CTZ and azaCTZ combination with three repeats for each measurement. The gain of the instrument was set to 3,250. A relative luminescence activity was obtained by integration of the observed luminescence intensity per second dependency and relating the result to used molar concentration of substrate and weight concentration of enzyme. The resulting luciferase activity was obtained in relative units of RLU.s^-1^.mg^-1^. The kinetic data were fitted using software GraphPad Prism 6 (GraphPad Software, Inc.).

### Numerical analysis of full conversion data

Solid CTZ was dissolved in ice-cold ethanol and stored under nitrogen atmosphere in dark glass vials at –20 °C. Before measurement, concentration and quality of the ethanol stock solution was verified spectrophotometrically. Series of buffer solutions with different azaCTZ concentration was prepared by manual injection of an appropriate volume of the ethanol stock solution into 10 ml of 100 mM phosphate buffer pH 7.5 immediately before the measurement. The reaction mixture was composed of 10 % (v/v) enzyme solution in 100 mM phosphate buffer pH 7.5 and 90 % (v/v) of buffer solution of CTZ. All reactions were carried out at 37 °C in microtiter plates using the microplate reader FLUOstar OPTIMA (BMG Labtech, Germany) set to broad-spectrum luminescence reading. Microplate well with pre-pipetted 25 μl of enzyme solution was first monitored for background light for 10 seconds, after which, 225 μl of a buffer solution with CTZ was added via an automatic syringe. The luminescence of the reaction mixture was then measured for the desired time until the luminescence intensity decreased under 0.5 % of its maximal measured value. Each reaction was performed in 3 repetitions. The gain of the reader was tailored to each enzyme separately, however, for each enzyme, all readings at different substrate concentrations were obtained using the same gain value. The recorded luminescence traces (rate vs. time) were transformed to reaction progress curves corresponding to cumulative luminescence in time. The transformed kinetic data (product vs. time) were fitted globally with the KinTek Explorer (KinTek Corporation, USA). The software allows for the input of a given kinetic model via a simple text description, and the program then derives the differential equations needed for numerical integration automatically. Numerical integration of rate equations searching a set of kinetic parameters that produce a minimum χ^2^ value was performed using the Bulirsch–Stoer algorithm with adaptive step size, and nonlinear regression to fit data was based on the Levenberg– Marquardt method.^1^ To account for fluctuations in experimental data, enzyme or substrate concentrations were slightly adjusted (± 5%) to derive best fits. Residuals were normalized by sigma value for each data point. The standard error (S.E.) was calculated from the covariance matrix during nonlinear regression. In addition to S.E. values, more rigorous analysis of the variation of the kinetic parameters was accomplished by confidence contour analysis by using FitSpace Explorer (KinTek Corporation, USA). In these analyses, the lower and upper limits for each parameter were derived from the confidence contour obtained from setting χ^2^ threshold at 0.95.^2^ The scaling factor, relating luminescence signal to product concentration, was applied as one of the fitted parameters, well constrained by end-point levels of kinetic traces recorded at particular substrate concentrations. Depletion of the available substrate after the reaction was ensured by repeated injection of the fresh enzyme.

### Anaerobic binding experiments

In the anaerobic glovebox, 200 μL of a differently concentrated enzyme was put into a separate well in a microplate and into each of these wells, 5 μL of CTZ or CEI stock solution was added. As a blank, 200 μL of the enzyme was mixed with 5 μL of ethanol for each enzyme concentration. The resulting concentration of an enzyme in the mixture ranged from 0 to 400 μM while the resulting concentration of CTZ and CEI was approximately 15 and 14 μM, respectively. The incubation buffer was 100 mM potassium phosphate buffer pH 7.5. The microplate was thoroughly covered with an air-tight UV transparent sealing to isolate wells from the surrounding environment and the plate was removed from the anaerobic glovebox. The extent of CTZ binding at each concentration was determined by measuring fluorescence spectra ranging from 480 to 580 nm upon excitation at 420 nm using the microplate spectrophotometer Varioskan LUX (Thermo Scientific, USA). Binding of CEI was monitored by measuring fluorescence spectra ranging from 350 to 700 nm upon excitation at 330 nm. Excitation bandwidth was set to 5 nm, measurement time to 100 ms, and the temperature inside the instrument to 30 °C.

The obtained fluorescence spectra were visually inspected for characteristic peaks, and the value of the maximal fluorescence intensity *Fluo*_max_ at these peaks was plotted against the enzyme concentration. The final value of the dissociation constant *K*_d_ was determined by fitting the dependence with the hyperbolic **Equation (1)** using the software Origin 6.1 (OriginLab, USA).

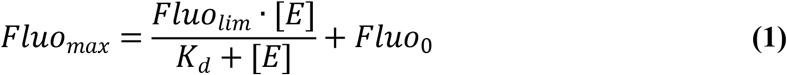

### Co-crystallization experiments

All crystallization experiments were done at 20°C using the hanging-drop vapor-diffusion method in EasyXtal 15-well plates (Qiagen, Germany) with drops equilibrated against 500 ul of reservoir solution.

For azaCTZ-bound AncFT complex, enzyme concentrated to ~9 mg/ml was mixed with azaCTZ in 1:4 molar enzyme-ligand ratio. Crystals were obtained after mixing 2 μl of enzyme-ligand mixture with 1 μl of the precipitant solution consisting of 0.2 M MgCl_2_, 0.1 M BisTris pH 6.5 and 21% PEG 3350. For CEI-bound AncFT complex, enzyme concentrated to ~22 mg/ml was mixed with native CTZ in 1:2 molar enzyme-ligand ratio. Crystals were obtained after mixing 1.5 μl of enzyme-ligand mixture with 1 μl of the precipitant solution consisting of 0.2 M Mg acetate, and 18% PEG 3350.

For azaCTZ-bound RLuc8-D162A complex, enzyme concentrated to ~10 mg/ml was mixed with azaCTZ in 1:2 molar enzyme-ligand ratio. Crystals were obtained after mixing 1 μl of enzyme-ligand mixture with 1 μl of the precipitant solution consisting of 0.1 M BisTris pH 6.5 and 25% PEG 3350. For CEI-bound RLuc8-D162A complex, enzyme concentrated to ~9 mg/ml was mixed with native CTZ in 1:3 molar enzyme-ligand ratio. Crystals were obtained after mixing 1 μl of enzyme-ligand mixture with 1 μl of the precipitant solution consisting of 0.04 M potassium phosphate monobasic and 16% PEG 8000. For CNM-bound RLuc8-D120A complex, enzyme concentrated to ~9 mg/ml was mixed with native CTZ in 1:3 molar enzyme-ligand ratio. Crystals were obtained after mixing 1 μl of enzyme-ligand mixture with 1 μl of the precipitant solution consisting of 0.2 M MgCl_2_, 0.1 M BisTris pH 6.5 and 20% PEG 3350.

All crystals were fished out, cryo-protected in the corresponding reservoir solutions supplemented with 20% glycerol, and flash-frozen in liquid nitrogen for X-ray data collection.

### Diffraction data collection and data processing

All X-ray data were collected at PXIII beamline at SLS Synchrotron (Villigen, CH) at the wavelength of 0.999 Å using a Pilatus 2M-F detector. The data were processed using XDS^3^, and Aimless^4^ was used for data merging. Initial phases were solved by molecular replacement using Phaser^5^ implemented in Phenix^6^ with *apo*-form AncFT (PDB ID: 6S97)^7^ and RLuc8 (PDB ID:6YN2)^7^ employed as search models. The refinement was carried out in cycles of automated refinement in phenix.refine program^8^ and manual model building in Coot^9^. The final models were validated using tools provided by Coot^9^ and Molprobity^10^. Visualizations of structural data were created using PyMOL^11^. Structural superposition was carried out using the secondary structure matching (SSM) superimpose tool in the Coot^9^. Atomic coordinates and structure factors of the enzyme-ligand complexes were deposited in the Protein Data Bank (www.wwpdb.org)^12^ under the PDB codes 7QXQ, 7QXR, 7OMD, 7OMR and 7OMO.

### Analysis and mapping of enzyme-ligand interactions

In general, enzyme-ligand interactions were studied in all monomers in the asymmetric units. Briefly, all residues within 4 Å of a ligand were found. Then all atoms within 4 Å of a ligand were colored by element to distinguish between hydrophobic interactions and hydrogen bonds. A π-π interaction was recognized as an interaction between two aromatic rings in which either (i) the angle between the ring planes is less than 30° and the distance between the ring centroids is less than 4.0 Å (face-to-face), or (ii) the angle between the ring planes is between 60° and 120° and the distance between the ring centroids is less than 5.0 Å (edge-to-face). Graphical visualizations were done in PyMOL.^11^ To prepare a 2D representation of ligand interactions with AncFT, the images of chemical structures were prepared in Chem3D Pro 12.0 (PerkinElmer, USA) software, exported as vector images to be further adjusted in CorelDRAW X6 (Corel Corporation, Canada) graphical software. The visual style for two-dimensional drawings was adapted from PoseView (BioSolveIT, Germany) software.^13^

### Luciferase activity measurements

Luciferase activity measurements were performed as described previously.^7,14^ Briefly, Luminescence activity measurement was performed with CTZ (≥95 %, for biochemistry, Carl Roth GmbH + Co. KG, Germany) as a substrate and determined using FLUOstar Omega Microplate reader (BMG Labtech, Germany) at 37 °C. Sample of 25 μL of purified enzyme was placed into the microtiter well. After baseline collection for 10 s, the luminescence reaction was initiated by addition of 225 μL of 8.8 μM CTZ in the reaction buffer (100 mM potassium phosphate buffer, pH 7.5). Luminescence was recorded for 72.5 s and each sample was measured in three replicates (gain 3250). The area of obtained luminescence intensities peak given in relative luminescence unit (RLU) was recalculated to RLU·mg^-1^·s^-1^.

### Determination of p*k*_a_ values of CTZ and CEI ionizable groups

The determination of all p*K*_a_ values was based on changes of absorbance spectra at different pH values. Ethanol stocks of either CTZ, h-CTZ, CTZ-400a, CTZ-Q170^15^, or CEI were diluted into 100 mM potassium phosphate buffer of various pH to obtain a pH dependent series of aliquots ranging from pH 3 to pH 14. Absorbance spectra from 200 nm to 800 nm with a 1 nm step were collected using the spectrophotometer Varioskan LUX (Thermo Scientific, USA) or Cary 100 UV-Vis (Agilent, USA). The resulting concentration of each of the CTZ variants in the buffer was approximately 45 μM while the resulting concentration of CEI was approximately 30 μM. To prevent spontaneous degradation mainly at basic pH values, CTZ samples were prepared anaerobically using the anaerobic glovebox Belle MR2 (Belle Technology, UK), and the spectra were measured in a hermetically sealed cuvette. The wavelength of the absorbance spectrum peak was plotted against the actual pH value, and the value of p*K*_a_ was determined by fitting the dependence with the sigmoid equation (**2**) where *w* represents the sigmoid width, *m* is the slope of a potential drift, and *λ*_low_ and *λ*_high_ stand for limiting wavelengths of peaks at low and high pH, respectively. All the nonlinear fitting was performed using the software Origin 6.1 (OriginLab, USA).

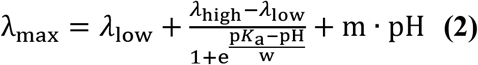

### Coelenterazine protonation changes upon anaerobic binding by luciferases

An aliquot of RLuc8 dissolved in oxygen-free 100 mM potassium phosphate buffer pH 7.5 was mixed with the anaerobic CTZ stock solution inside the anaerobic glovebox Belle MR2 (Belle Technology, UK). The concentration of the luciferase and the CTZ was approximately 105 μM and 16 μM, respectively. Anaerobic absorbance spectra from 200 nm to 800 nm with a 1 nm step were determined for each well using the microplate reader Varioskan LUX (Thermo Scientific, USA) and observed for any changes upon substrate binding.

### Coelenterazine conversion chemiluminescence rate and spectra at various pH

The rate of chemiluminescence was studied for oxidative decomposition of CTZ by DMSO in the presence of 10 % buffer of various pH. CTZ stock solution was added 10 % (v/v) mixture of 100 mM potassium phosphate buffer of various pH in DMSO and immediately after that, chemiluminescence signal was collected using the microplate reader Varioskan LUX (Thermo Scientific, USA). In addition, chemiluminescence spectra at each pH value (270–840 nm) and absorbance spectra of the fully reacted mixture (300–700 nm) were measured. The initial rate of the integrated chemiluminescence signal increase was plotted against the real pH value and fitted with the double-sigmoid curve.

### Bioluminescence and fluorescence spectra during and after enzymatic conversion

To reach both fully bound and nearly fully released substrate/product molecules states in the mixture with luciferase, either highly diluted or highly concentrated (in excess) enzyme was mixed with a CTZ variant in 100 mM potassium phosphate buffer pH 7.5. The resulting concentration of the enzyme was ca 8 nM (diluted sample) or ca 55 μM (concentrated sample). The procedure was repeated for each of the enzyme-ligand combinations for CTZ, h-CTZ, CTZ-400a, and CEI as ligands and RLuc8 and RLuc8-D162A as enzymes. After mixing, luminescence spectra from 300 nm to 800 nm were determined using the microplate reader Varioskan LUX (Thermo Scientific, USA). In addition, fluorescence spectra of the fully reacted mixtures were collected from 350 nm to 800 nm upon excitation at 330 nm.

### Coelenteramide and coelenteramine production during enzymatic reaction

Enzymatic reaction was initiated by mixing CTZ sample with an enzyme solution in 100 mM potassium phosphate buffer pH 7.5 and then incubated at 37 °C and 1,000 rpm in the thermal block ThermoMixer C (Eppendorf, Germany). The resulting concentration of CTZ ranged from 0 to 40 μM while the resulting concentration of the enzyme was 0.13 μM. The reaction was stopped by addition of 0.8 M sulfuric acid at various predefined times and extracted into ice-cold ethyl acetate. The extracted samples of CTZ, CEI, and CMN were separated and quantified using the HPLC instrument Agilent 1100 (Agilent Technologies, USA) equipped with the Kinetex 5μm PFP 100 Å reversed phase column (Phenomenex, USA).

Volume of 270 μL of CTZ sample was preincubated at 37 °C and 1,000 rpm for 20 seconds in the thermal block ThermoMixer C (Eppendorf, Germany) inside a glass vial. The reaction was initiated by manual addition of 30 μL of an enzyme solution, incubated for defined times and then stopped by addition of 200 μL of 0.8 M sulfuric acid. All the components of the reaction mixture were prepared in 100 mM potassium phosphate buffer pH 7.5 containing 6.2 % ethanol as a cosolvent. The final concentration of CTZ ranged from 0 to 40 μM while the final concentration of the enzyme was 4.7 μg/mL (≈ 0.13 μM).

### Electron paramagnetic resonance spectroscopy

The presence of the superoxide radical during the chemiluminescence reaction was detected via electron paramagnetic resonance (EPR) using the spin trapping technique.^16^ The EPR silent compound 5,5-Dimethyl-1-Pyrroline-N-Oxide (DMPO) was used as the spin trap agent to form the EPR active adducts DMPO-O^2-^ and/or DMPO-OOH by trapping the superoxide species generated in the reaction. The DMPO adducts are relatively stable nitroxide radicals and the trapped radicals can be identified according to the separation between the EPR absorption peaks caused by the hyperfine interaction of the electron with the nuclear spin around it. The splitting corresponds to the hyperfine coupling constants (denoted by *a*) that can be precisely calculated through spectral simulation. The simulations were performed using Easyspin^17^, a freely available package for EPR spectral simulations working for Matlab.^18^

The reactions were prepared in glass vials with a final volume of 100 μL containing CTZ (685 μM) and DMPO (200 mM) diluted in dimethyl sulfoxide (DMSO). 40 μL of 50 mM phosphate buffer solution (PBS) pH 7.5 were added to the reaction prior to EPR data acquisition. As a control, we measured a solution with the same content except for CTZ that was substituted by pure DMSO. The excess of DMPO was intended to avoid secondary radical formation in the reaction. The measured solution was pipetted into a quartz cuvette (flat cell) and measured at room temperature using a Magnettech X-band EPR spectrometer. The spectra were obtained with 10 mW microwave power and modulation of 0.9 G at 100 kHz. The presented results are the average of 36 accumulations of 120 G sweeps performed for 1 minute each.

### Computer modelling

The X-ray structures of the azaCTZ-bound AncFT (PDB ID: 7QXR) and CEI-bound AncFT (PDB ID: 7QXQ) complexes solved in this study were used to model E.CTZ, E.2-peroxy-CTZ, E.dioxetanone intermediate complexes. Then, PyMOL^11^ was used to model the RLuc complex structures by superimposing the *apo*-form RLuc8 (PDB ID: 2PSF11; chain B)^19^ and AncFT complexes, producing analogous RLuc8-ligand complexes for MD simulations.

### Energy minimization of structural models

The eight complex structures representing each ground-state of the putative reaction (four states for AncFT and four states for RLuc) were minimized using YASARA.^20^ We started by cleaning the structure and adding all the hydrogens that were missing at pH 7.5. The putative pentad of catalytic residues was visually inspected, and the D162 (D160 in AncFT) on the α4 helix was checked to confirm that their protonation states were as shown in Figure S19. We selected the AMBER FF14SB^21^ force field on YASARA and used the option “clean” that does the following: a) detects missing bonds, adds them, and assigns bond orders, b) adds missing cysteine bridges between close cysteine atoms, provided that their positions allow bridge formation, c) adds missing hydrogens to provide a starting point for the following analysis, and d) assigns force field parameters needed in the next steps. In the context of the AMBER force fields, ligands are automatically parameterized using GAFF2^22^ and AM1-BCC charges. After these steps, we ran structure minimization using the YASARA macro em_run.mcr to obtain the starting points for the production simulations. The minimization was considered as converging as soon as the energy improves by less than 0.05 kJ/mol per atom during 200 steps.

### Molecular dynamics simulations

For the analysis and optimization of substrate, intermediates, and product complexes, short molecular dynamics (MD) simulations were performed. The minimized structures of all complexes were used as starting points and 10 ns long production molecular dynamics simulations were performed using AMBER 14. These production MD simulations were run using YASARA’s macro, md_run.mcr. The systems were solvated in a cubical water box of explicit solvent, with density 0.997 g/ml, so that all atoms were at least 12 Å from the boundary of the box. Cl^-^ and Na^+^ ions were added to neutralize the protein’s charge and get a final concentration of 0.1 M. All the simulations employed periodic boundary conditions, the particle mesh Ewald method was used to treat interactions beyond a 10.5 Å cut-off, electrostatic interactions were suppressed more than 4 bond terms away from each other, and the smoothing and switching of van der Waals and electrostatic interactions were cut-off at 8 Å. For the MD simulations analysis, md_analyze.mcr was used for the RMSD and RMSF analysis. The distances between atoms were measured in PyMOL^11^. Three replicas of the production simulations were performed to ensure the significance and convergence of the simulation.

## Notes

### Competing Interest Statement

The authors have declared no competing interest.

