## Supplementary Data File for "A catalytic mechanism for *Renilla*-type bioluminescence"

#### Contents

Supplementary figures, tables and notes …………………………………………………. Pages 2 to 27
Supplementary references ……………………………………………………………....……… Page 28
NMR spectra ……………………………………………………………………………..… Pages 29 to 35

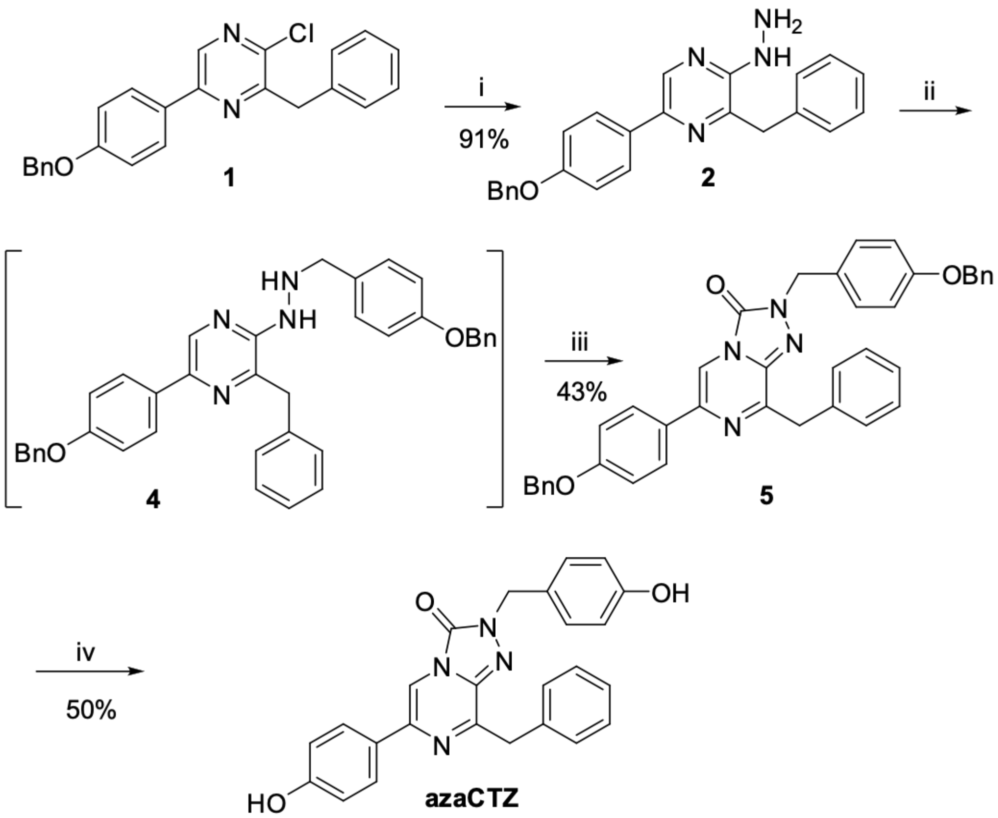


**Supplementary Figure 1.** Synthesis of azacoelenterazine (azaCTZ). Explanatory notes: (i) $NH_2NH_2$, nBuOH, 170 °C, MW; (ii) a 1) 4-$BnOC_6H_4CHO$ (**3**), AcOH, 2) $NaCNBH_3$, THF; (iii) $CCl_3COCCl_3$, THF; and (iv) $BCl_3$, $CH_2Cl_2$, -78 to 20 °C.

**Supplementary Note 1. Conventional steady-state analysis.**

The conventional steady-state analysis of AncFT inhibition by azaCTZ was determined by measuring initial reaction velocity at twelve different starting concentration of CTZ without and with different concentrations of azaCTZ (**Supplementary Figure 2a**). The double reciprocal (Lineweaver- Burk) plot of the kinetic data (**Supplementary Figure 2b**) showed a characteristic pattern indicating competitive inhibition of CTZ reaction by azaCTZ. The kinetic data were fitted using competitive inhibition model (**Supplementary Equation 1**) using nonlinear regression providing estimates for Michaelis constant ($K_m$ = 0.023 ± 0.002 μM), inhibition constant ($K_I$ = 0.012 ± 0.001 μM) and the relative value for maximal reaction velocity ($V_{max}$ = 0.91 ± 0.1 x$10^6$ RLU $s^{-1}$ $mg^{-1}$). The data also indicated weak substrate inhibition with $K_{SI}$ >> $K_m$. The values of kinetic constants obtained by conventional analysis (**Supplementary Table 1**) were subsequently used as initial values for more rigorous numerical analysis of full conversion curves.

$$v = \frac{V_{\mathrm{max}} \cdot [\mathrm{S}]}{K_{\mathrm{m}} \cdot \left(1 + \frac{[\mathrm{I}]}{K_{\mathrm{I}}}\right) + [\mathrm{S}] \cdot \left(1 + \frac{[\mathrm{S}]}{K_{\mathrm{SI}}}\right)}$$

*Supplementary Equation 1*

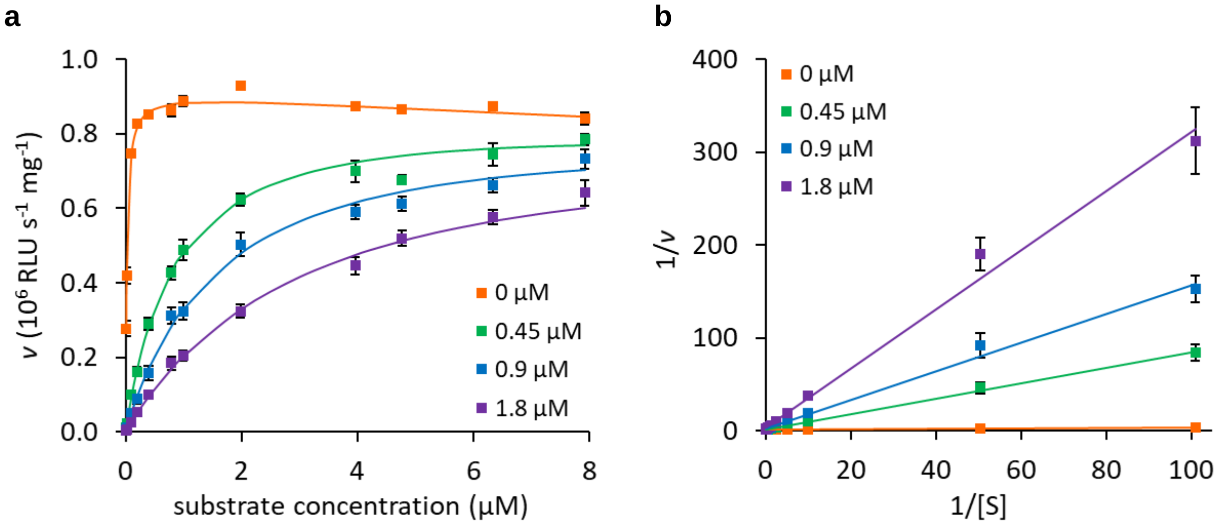


**Supplementary Figure 2.** Conventional steady-state analysis of AncFT inhibition by azaCTZ. (a) The initial velocity of AncFT reaction at a range of CTZ concentration analysed at the absence and presence of different concentrations of azaCTZ. Each data point is the average of three repetitions, the error bars represent standard deviation. The solid lines represent the best fit using **Supplementary Equation 1**. (**b**) The double reciprocal (Lineweaver-Burk) plot demonstrating the effect of azaCTZ on CTZ conversion catalyzed by AncFT. The solid lines represent individual linear fits.

**Supplementary Table 1.** Kinetic constants obtained by global data analysis for AncFT. The parameters were derived by fitting kinetic data using numerical integration of rate equations derived for competitive, uncompetitive, non-competitive, and mixed-type inhibition models described in **Supplementary Figure 3**, where $K_m$ is Michaelis constant, $k_{cat}$ is the turnover number, $K_P$ is the equilibrium dissociation constants for enzyme-product complex, $K_{SI}$ is equilibrium constant describing substrate inhibition, $K_{I1}$ and $K_{I2}$ is an inhibition equilibrium constant describing binding of the inhibitor to the free enzyme and enzyme-substrate/product complex, respectively. The kinetic parameters are reported as values obtained from the best fitted ± standard errors (S.E.). In addition, the lower and upper confidence limits at $\chi^2$ threshold 0.95 are summarized in the brackets for each parameter of the best fit.

| | **Competitive** | Uncompetitive I | Uncompetitive II | Noncompetitive | Mixed-type I | Mixed-type II |
|---|---|---|---|---|---|---|
| **$\chi^2$ x $10^7$** | **2.7** | 4.40 | 9.3 | 3.2 | 2.7 | 2.7 |
| **$K_m$ (μM)** | **0.042 ±0.001**<br>(0.026; 0.068) | 0.14 ±0.01 | 0.050 ±0.003 | 0.094 ±0.003 | 0.046 ±0.001 | 0.034 ±0.001 |
| **$k_{cat}$ ($s^{-1}$)** | **0.074 ±0.001**<br>(0.066; 0.085) | 0.120 ±0.003 | 0.069 ±0.001 | 0.100 ±0.001 | 0.076 ±0.001 | 0.068 ±0.001 |
| **$K_P$ (μM)** | **0.39 ±0.02**<br>(0.235; 0.621) | 0.63 ±0.02 | 0.87 ±0.07 | 0.55 ±0.02 | 0.40 ±0.02 | 0.41 ±0.01 |
| **$K_{I1}$ (μM)** | **0.092 ±0.002**<br>(0.060; 0.137) | n.a. | n.a. | 0.370 ±0.003 | 0.111 ±0.002 | 0.085 ±0.002 |
| **$K_{I2}$ (μM)** | n.a. | 0.186 ±0.004 | 0.024 ±0.001 | n.a. | 2.4 ±0.2 | 0.47 ±0.05 |
| **$K_{SI}$ (μM)** | **2.37 ±0.04**<br>(1.48; 4.15) | 0.70 ±0.03 | 3.4 ±0.3 | 1.00 ±0.03 | 2.11 ±0.04 | 3.24 ±0.07 |

[a] The dissociation constants were kept identical for non-competitive binding to all enzyme forms, free enzyme as well as enzyme-substrate/product complexes.
n.a. not applied

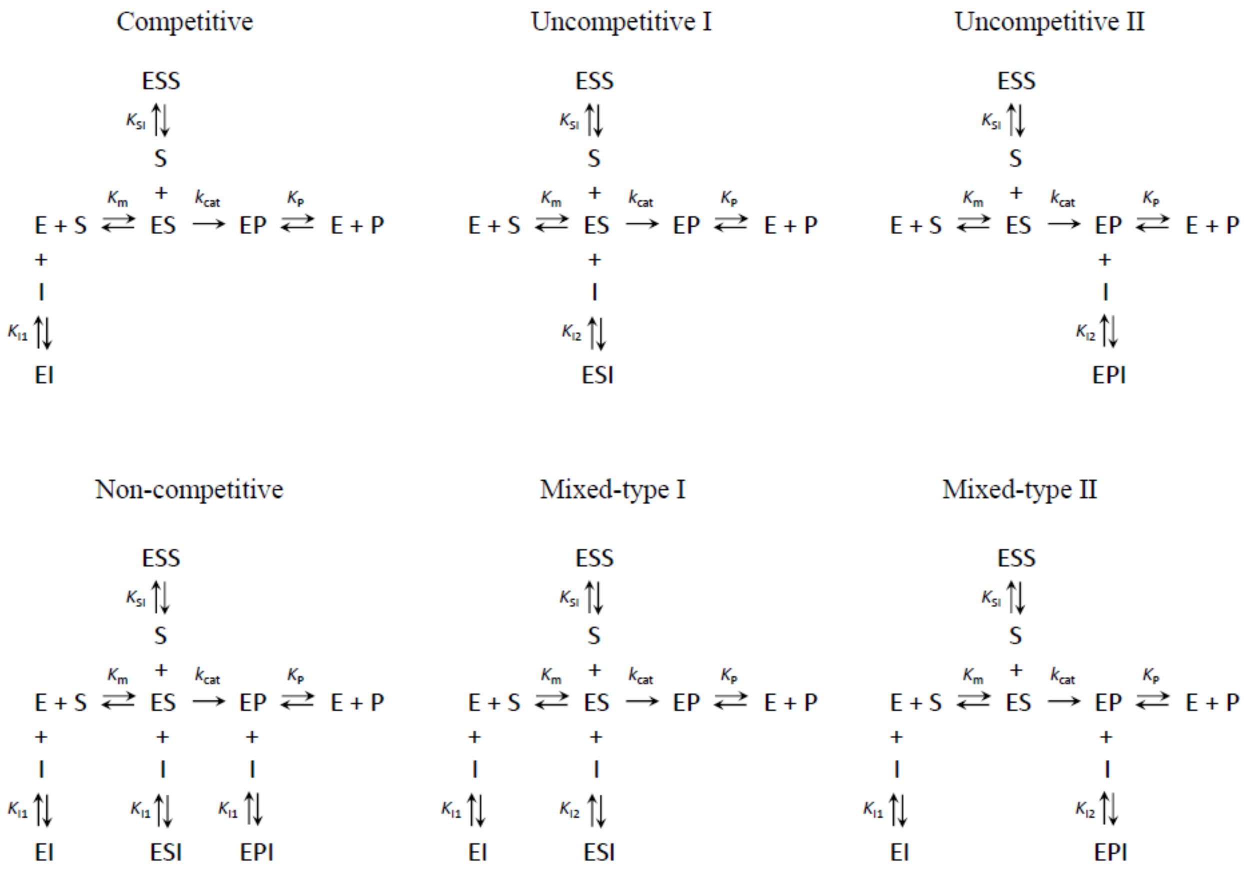


**Supplementary Figure 3.** Inhibition models. $K_m$ is Michaelis constant, $k_{cat}$ is the turnover number, $K_P$ is the equilibrium dissociation constants for enzyme-product complex, $K_{SI}$ is equilibrium constant describing substrate inhibition, $K_{I1}$ and $K_{I2}$ is an inhibition equilibrium constant describing binding of the inhibitor to the free enzyme and enzyme-substrate/product complex, respectively.

**Supplementary Note 2. Numerical analysis of full conversion data.**

Instead of multiple fitting steady-state data using a linear function and then next fitting the concentration dependence of the observed rate which cumulates fitting errors, during the numerical integration the raw data are fitted directly in a single step with fewer unknown variables, resulting in less error on the estimates for steady-state kinetic parameters.[1] Unlike the classical initial velocity analysis, which cannot provide the value of the turnover number without sophisticated luminometer calibration and quantum yield evaluation, the more rigorous numerical analysis of full conversion data (**Supplementary Figure 4**) provided precise estimates of all steady-state kinetic parameters of AncFT, the turnover number ($k_{cat}$ = 0.074 ± 0.001 $s^{-1}$), Michaelis constant ($K_m$ = 0.042 ± 0.001 μM), and additionally, the equilibrium dissociation constants for enzyme-product complex ($K_P$ = 0.39 ± 0.02 μM) thanks to the complete monitoring of the reaction far beyond the initial phases. The weak substrate inhibition observed in the conventional analysis was also well defined by numerical analyses of the full conversions data even though the substrate inhibitory effect was weak. The equilibrium constant ($K_{SI}$ = 2.37 ± 0.04 μM) for dissociation of the substrate inhibitory complex (ESS) was an order of magnitudes weaker in comparison to Michaelis constant. The kinetic parameters obtained by numerical methods (**Supplementary Table 1**) correspond well with previously reported values for the kinetics of AncFT with CTZ.[2]

Next, the numerical integration of rate equations allowed to simultaneously fit complex kinetic data recorded in the absence of azaCTZ with the full conversion signal traces recorded in the presence of azaCTZ. Global fitting provided a robust and accurate model that fully accounted for all experimental observations. In accordance with the initial observations from the conventional analysis, the numerical fitting provided the best solution for the model when azaCTZ competed directly for the same binding site with the substrate (**Supplementary Figure 4**). The confidence contour analysis showed that the parameters were well defined by the data, all kinetic constants were well constrained, the upper and lower limits derived from the confidence contour analysis were reasonably ranged (**Supplementary Figure 5**).

The fit of uncompetitive and noncompetitive models (**Supplementary Figure 3**) provided significantly worse correspondence with the measured data (**Supplementary Figure 6**) indicated also by increase in $\chi^2$ values (**Supplementary Table 1**). Also, additional extension of the competitive model to mixed-type did not bring an adequate improvement of the fit, $\chi^2$ value obtained for both mixed-type I and II models was identical to the value calculated for simple competitive mode of inhibition. Together with the low significance for additional equilibrium constants (an order of magnitude higher (weaker) in comparison to competitive inhibition constant), the extension to any of tested mixed-type model is not supported by the fit analysis.

In conclusion, the model in which azaCTZ competes with the substrate for the same binding site provided the most reliable description of the kinetic data. The equilibrium constants for the dissociation of enzyme-inhibitor complex ($K_{I1}$ = 0.092 ± 0.002 μM) is closely similar to Michaelis constant ($K_m$ = 0.042 ± 0.001 μM) which further supports the assumption that azaCTZ binds to the active site correspondingly to the substrate.

The same systematic analysis was performed for kinetics of RLuc8 and its inhibition by azaCTZ (**Supplementary Figure 6-8**). The best fit was provided by mixed-type inhibition model II (**Supplementary Figure 3**) suggesting that azaCTZ binds to the free enzyme ($K_{I1}$ = 16.1 ± 0.1 μM) and competes with the substrate for the active site, but it can also bind to the enzyme-product complex with comparable efficiency ($K_{I2}$ = 19.8 ± 0.1 μM). This inhibition model provides an order of magnitude better statistics (**Supplementary Figure 6b, Supplementary Table 2**) in comparison to any other tested inhibition models. The confidence contour analysis confirmed that all the parameters of mixed-type model II are very well constrained by the kinetic data, with sharp confidence intervals defined by narrow and symmetric confidence contours (**Supplementary Figure 8**). Moreover, the estimates of the turnover number ($k_{cat}$ = 4.69 ± 0.01 $s^{-1}$), Michaelis constant ($K_m$ = 1.61 ± 0.01 μM), and the equilibrium dissociation constants for enzyme-product complex ($K_P$ = 1.41 ± 0.01 μM) provided by Mixed-type inhibition model II are nearly identical to previously reported values of RLuc8 steady-state kinetic parameters.[2,3]

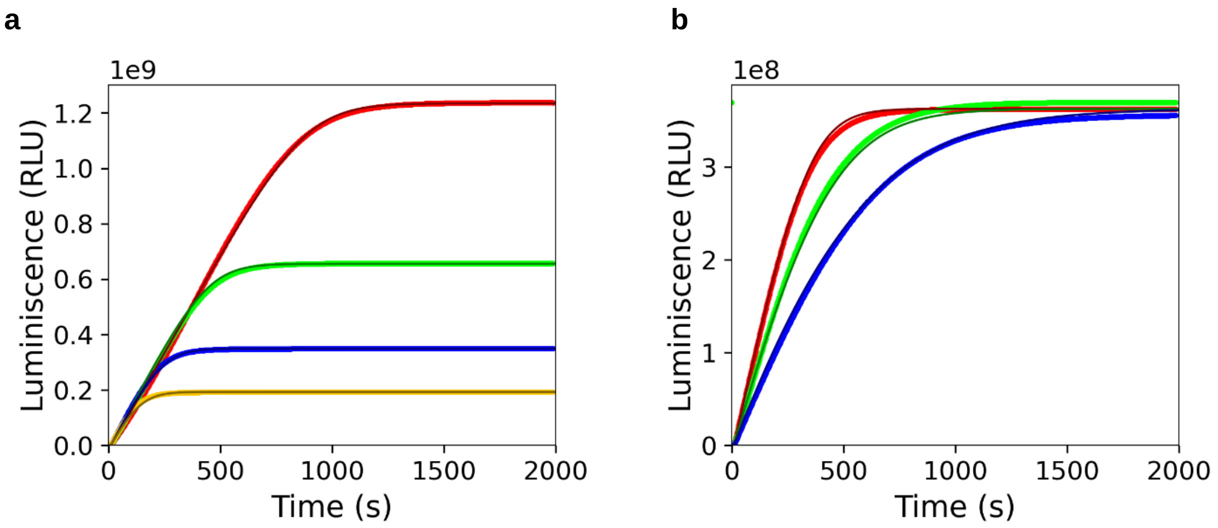


**Supplementary Figure 4.** Analysis of AncFT inhibition by azaCTZ using numerical methods. (a) The reaction progress curves corresponding to cumulative luminescence production in time recorded upon mixing 0.02 μM AncFT with 0.138 μM (yellow), 0.275 μM (blue), 0.55 μM (green) and 1.1 μM coelenterazine (red). (b) The reaction progress curves obtained upon mixing 0.013 μM AncFT with 0.275 μM coelenterazine in the presence of 0 μM (red), 0.3 μM (green) and 0.5 μM azaCTZ (blue). Each trace represents an average of three repetitions. The solid lines represent the best global fit using the competitive model.

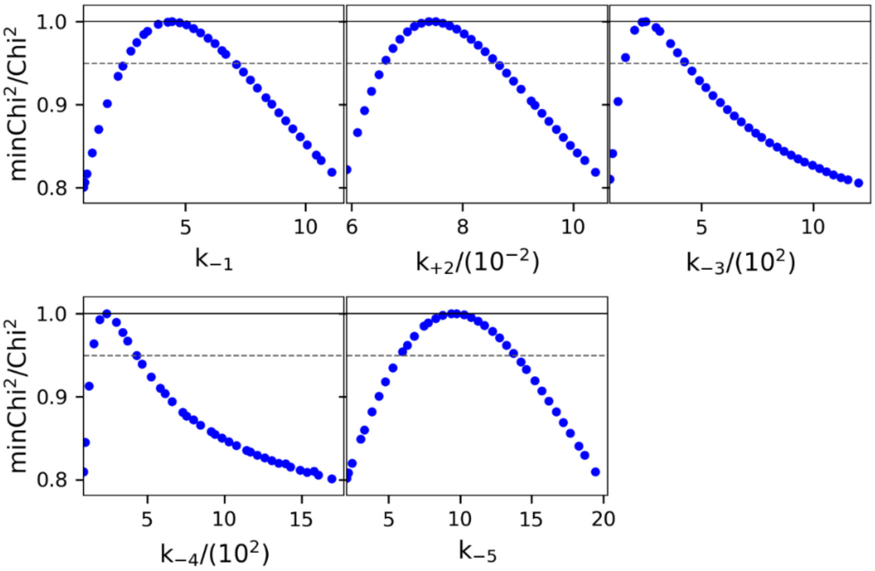


**Supplementary Figure 5.** Confidence contour analysis of parameters obtained for AncFT kinetics using the competitive inhibition model. The figures represent the dependence of the error on the individual fitted parameter while varying all other parameters to achieve the best fit. The dashed line shows the $\chi^2$ threshold (0.95) used to establish confidence intervals reported in **Supplementary Table 1**.

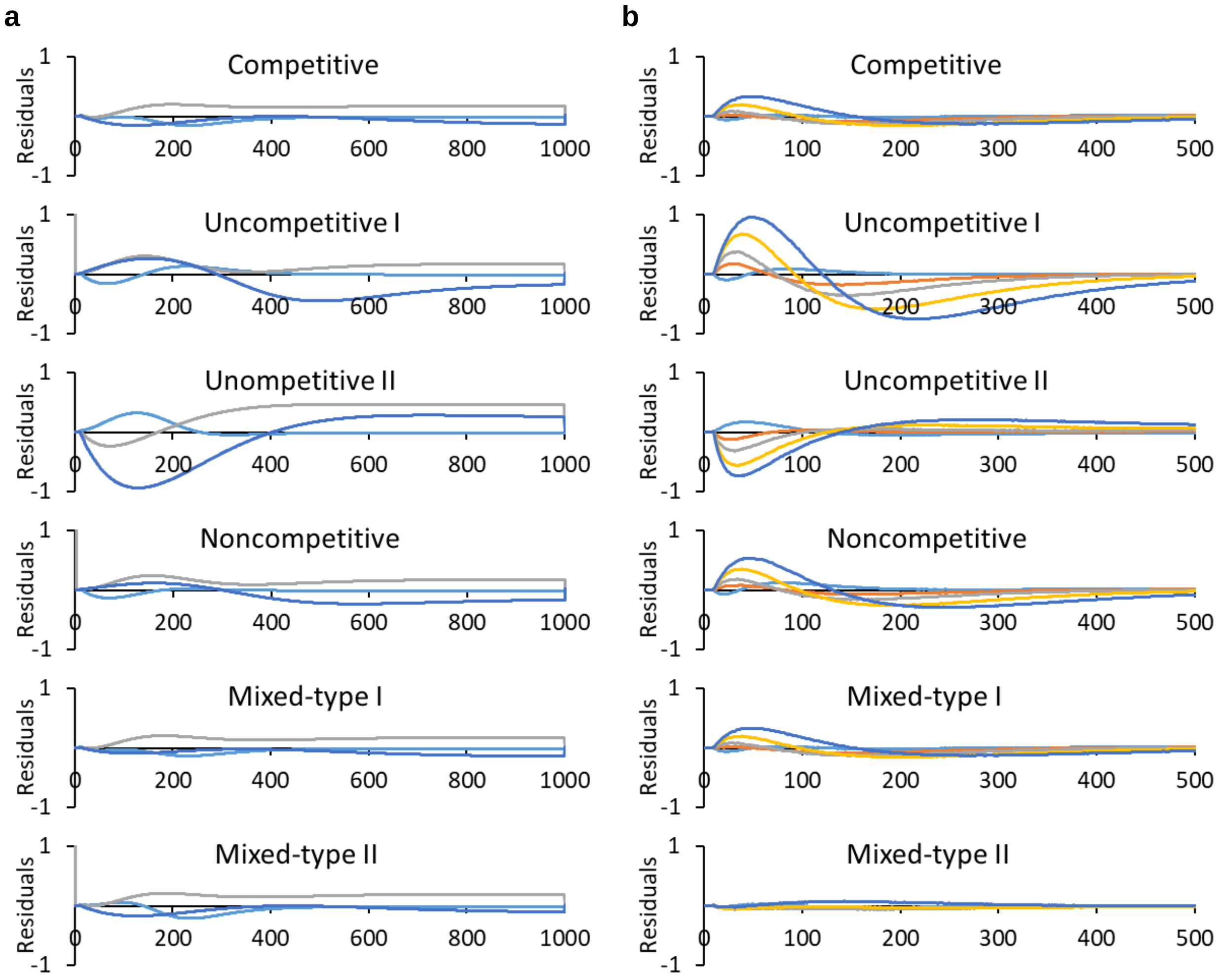


**Supplementary Figure 6.** Comparison of different models for inhibition effect of azaCTZ. To evaluate goodness of fit, the residuals ($y_{observed}$ - $y_{calculated}$) was displayed for all inhibition models fitted to kinetic data recorded with AncFT (a) and RLuc8 (b) in the presence of azaCTZ.

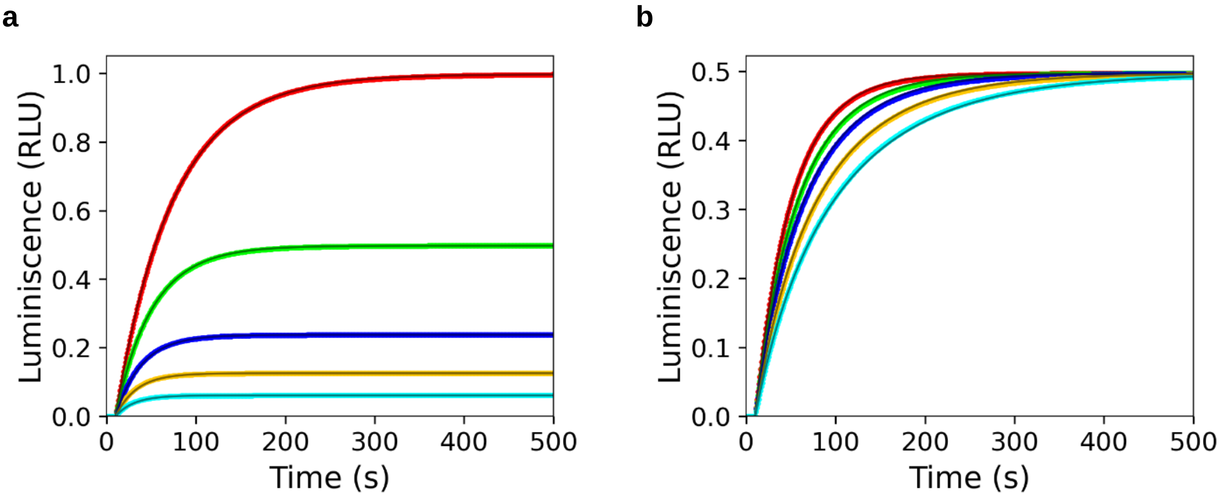


**Supplementary Figure 7.** Analysis of RLuc8 inhibition by azaCTZ using numerical methods. (a) The reaction progress curves corresponding to cumulative luminescence production in time recorded upon mixing 0.02 μM RLuc8 with 0.25 μM (cyan), 0.5 μM (yellow), 1 μM (blue), 2 μM (green) and 4 μM coelenterazine (red). (b) The reaction progress curves obtained upon mixing 0.02 μM Rluc8 with 2 μM coelenterazine in the presence of 0 μM (red), 4.5 μM (green), 9 μM (blue) 18 μM (yellow) and 30 μM (cyan) azaCTZ. Each trace represents an average of three repetitions. The solid lines represent the best global fit using the mixed-type model.

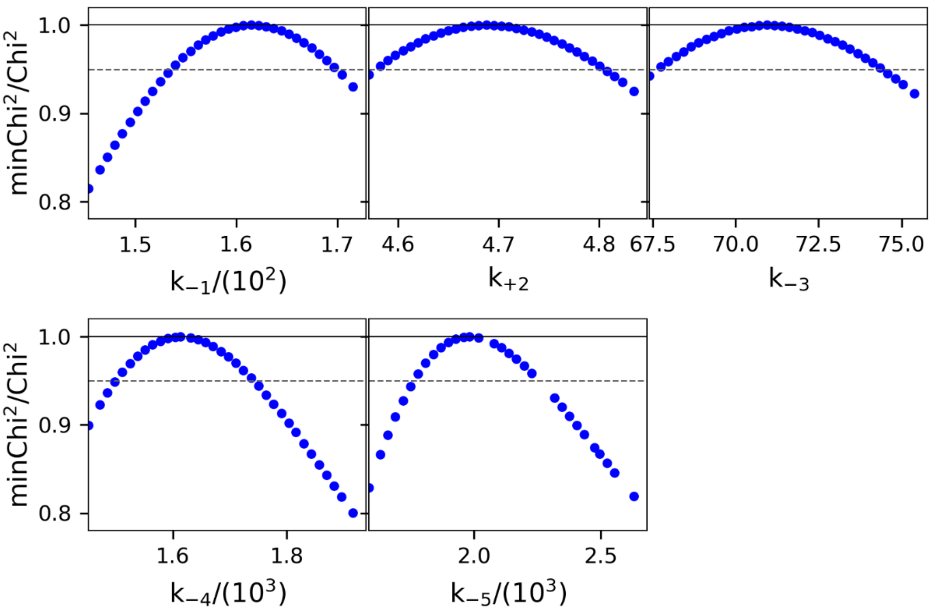


**Supplementary Figure 8.** Confidence contour analysis on parameters obtained for Rluc8 kinetics using the mixed-type inhibition model. The figures represent the dependence of the error on the individual fitted parameter while varying all other parameters to achieve the best fit. The dashed line shows the $\chi^2$ threshold (0.95) used to establish confidence intervals reported in **Supplementary Table 2**.

**Supplementary Table 2.** Kinetic constants obtained by global data analysis for RLuc8. The parameters were derived by fitting kinetic data using numerical integration of rate equations derived for competitive, uncompetitive, non-competitive, and mixed-type inhibition models described in **Supplementary Figure 3**, where $K_m$ is Michaelis constant, $k_{cat}$ is the turnover number, $K_P$ is the equilibrium dissociation constants for enzyme-product complex, $K_{SI}$ is equilibrium constant describing substrate inhibition, $K_{I1}$ and $K_{I2}$ is an inhibition equilibrium constant describing binding of the inhibitor to the free enzyme and enzyme-substrate/product complex, respectively. The kinetic parameters are reported as values obtained from the best fitted ± standard errors (S.E.). In addition, the lower and upper confidence limits at $\chi^2$ threshold 0.95 are summarized in the brackets for each parameter of the best fit.

| | Competitive | Uncompetitive I | Uncompetitive II | Noncompetitive | Mixed-type I | **Mixed-type II** |
|---|---|---|---|---|---|---|
| **$\chi^2$ x $10^6$** | 3.8 | 39.5 | 12.6 | 2.9 | 3.8 | **0.75** |
| **$K_m$ (μM)** | 1.66 ±0.01 | 1.69 ±0.03 | 1.81 ±0.02 | 1.56 ±0.01 | 1.66 ±0.01 | **1.61 ±0.01**<br>(1.54; 1.70) |
| **$k_{cat}$ ($s^{-1}$)** | 4.88 ±0.01 | 5.05 ±0.05 | 4.67 ±0.02 | 4.81 ±0.01 | 4.88 ±0.01 | **4.69 ±0.01**<br>(4.58; 4.8) |
| **$K_P$ (μM)** | 1.33 ±0.01 | 1.19 ±0.02 | 1.74 ±0.01 | 1.24 ±0.01 | 1.33 ±0.01 | **1.41 ±0.01**<br>(1.34; 1.48) |
| **$K_{I1}$ (μM)** | 10.5 ±0.1 | n.a. | n.a. | 24.1 ±0.1 | 10.5 ±0.1 | **16.1 ±0.1**<br>(15.1; 17.4) |
| **$K_{I2}$ (μM)** | n.a. | 7.1 ±0.1 | 7.1 ±0.1 | n.a. | > 100 | **19.8 ±0.1**<br>(17.8; 22.3) |
| **$K_{SI}$ (μM)** | n.d. | n.d. | n.d. | n.d. | n.d. | n.d. |

n.a. not applied

n.d. not determined (the parameter did not provide significant value)

**a** **b**

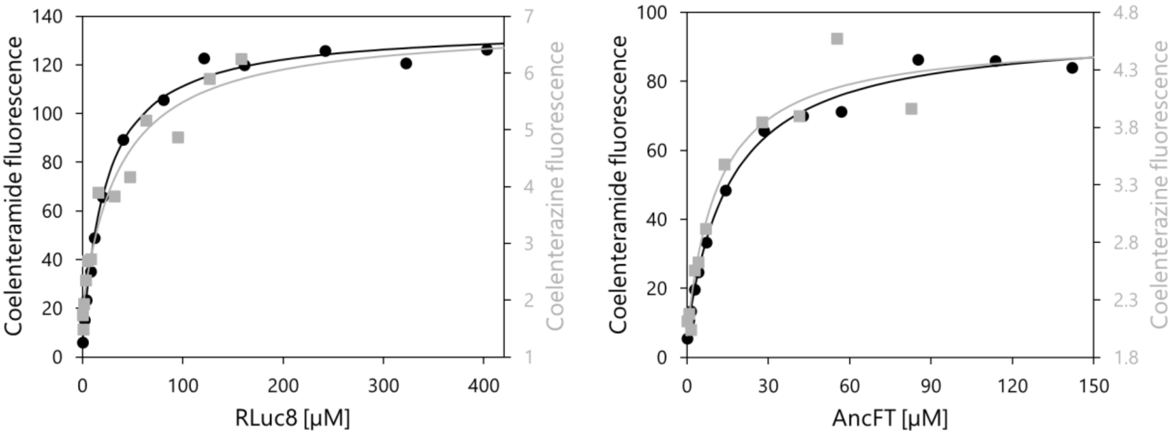


**c**

| Luciferase enzyme | Dissociation constant $K_d$ [μM] | | $K_d$ ratio |
|---|---|---|---|
| | Coelenterazine substrate | Coelenteramide product | substrate/product |
| RLuc8 | 37 ± 15 | 22 ± 2 | 1.7 ± 0.7 |
| AncFT | 12 ± 5 | 16 ± 2 | 0.7 ± 0.3 |

**Supplementary Figure 9.** Anaerobic equilibrium binding experiments. The figure displays concentration dependence of CTZ/CEI fluorescence peak maxima with increasing concentration of RLuc8 (**a**) and AncFT (**b**) due to the binding by the enzymes. The data were collected under anaerobic conditions in 100 mM potassium phosphate buffer pH 7.5 at 30 °C. Solid lines represent the best fit according to the hyperbolic relationship. (**c**) The derived dissociation constants $K_d$ for each enzyme–ligand pair.

**Supplementary Table 3.** Crystallographic data collection and refinement statistics of AncFT complexes.

| | AncFT/azaCTZ | AncFT/CEI |
|---|---|---|
| **Data collection** | | |
| Wavelength (Å) | 1 | 1 |
| Space group | $P12_11$ | $P12_11$ |
| Cell dimensions | | |
| a, b, c (Å) | 51.284, 87.549, 102.693 | 49.224, 83.332, 102.017 |
| α, β, γ (°) | 90, 93.462, 90 | 90, 91.885, 90 |
| Resolution (Å) | 44.73 - 2.052 (2.125 - 2.052) | 44.89 - 2.251 (2.332 - 2.251) |
| Total reflections | 387,158 (37,596) | 259,714 (24,256) |
| Unique reflections | 56,417 (5,534) | 39,019 (3,794) |
| Rmerge | 0.1258 (1.525) | 0.07774 (0.3767) |
| I/σI | 10.66 (1.29) | 18.90 (5.44) |
| Completeness (%) | 99.34 (98.03) | 99.60 (98.80) |
| Multiplicity | 6.9 (6.8) | 6.7 (6.4) |
| CC (1/2) | 0.998 (0.645) | 0.998 (0.958) |
| Wilson B-factor | 33.65 | 25.67 |
| **Refinement** | | |
| Resolution (Å) | 44.73 - 2.052 (2.125 - 2.052) | 44.89 - 2.251 (2.332 - 2.251) |
| No. reflections | 56,417 (5,525) | 39,022 (3,786) |
| Rwork / Rfree (%) | 19.24 / 24.70 | 17.15 / 24.08 |
| No. atoms | | |
| Protein | 7,172 | 7,261 |
| Ligand | 96 | 93 |
| Water | 303 | 275 |
| B-factors | 37.43 | 29.97 |
| Protein | 37.50 | 29.90 |
| Ligand | 32.81 | 36.39 |
| Water | 37.21 | 29.67 |
| R.m.s. deviations | | |
| Bond lengths (Å) | 0.008 | 0.008 |
| Bond angles (°) | 0.91 | 0.94 |
| Ramachandran favored (%) | 94.52 | 95.12 |
| Ramachandran allowed (%) | 5.48 | 4.54 |
| Ramachandran outliers (%) | 0.00 | 0.34 |
| PDB ID code | 7QXR | 7QXQ |

Values in parentheses are for the highest-resolution shell.

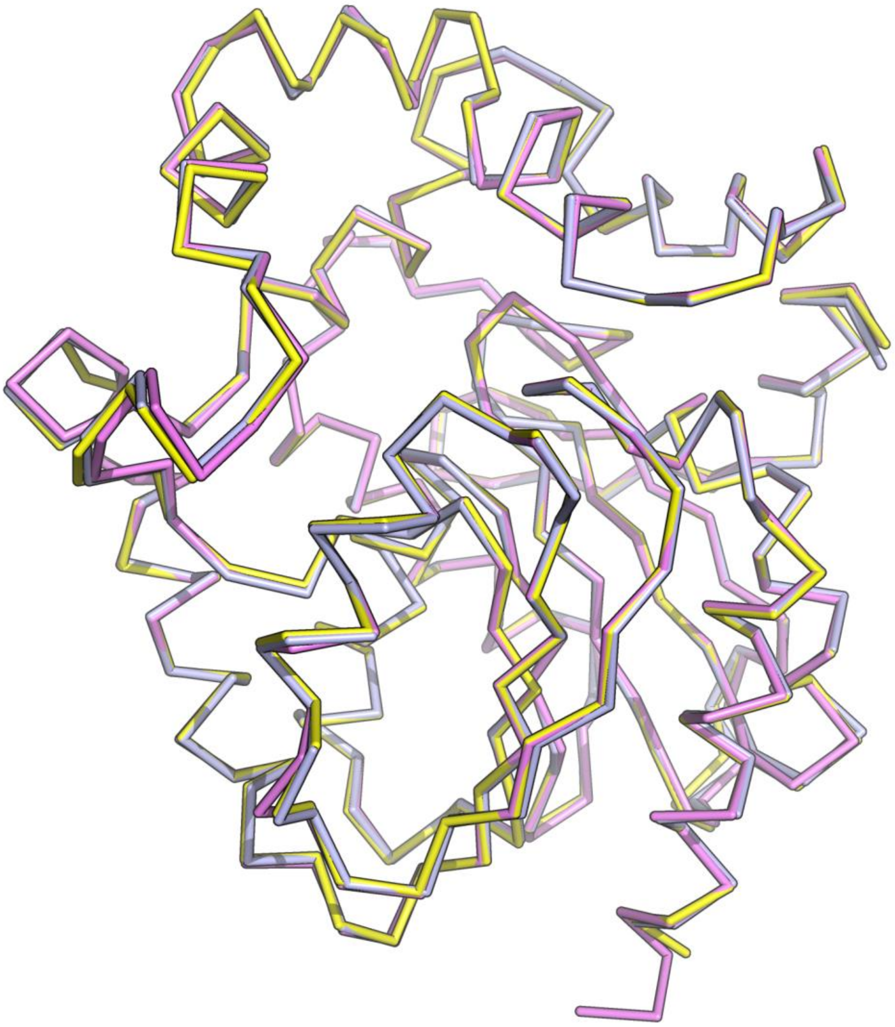

**Supplementary Figure 10.** Comparison between apo-AncFT (yellow), azaCTZ-bound AncFT (violet) and CEI-bound AncFT (light-blue) structures. The root-mean-square deviation (RMSD) values on Cα atoms range from 0.1 to 0.3 Å.

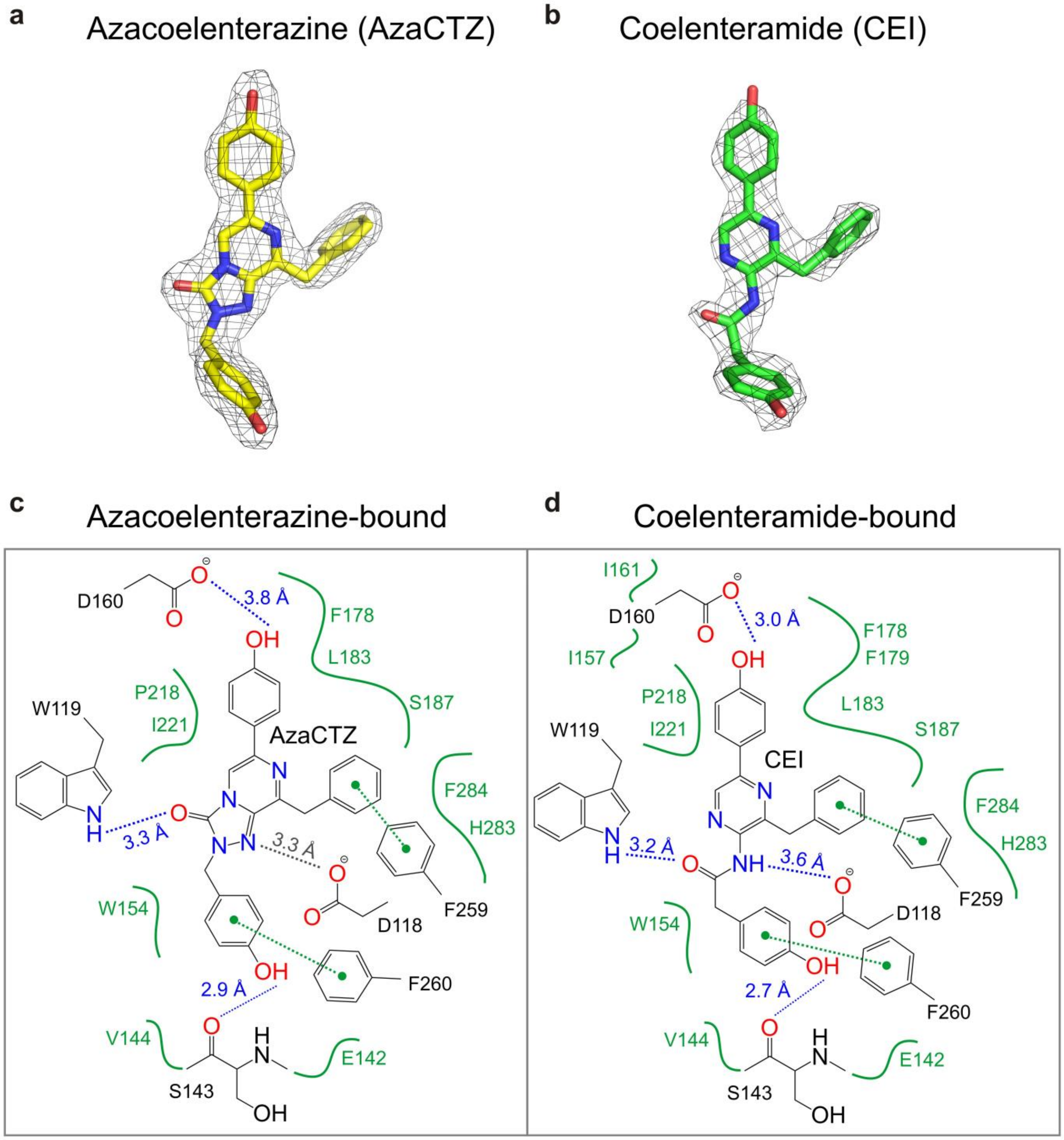


**Supplementary Figure 11.** Structures and interactions of azaCTZ and CEI bound to AncFT enzyme. (**a**,**b**) Simulated annealing omit electron density maps contoured at 1σ for azaCTZ molecule bound in AncFT/azaCTZ complex (**a**; PDB ID: 7QXR), and CEI molecule bound in AncFT/CEI complex (**b**; PDB ID: 7QXQ). (**c**,**d**) Two-dimensional representations of AncFT amino acid residues interacting with azaCTZ (**c**) and CEI (**d**). Hydrogen bonds are depicted as blue dotted lines, π-π interactions as green dotted lines, amino acids within 4 Å around ligands creating a hydrophobic cavity are shown in green.

**Supplementary Table 4.** Crystallographic data collection and refinement statistics of RLuc8 complexes.

| | RLuc8-D162A/azaCTZ | RLuc8-D162A/CEI | RLuc8-D120A/CNM |
|---|---|---|---|
| **Data collection** | | | |
| Wavelength (Å) | 1 | 1 | 1 |
| Space group | $P12_11$ | $P2_122_1$ | $P12_11$ |
| Cell dimensions | | | |
| a, b, c (Å) | 51.569, 84.057, 77.437 | 56.445, 74.166, 80.346 | 49.315, 139.972, 50.623 |
| α, β, γ (°) | 90, 90.72, 90 | 90, 90, 90 | 90, 112.555, 90 |
| Resolution (Å) | 38.05 - 1.601 (1.658 - 1.601) | 46.19 - 1.5 (1.554 - 1.5) | 46.75 - 1.451 (1.503 - 1.451) |
| Total reflections | 585,208 (55,569) | 717,149 (67,306) | 760,919 (76,615) |
| Unique reflections | 86,713 (8,547) | 54,640 (5,336) | 111,422 (11,106) |
| Rmerge | 0.05303 (0.9663) | 0.08651 (1.825) | 0.0454 (0.7565) |
| I/σI | 21.21 (1.87) | 19.98 (1.42) | 22.41 (2.44) |
| Completeness (%) | 99.79 (98.79) | 99.89 (99.11) | 99.97 (99.96) |
| Multiplicity | 6.7 (6.5) | 13.1 (12.6) | 6.8 (6.9) |
| CC (1/2) | 1 (0.714) | 1 (0.634) | 1 (0.836) |
| Wilson B-factor | 21.41 | 18.20 | 17.13 |
| **Refinement** | | | |
| Resolution (Å) | 38.05 - 1.601 (1.658 - 1.601) | 46.19 - 1.5 (1.554 - 1.5) | 46.75 - 1.451 (1.503 - 1.451) |
| No. reflections | 86,713 (8,547) | 54,640 (5,333) | 111,421 (11,106) |
| Rwork / Rfree (%) | 17.61 / 21.10 | 17.05 / 18.61 | 16.32 / 18.12 |
| No. atoms | | | |
| Protein | 5,123 | 2,562 | 5,013 |
| Ligand | 140 | 55 | 49 |
| Water | 692 | 345 | 463 |
| B-factors | 28.83 | 24.01 | 22.91 |
| Protein | 27.53 | 22.47 | 22.13 |
| Ligand | 29.93 | 36.47 | 34.51 |
| Water | 38.21 | 33.44 | 30.11 |
| R.m.s. deviations | | | |
| Bond lengths (Å) | 0.006 | 0.006 | 0.005 |
| Bond angles (°) | 0.86 | 0.93 | 0.82 |
| Ramachandran favored (%) | 96.10 | 96.74 | 96.65 |
| Ramachandran allowed (%) | 3.90 | 3.26 | 3.35 |
| Ramachandran outliers (%) | 0.00 | 0.00 | 0.00 |
| PDB ID code | 7OMD | 7OMR | 7OMO |

Values in parentheses are for the highest-resolution shell

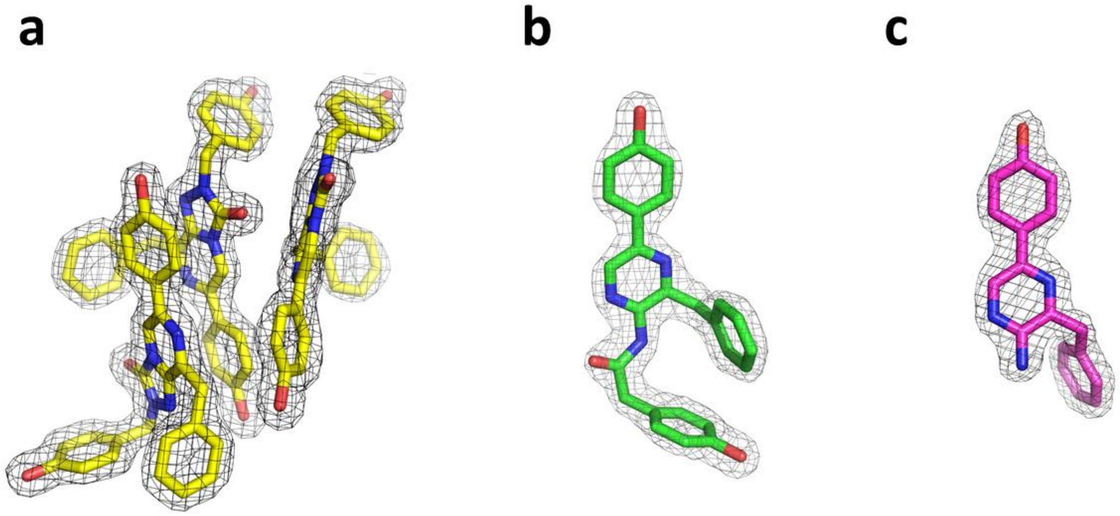

**Supplementary Figure 12.** Crystal structures of azaCTZ, CEI, and CNM bound to RLuc8 mutants. Simulated annealing omit electron density maps contoured at 1σ for azaCTZ molecules (yellow) bound in RLuc8-D162A/azaCTZ complex (**a**; PDB ID: 7OMD), CEI molecule (green) bound in RLuc8-D162A/azaCTZ complex (**b**; PDB ID: 7OMR) and CNM molecule (violet) bound in RLuc8-D120A/CNM complex (**c**; PDB ID: 7OMO). Note that in the RLuc8-D162A/azaCTZ complex, there are up to three azaCTZ molecules (yellow) found in the enzymatic pocket.

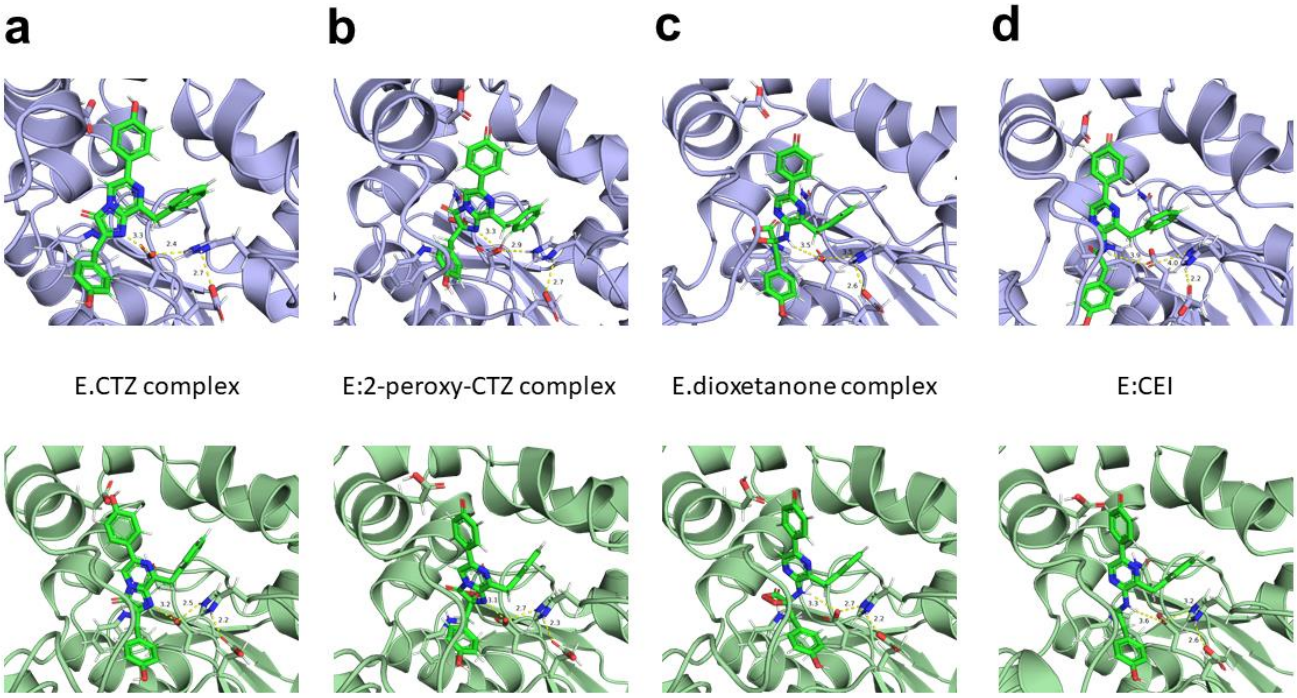


**Supplementary Figure 13.** Representative molecular dynamics simulation snapshots of enzyme-ligand complexes for AncFT (light-blue; upper panels) and RLuc8 (palegreen; bottom panels). (**a**) E.CTZ complexes, (**b**) E.2-peroxy-CTZ complexes, (**c**) E.dioxetanone intermediate complexes, and (**d**) E.CEI complexes. Hydrogen bonds are shown as yellow dashed lines, and the corresponding distances between the atoms and/or chemical groups are given in Å.

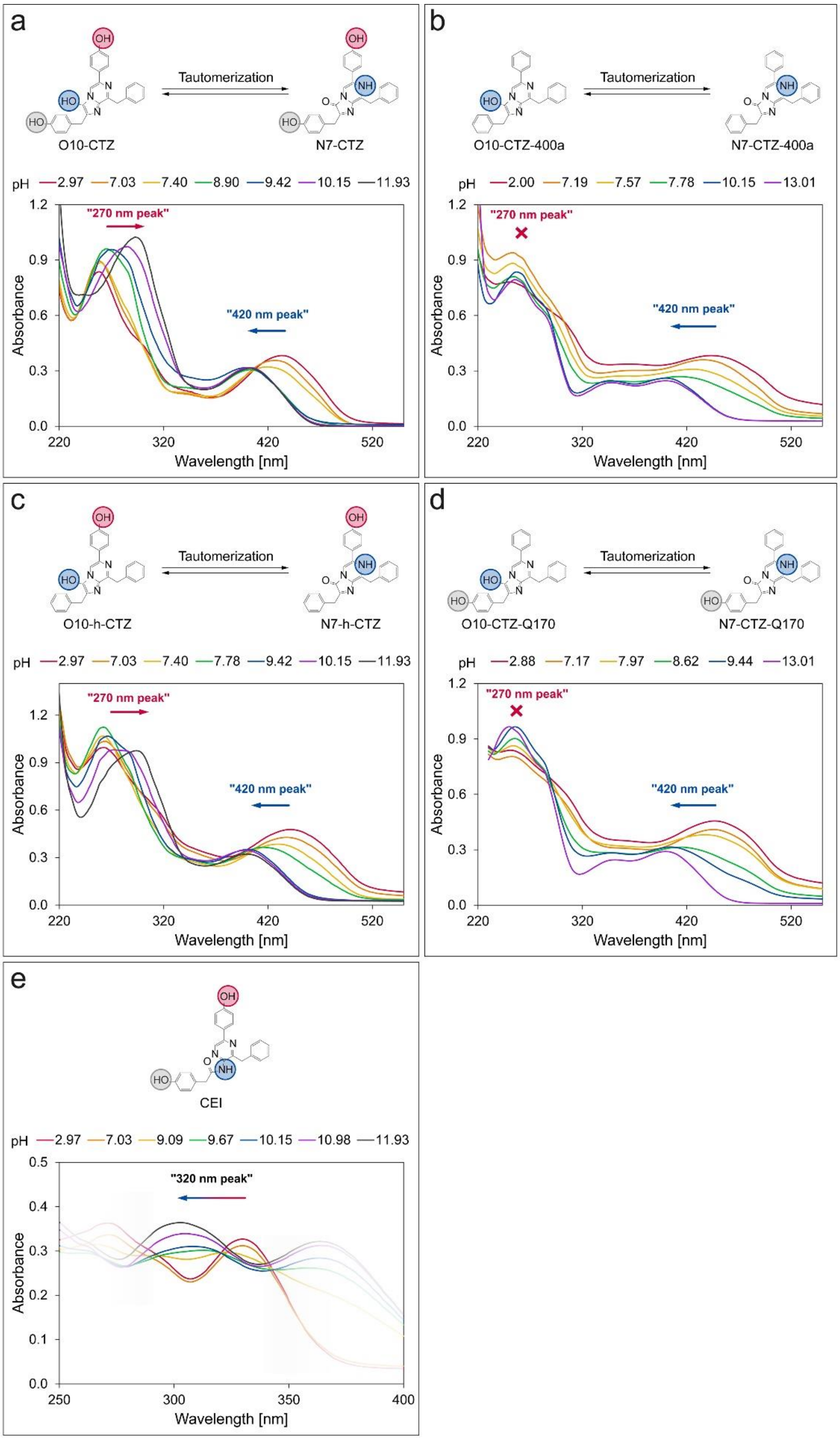


**Supplementary Figure 14.** Comparison of pH-dependent absorbance spectra of the CTZ substrate, its analogues, and the CEI product. The spectra were collected for native CTZ (**a**), CTZ-400a (**b**), h-CTZ (**c**), CTZ-Q170[4] (**d**), and CEI (**e**). The peaks exhibit a clear shift with increasing pH due to the (de)protonation of ionizable groups highlighted with red and blue circles. The corresponding peaks are marked with arrows of the same color.

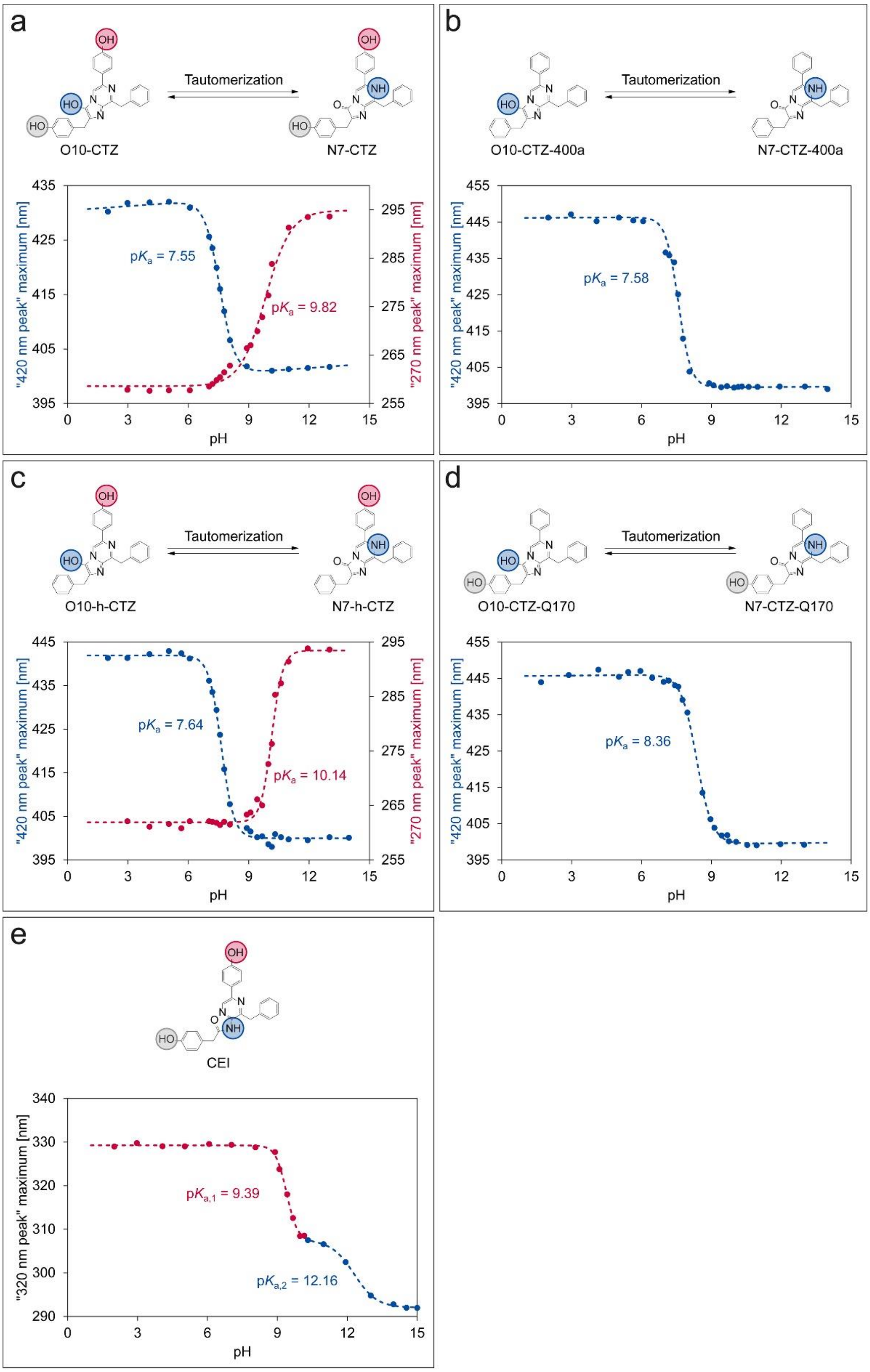


**Supplementary Figure 15.** Analysis of the CTZ substrate, its analogues, and the CEI product absorbance peaks position maxima at various pH. The spectra were collected for native CTZ (**a**), CTZ-400a (**b**), h-CTZ (**c**), CTZ-Q170[4] (**d**), and CEI (**e**). The dependence yields typical sigmoid pH-titration curves, allowing to determine p$K_a$ values of individual ionizable groups highlighted with red and blue circles. The corresponding sigmoid transitions are marked with the same color.

**Supplementary Table 5.** Comparison of p$K_a$ values of (de)protonable groups of CTZ and its analogues and CEI. The results are provided as best fit values ± standard errors calculated from nonlinear sigmoidal regression. n.a. = not applicable.

| **Substrate/analogue molecule** | **p$K_a$ ± standard error** | |
|---|---|---|
| | **O10/N7 –OH/–NH group** | **6-(p-HOPh) –OH group** |
| Coelenterazine | 7.55 ± 0.01 | 9.82 ± 0.07 |
| Coelenterazine-400a | 7.58 ± 0.03 | n.a. |
| h-Coelenterazine | 7.64 ± 0.02 | 10.14 ± 0.03 |
| Coelenterazine-Q170 | 8.36 ± 0.02 | n.a. |
| **Product molecule** | **p$K_a$ ± standard error** | |
| | **Amide –NH group** | **5-(p-HOPh) –OH group** |
| Coelenteramide | 12.16 ± 0.08 | 9.40 ± 0.02 |

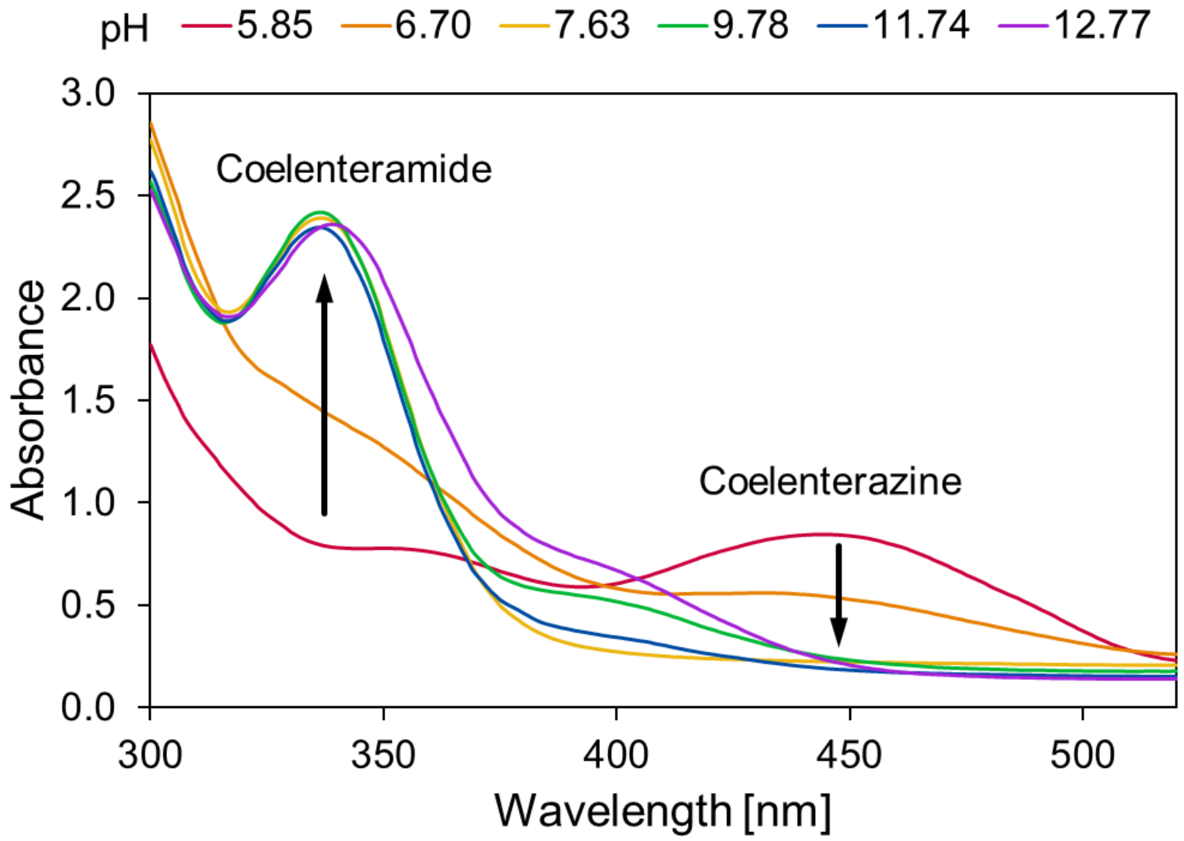


**Supplementary Figure 16.** Absorbance spectra of the fully reacted mixture after mixing the CTZ substrate with DMSO at various pH. The spectra were collected after the chemiluminescence signal, due to the CTZ autooxidation, dropped back to the background level to assure that the reaction fully completed. At low pH, CTZ remains in the mixture and no CEI is formed.

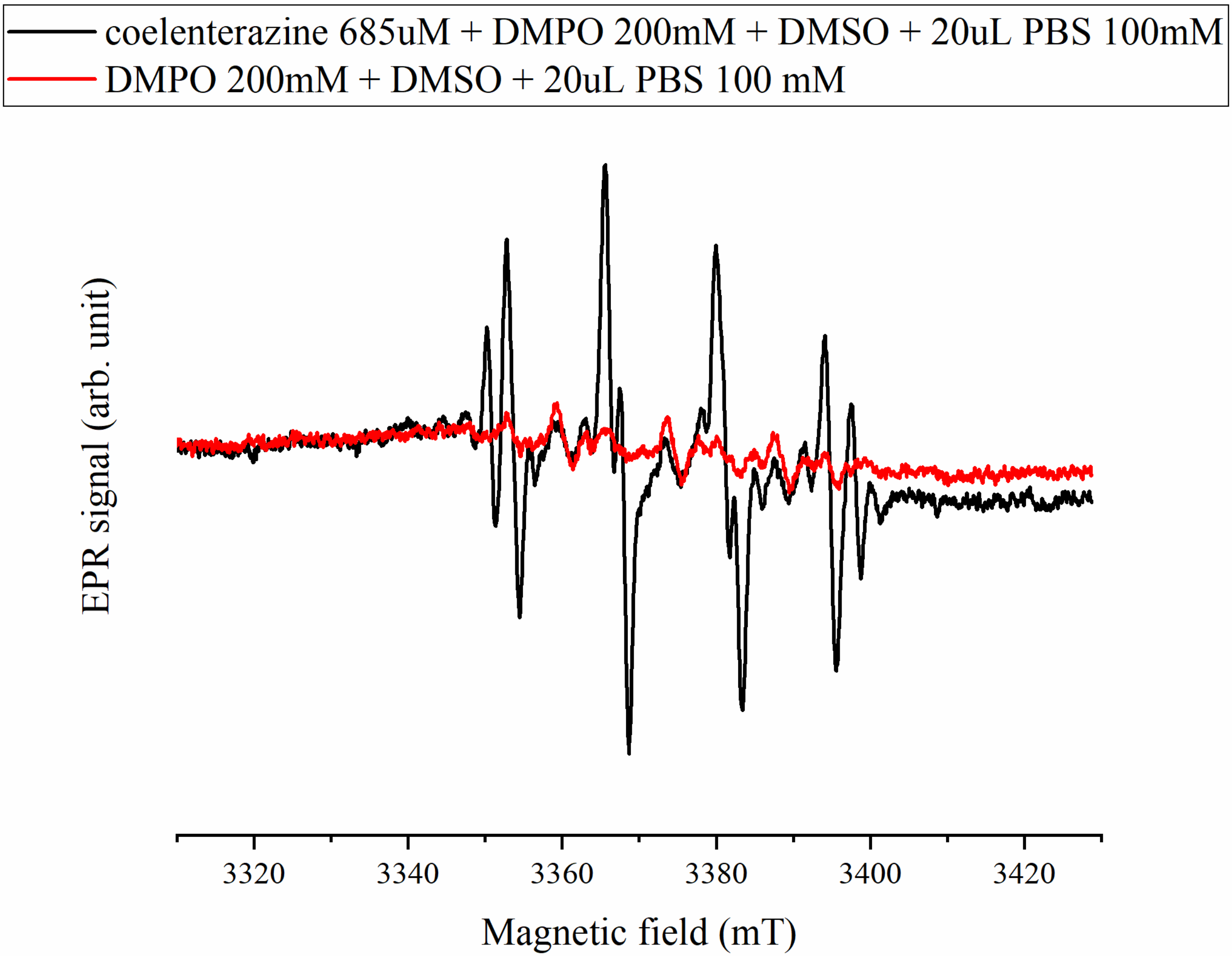


**Supplementary Figure 17.** Original experiment with CTZ and control measurement without CTZ both accumulated 36 times and averaged. The simulated EPR spectrum in the main text was obtained as the difference between these two spectra.

**Supplementary Table 6.** The individual contributions of four components C1-4 together with the hyperfine coupling constants and the g-factor for each component.

| | ***g*** | **$a_N$ (G)** | **$a_H^{\beta}$ (G)** | **$a_H^{\gamma}$ (G)** | **rel. conc. (%)** | **assigned to** |
|---|---|---|---|---|---|---|
| **C1** | 2.0072 | 14.0 | 11.4 | 1.30 | 28 | DMPO-OOH |
| **C2** | 2.0071 | 13.3 | 10.9 | 1.87 | 19 | DMPO-OOH |
| **C3** | 2.0070 | 14.2 | 12.7 | - | 16 | DMPO-OH |
| **C4** | 2.0070 | 15.2 | 17.0 | - | 37 | Unidentified carbon-based radical |

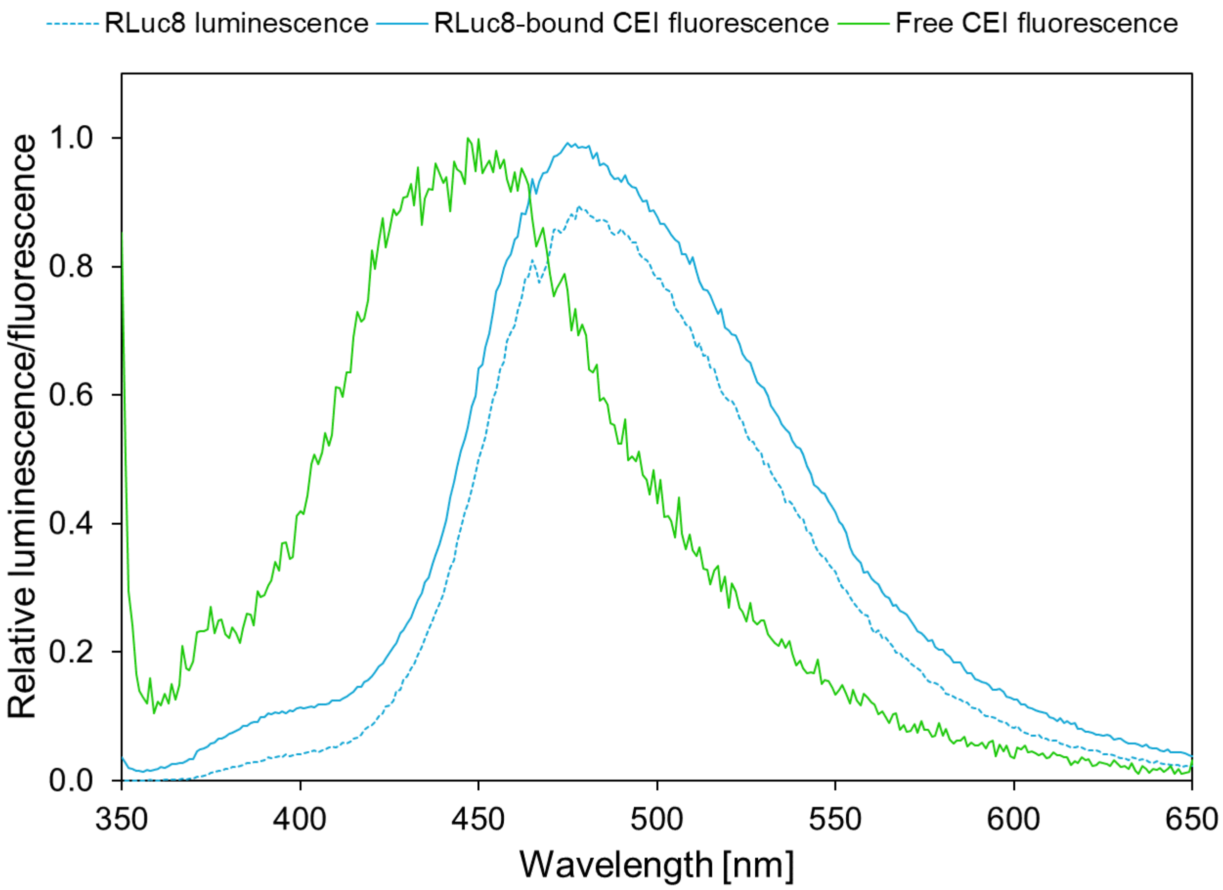


**Supplementary Figure 18.** Comparison of RLuc8 luminescence emission spectrum with fluorescence emission spectra of RLuc8-bound and free (bulk) form of CEI. The data were collected in 100 mM potassium phosphate buffer pH 7.5 at ambient temperature. The luminescence spectrum corresponds to the enzyme-bound fluorescence spectrum, showing that light emission from the CEI molecule during enzymatic conversion originates from the enzyme active site before the CEI product is released.

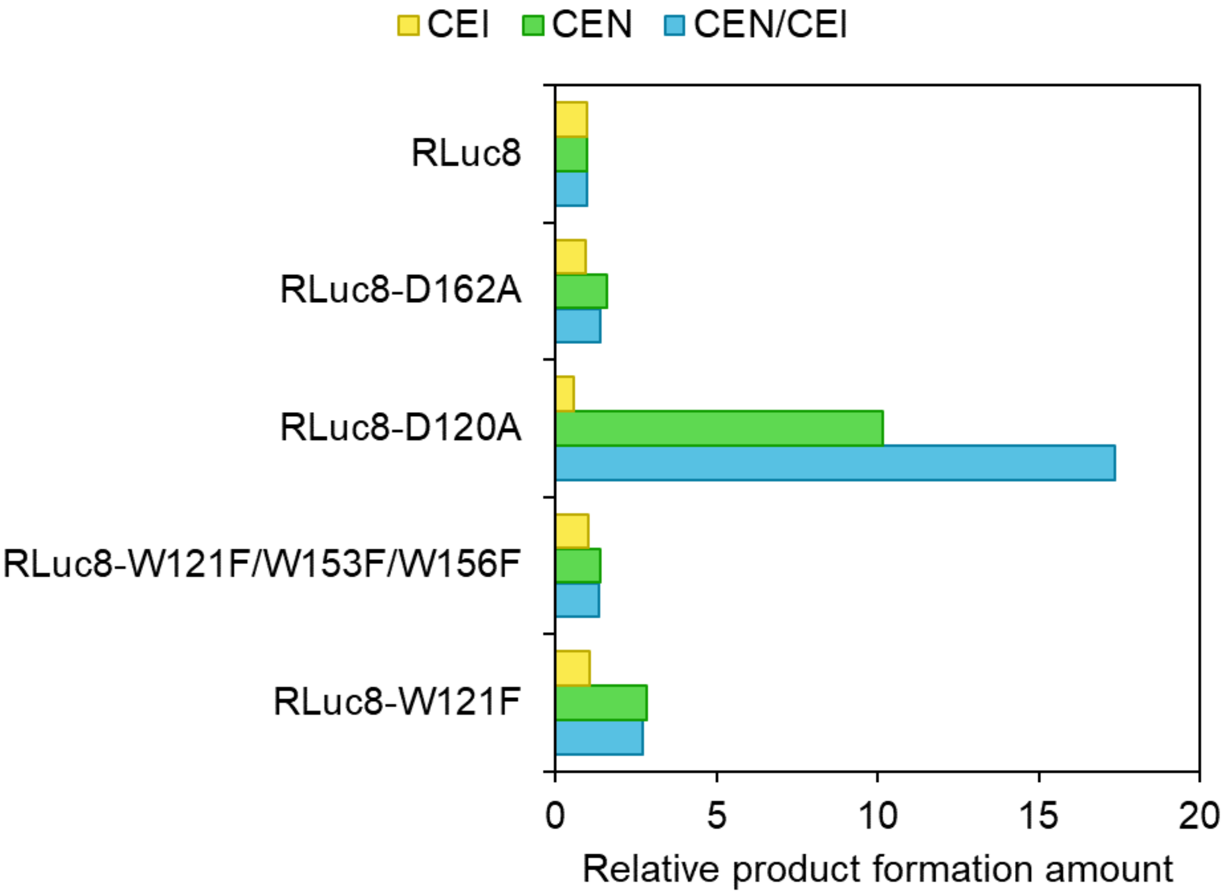


**Supplementary Figure 19.** Comparison of variability in total production of the main product CEI and the side product CEN by RLuc8 mutant variants after mixing with the same amount of the CTZ substrate. The experiment was performed in 100 mM potassium phosphate buffer pH 7.5 at 37 °C. The amount of formed product is relativized by setting the amount of produced CEI and CEN by RLuc8 to 1.0. All the variants are consistent in CEI and CEN production except for RLuc8-D120A, producing 17-fold more CEN over CEI than RLuc8.

**Supplementary Table 7.** A list of PCR primers used for gene amplifications and mutagenesis.

| Protein Variant | Template | Fw primer sequence (5' - 3') | Rv primer sequence (5' - 3') | Method |
|---|---|---|---|---|
| AncFT-D118A | *AncFT* | AAAGTTACCATTGTTTGTC ATGCCTGGGGTAGCGGTC TGGGTTTT | GCTAGTTATTGCTCAGCG G | MegaPrimer |
| AncFT-H283F | *AncFT* | TAATACGACTCACTATAGG G | CGGTGAATCTTCTTGCAG AAAAAACAGACCTTTAACG GTAACGGT | MegaPrimer |
| AncFT-W119F | *AncFT* | GTTACCATTGTTTGTCATG ATTTTGGTAGCGGTCTGGG TTTTCAT | GCTAGTTATTGCTCAGCG G | MegaPrimer |
| AncFT-N51A | *AncFT* | ACCGTGATTTTTCTGCATG GTGCCCCGACCAGCAGCT ATCTGTGG | GCTAGTTATTGCTCAGCG G | MegaPrimer |
| AncFT-D160A | *AncFT* | GAATGGCCTGATATCGAAG AAGCCATTGCCCTGATTAA AAGCGAA | GCTAGTTATTGCTCAGCG G | MegaPrimer |
| RLuc8-D120A | *RLuc8* | ATCTTTGTTGGTCATGCGT GGGGTGCAGCACTG | CAGTGCTGCACCCCACGC ATGACCAACAAAGAT | QuickChange |
| RLuc8-H285F | *RLuc8* | TAATACGACTCACTATAGG G | CGGTGCATCCTCTTGCAG AAAAAACAGACCTTTCACT TTCACAAA | MegaPrimer |
| RLuc8-E144Q | *RLuc8* | GCCATTGTTCACATGCAGA GCGTTGTGGATGTT | AACATCCACAACGCTCTG CATGTGAACAATGGC | QuickChange |
| RLuc8-W121F | *RLuc8* | TAATACGACTCACTATAGG G | TGAAATGCCAGTGCTGCA CCAAAATCATGACCAACAA AGATGA | MegaPrimer |
| RLuc8-N53A/W121F | *RLuc8* | CCGTGATTTTTCTGCATGG TGCGGCAACCAGCAGCTA TCTGTG | GAAATGCCAGTGCTGCAC CAAAATCATGACCAACAAA GATGA | MegaPrimer |
| RLuc8-D162A | *RLuc8* | CCTGATATCGAAGAAGCGA TTGCCCTGATTAAA | TTTAATCAGGGCAATCGCT TCTTCGATATCAGG | QuickChange |
| RLuc8-D148A/E151A/S152A | *RLuc8* | GTTCACATGGAAAGCGTTG TGGCTGTTATTGCTGCTTG GGATGAATGGCCTGATATC | GCTAGTTATTGCTCAGCG G | MegaPrimer |
| RLuc8-D154A/E155A | *RLuc8* | GTGGATGTTATTGAAAGCT GGGCTGCTTGGCCTGATA TCGAAGAAGAT | GCTAGTTATTGCTCAGCG G | MegaPrimer |
| RLuc8-D158A/E160A/E161A | *RLuc8* | GAAAGCTGGGATGAATGG CCTGCTATCGCTGCTGATA TTGCCCTGATTAAAAGC | GCTAGTTATTGCTCAGCG G | MegaPrimer |
| RLuc8-W153F/W156F | *RLuc8* | TAATACGACTCACTATAGG G | ATATCTTCTTCGATATCAG GAAATTCATCAAAGCTTTC AATAACATCCACAA | MegaPrimer |
| RLuc8-W153A/W156A | *RLuc8* | TAATACGACTCACTATAGG G | ATATCTTCTTCGATATCAG GGGCTTCATCAGCGCTTT CAATAACATCCACAA | MegaPrimer |
| RLuc8-W156A | *RLuc8* | GTTATTGAAAGCTGGGATG AAGCCCCTGATATCGAAGA AGATATT | GCTAGTTATTGCTCAGCG G | MegaPrimer |
| RLuc8-F262A | *RLuc8* | TAATACGACTCACTATAGG G | ACCCTCAACAATTGCATTG CTGGCAAAACCCGGATCG CTTTCGAT | MegaPrimer |
| RLuc8-W156A/F262A | *RLuc8* | GTTATTGAAAGCTGGGATG AAGCCCCTGATATCGAAGA AGATATT | ACCCTCAACAATTGCATTG CTGGCAAAACCCGGATCG CTTTCGAT | MegaPrimer |

### NMR spectra

3-Benzyl-5-(4-(benzyloxy)phenyl)-2-hydrazinylpyrazine (**2**)

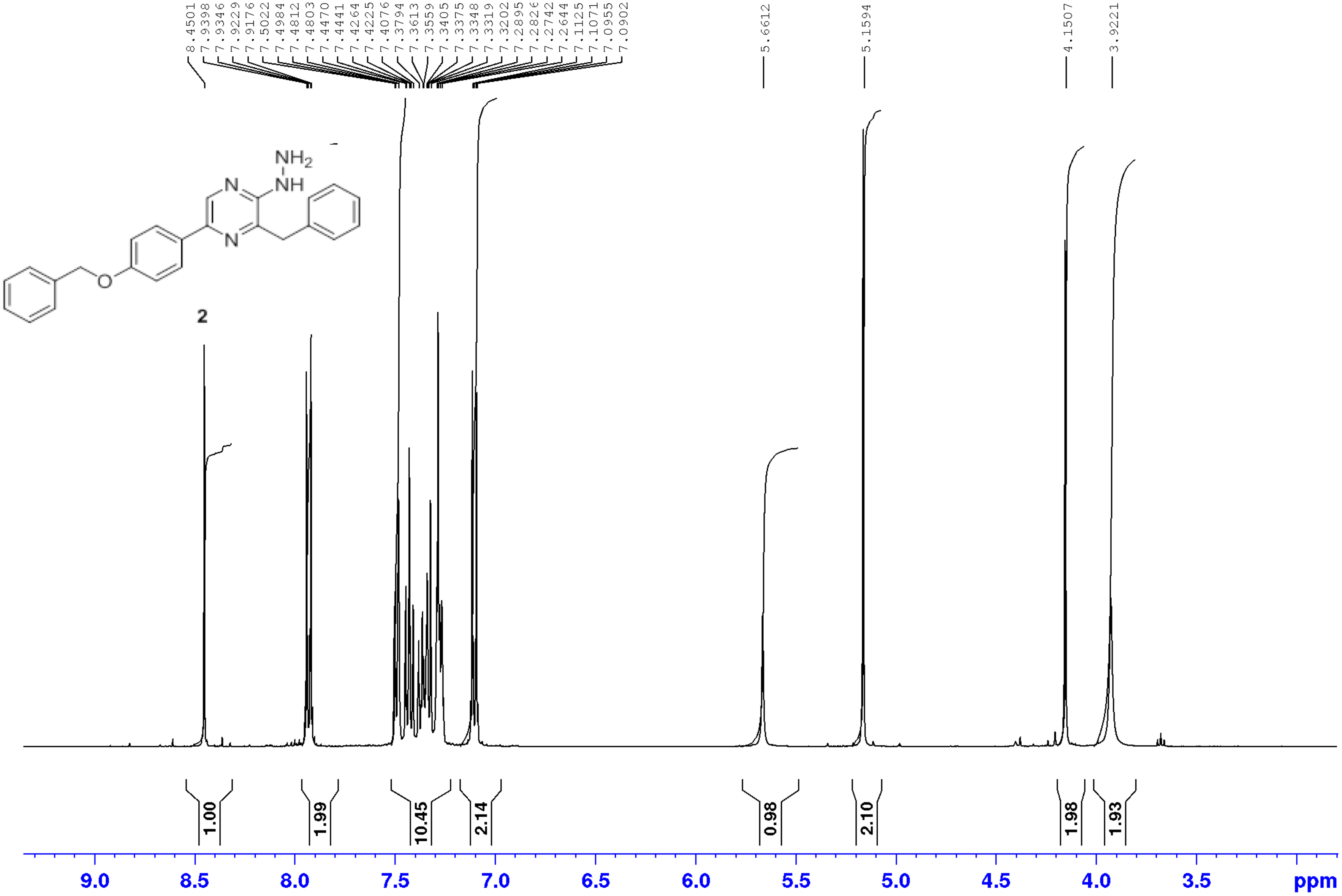

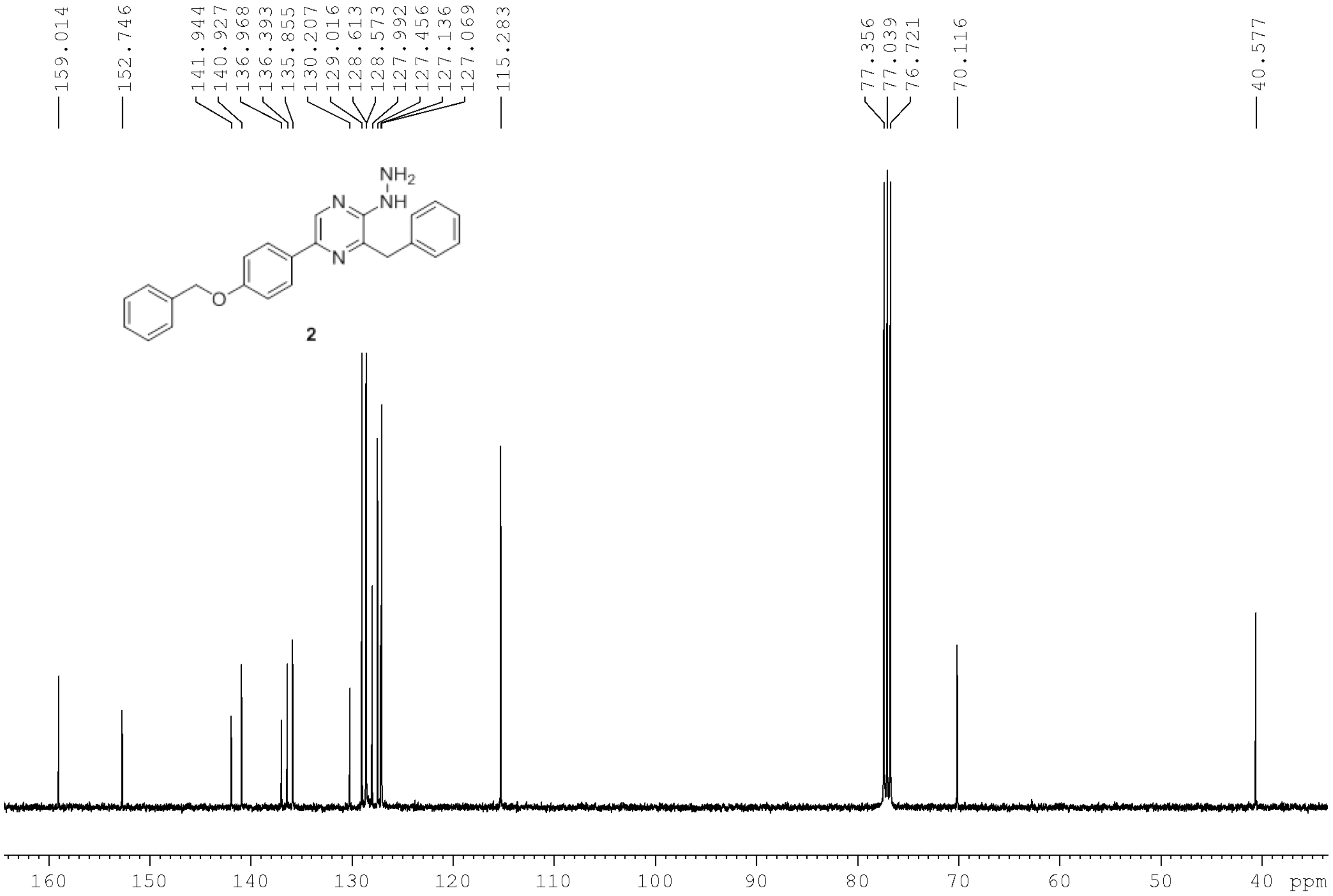
159.014
152.746
141.944
140.927
136.968
136.393
135.855
130.207
129.016
128.613
128.573
127.992
127.456
127.136
127.069
115.283
77.356
77.039
76.721
70.116
40.577
NH2
NH
N
N
O
2
160
150
140
130
120
110
100
90
80
70
60
50
40 ppm

8-Benzyl-2-(4-(benzyloxy)benzyl)-6-(4-(benzyloxy)phenyl)-[1,2,4]triazolo[4,3-*a*]pyrazin-3(2*H*)-one (**5**)

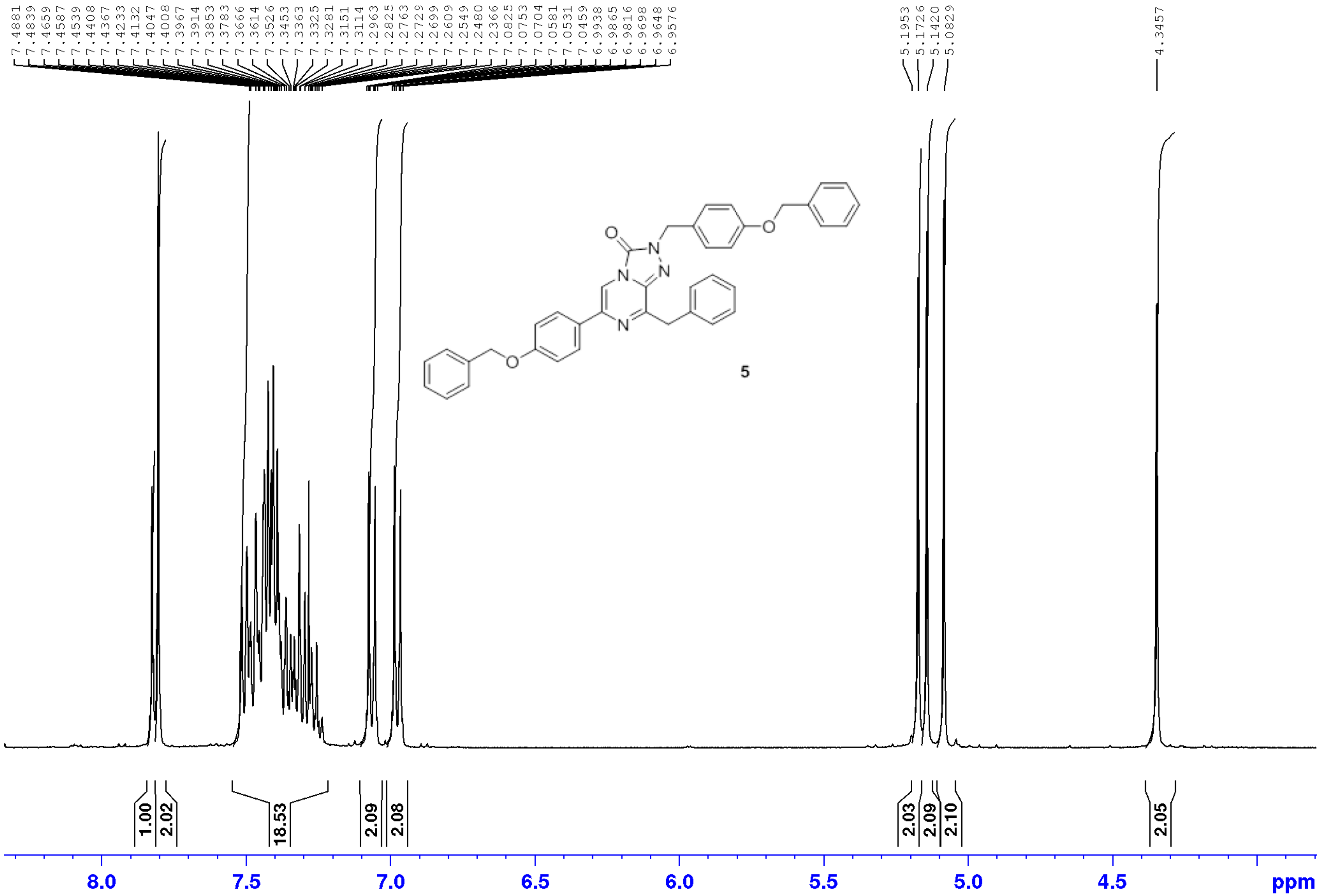

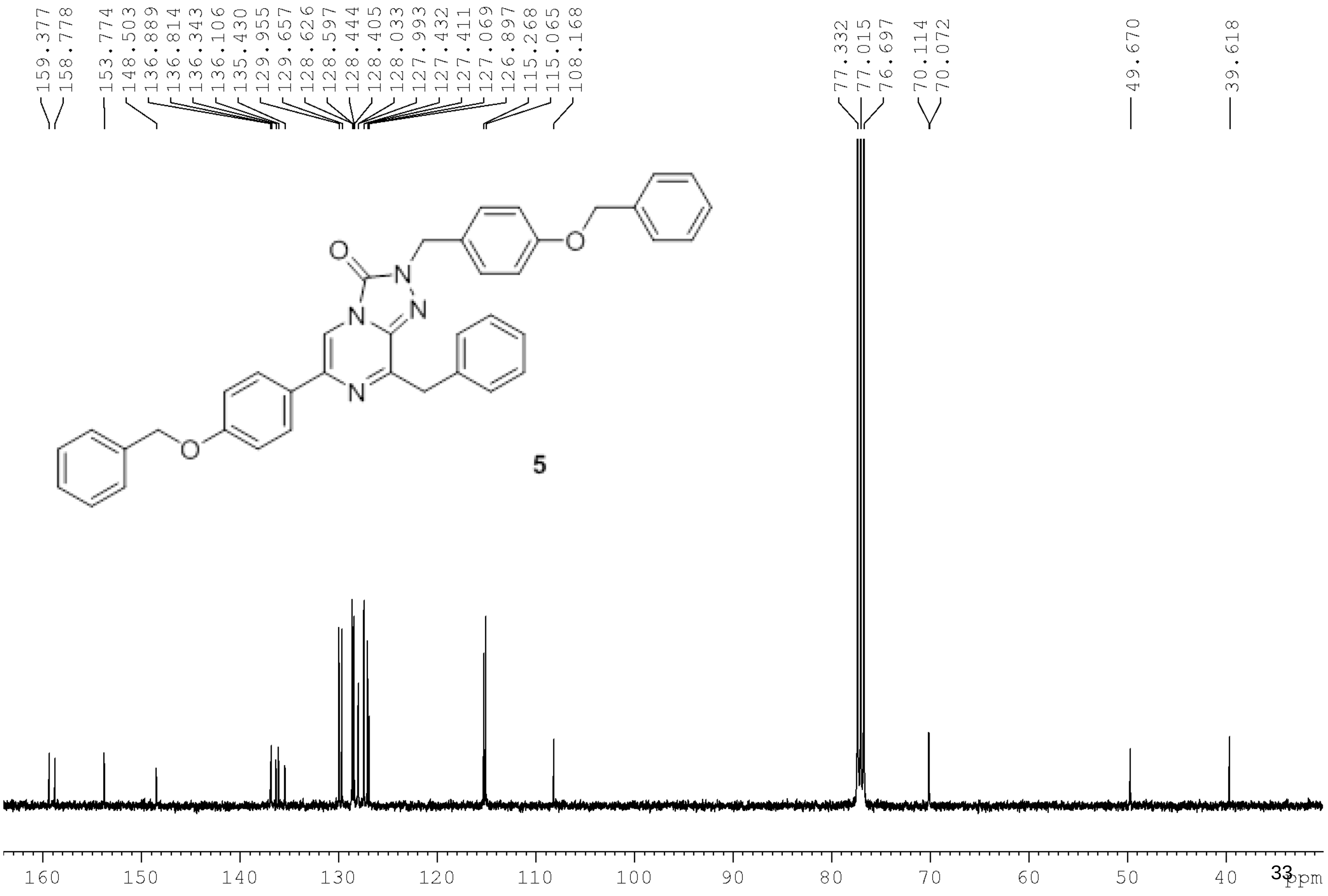
159.377
158.778
153.774
148.503
136.889
136.814
136.343
136.106
135.430
129.955
129.657
128.626
128.597
128.444
128.405
128.033
127.993
127.432
127.411
127.069
126.897
115.268
115.065
108.168
77.332
77.015
76.697
70.114
70.072
49.670
39.618
O
N
N
N
N
O
O
5
160
150
140
130
120
110
100
90
80
70
60
50
40
ppm

8-Benzyl-2-(4-hydroxybenzyl)-6-(4-hydroxyphenyl)-[1,2,4]triazolo[4,3-a]pyrazin-3(2H)-one (**azaCTZ**)

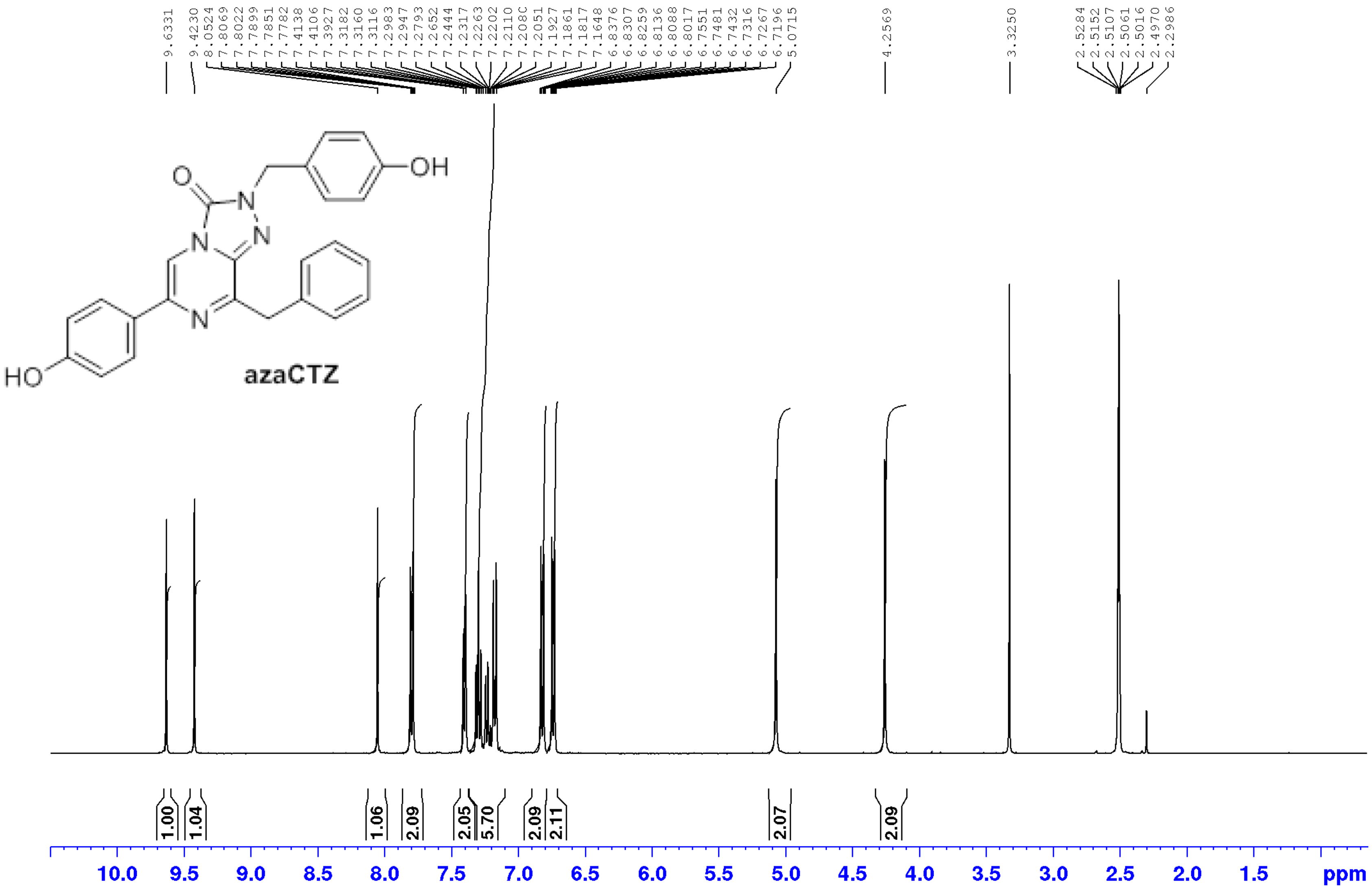

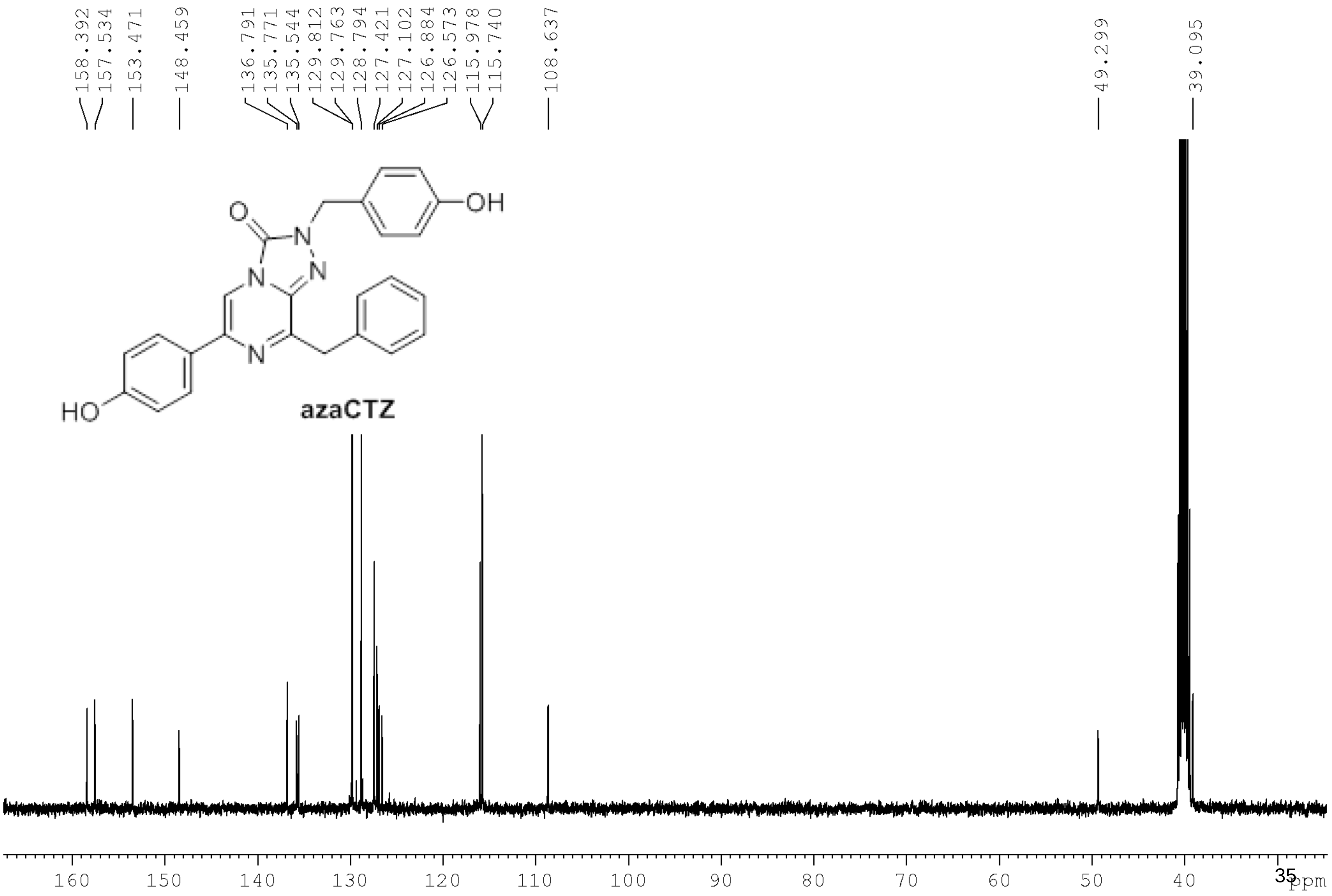
158.392
157.534
153.471
148.459
136.791
135.771
135.544
129.812
129.763
128.794
127.421
127.102
126.884
126.573
115.978
115.740
108.637
49.299
39.095
O
N
N
N
N
OH
HO
azaCTZ
160
150
140
130
120
110
100
90
80
70
60
50
40
ppm
